# Natural sensory context drives diverse brain-wide activity during *C. elegans* mating

**DOI:** 10.1101/2020.09.09.289454

**Authors:** Vladislav Susoy, Wesley Hung, Daniel Witvliet, Joshua E. Whitener, Min Wu, Brett J. Graham, Mei Zhen, Vivek Venkatachalam, Aravinthan D.T. Samuel

## Abstract

Natural goal-directed behaviors often involve complex sequences of many stimulus-triggered components. Understanding how brain circuits organize such behaviors requires mapping the interactions between an animal, its environment, and its nervous system. Here, we use continuous brain-wide neuronal imaging to study the full performance of mating by the *C. elegans* male. We show that as each mating unfolds in its own sequence of component behaviors, the brain operates similarly between instances of each component, but distinctly between different components. When the full sensory and behavioral context is taken into account, unique roles emerge for each neuron. Functional correlations between neurons are not fixed, but change with behavioral dynamics. From the contribution of individual neurons to circuits, our study shows how diverse brain-wide dynamics emerge from the integration of sensory perception and motor actions within their natural context.

## Introduction

Natural behaviors involve strong interactions between animals and their surroundings. When an animal is isolated from its natural context, brain dynamics do not display their full range of mechanisms (**1, 2**). For example, recent brain-wide imaging studies in immobilized *C. elegans* and other animals have uncovered strong activity correlations among large populations of neurons (**3, 4**). However, circuit activity becomes more complex when animals are allowed to move and display a larger variety of behaviors. Studies in freely-moving mice have uncovered changes in the functional correlations between brain areas when the animal is engaged in different behaviors (**5**). Freely-moving *C. elegans* hermaphrodites exhibit less correlated brain activity than immobilized worms (**6**). In the mouse, increasing task complexity and the richness of the sensory environment have been shown to progressively decorrelate neurons and brain regions (**7**). Task-dependent functional reorganization of neuronal circuits has been observed in several invertebrate, vertebrate, and artificial systems (**1, 8**–**11**). Here, we use the mating behavior of the male *C. elegans* to ask how the full richness of a natural behavior is embedded in brain structure and function.

Mating behavior is highly interactive and involves variable sequences of recurrent behavioral components or *motifs* that carry the animal towards its goal. The male *C. elegans* uses a dedicated circuit in his tail – a posterior brain with over 100 sensory neurons, interneurons, and motoneurons – to drive the many motifs of mating (**12**–**14**) (**Fig. 1A, B**). He searches for a hermaphrodite by detecting pheromones. When he contacts a hermaphrodite, he presses the ventral sensilla of his tail against her body. He then starts scanning along her body to locate the vulva. During scanning, he frequently changes his movement direction, executes sharp turns when moving around the anterior and posterior ends of her body, and slows at the vulva. Upon vulva location, he will attempt to insert his mating organs (spicules). Successful spicule insertion triggers sperm transfer. Copulation is followed by a prolonged rest (**15**–**18**). All of these behavioral motifs can only occur in the context of freely-moving males interacting with a hermaphrodite.

**Fig. 1.**
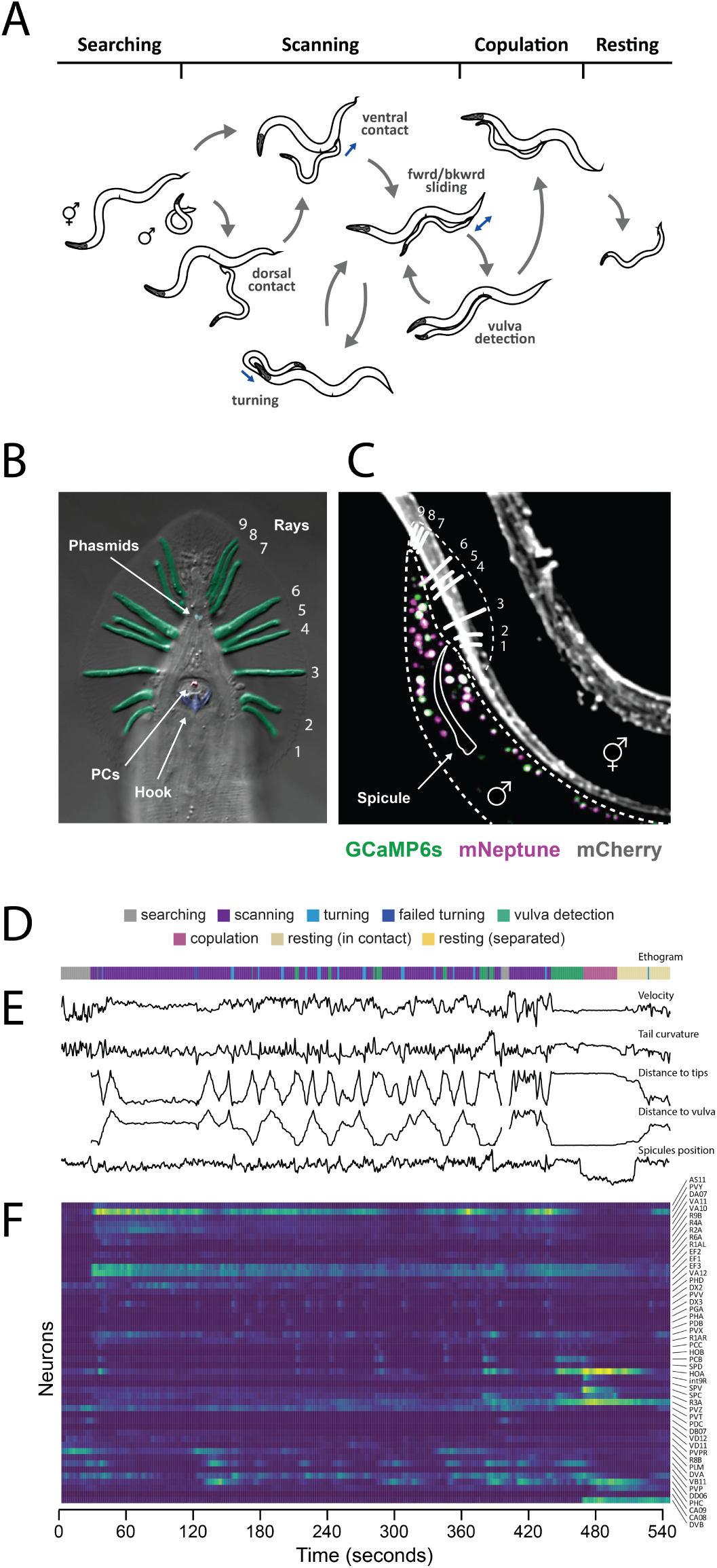
Mating in the male *C. elegans* is a multi-step natural behavior. (**A**) The male switches between several behavioral motifs using inputs from tail sensory organs including 9 pairs of sensory rays (pseudo-colored in green) innervated by two neurons each, the hook (blue), which harbors two sensory neurons, two postcloacal sensilla (pink) innervated by three neuron pairs, and two phasmids (aqua) innervated by two neuron pairs (**B**). (**C**) We used a tracking microscope to simultaneously record male behavior and neuron activity in the mating circuit using a strain expressing a nuclear-localized calcium indicator (GCaMP6s) and a red fluorescent marker (mNeptune). Fluorescent hermaphrodites were used to track male behavior. (**D**) An ethogram showing a behavioral trajectory for a single mating event. (**E**) Continuous behavioral features including male velocity, posture, and position relative to the partner were extracted from animal movements. (**F**) Throughout the mating event, the activity of all visible neurons in the tail was continuously recorded. Most neurons could be identified.

We recorded the activity of nearly every neuron in the posterior brain of freely-moving male *C. elegans* from beginning to end of mating. Brain-wide imaging revealed functional relationships among all neurons for every step of mating behavior. When brain-wide dynamics are considered in the context of an entire natural behavior, nearly every neuron exhibits a unique activity pattern. Each of the many different behavioral motifs during mating has a specific and consistent circuit representation. As the behavioral sequence unfolds, functional correlations between neurons in the brain are not fixed, but depend on context and change with the dynamic interactions between the animal, the nervous system, and the environment. For one wiring diagram to support many correlation patterns, the relationship between the connectome and activity cannot be one-to-one. By combining functional imaging, behavioral analysis, and single-neuron manipulations, we identify circuit-level mechanisms for several computations that are ethnologically relevant for mating. We show how brain-wide dynamics are shaped by the flow of sensory inputs and motor actions across the full performance of a natural behavior.

## Results

### Brain-wide imaging throughout mating

To record the activity of the male’s entire posterior brain, we used a spinning-disc confocal microscope customized for multi-color, multi-neuronal imaging (**19**). We used strains with pan-neuronal expression of both nuclear-localized calcium indicator (GCaMP6s) and red fluorescent protein (mNeptune) imaged at 10 volumes per second with single-cell resolution (**Fig. 1C**, Movie S1). Neuronal activity in each male was continuously imaged through every step of mating: initial search, contact, scanning, vulva detection, copulation, and rest.

For comprehensive analysis, we selected datasets of continuous 5-10 min recordings from seven animals that showed the broadest range of behavioral dynamics (**Table S1**). Most neurons in the mating circuit were recorded in multiple datasets. Each neuron was identified by its relative position, morphology, and expression of cell-specific fluorescent markers (**Table S2**). The seven selected datasets contained the activities of 76 identifiable neurons spanning ∼75% of the circuit (**Fig. S1A**). Nearly all pairs of left-right symmetric neurons were highly correlated (**Fig. S2**). Grouping left-right pairs as one neuron type, we recorded 57 neuron types (**Fig. S1B**). We also acquired additional datasets focused on specific neuron types or behavioral motifs.

We extracted behavioral features concurrent with circuit activity from video analysis of the movements and interactions of the male and hermaphrodite. We quantified a set of continuous features – velocity, sliding relative to the hermaphrodite, tail curvature, spicule position, and position of the tail along the hermaphrodite body – as well as discrete behavioral events and motifs (**Fig. 1D–F**).

### The collective activity of individual neurons predicts behavior across animals

Does brain-wide activity map to behavioral dynamics in the same way from animal to animal? If so, it should be possible to build a computational model that predicts one animal’s behavior based on the brain-wide activity of other animals. Because we could identify the same neurons in every dataset, we were able to build models that account for the individual contributions of each neuron, and test whether neuron-level models built using one group of animals could predict the behavior of other animals.

We built brain-to-behavior models for mating using sparse linear regression with short-term memory applied to the concatenated activity of 46 neurons that we recorded in nearly every experiment and a set of continuous behavioral features (**Fig. 2A, B**). In each iteration, we used six of our seven comprehensive datasets for concatenation and model training. We evaluated the resulting model in each iteration by its accuracy in predicting behavioral dynamics from brain-wide activity for the dataset that was left out (**Fig. 2A–C**). We assessed prediction accuracy across seven iterations. Sparse linear regression models accurately predicted most behavioral features, including tail velocity (mean *R*^2^ = 0.37), curvature (mean *R*^2^ = 0.32), sliding velocity (mean *R*^2^ = 0.27), distance of the tail to the hermaphrodite head or tail (mean *R*^2^ = 0.29) and the vulva (mean *R*^2^ = 0.31), and position of the spicules (mean *R*^2^ = 0.16) (**Fig. 2D**). Thus, these behavioral features are represented in the collective activity of individual neurons in a consistent manner across animals.

**Fig. 2.**
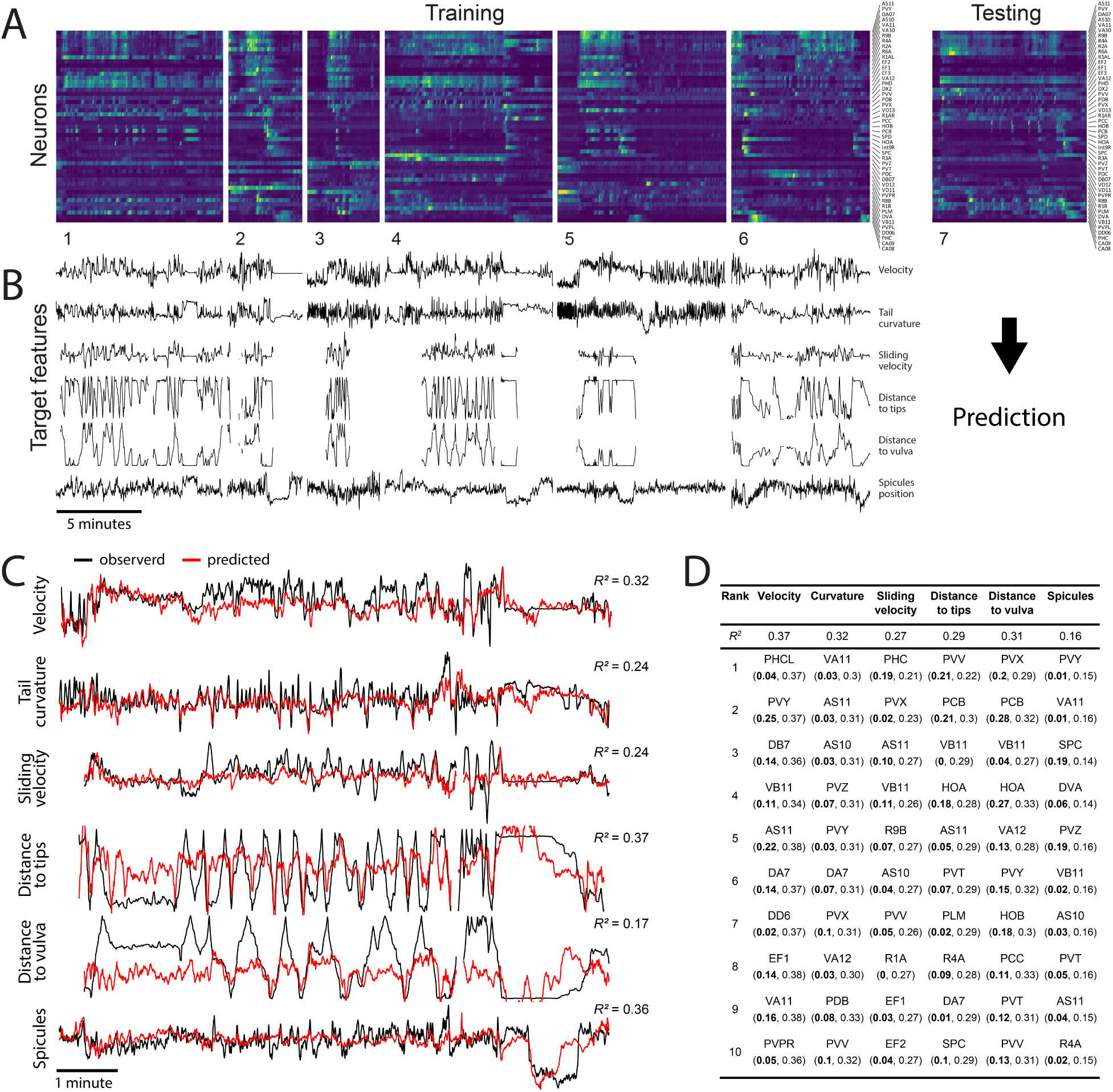
Continuous behavioral features can be decoded from neuronal activity across animals. (**A**) Concatenated activity traces of 46 neurons were used to train sparse linear models to predict several continuous behavioral features (**B**). (**C**) These models were able to predict male velocity, tail curvature, and male tail distance to the hermaphrodite tips and vulva. Observed and predicted behavioral features for one of seven datasets are shown. (**D**) Individual neurons ranked based on their importance for predicting specific behavioral features. Numbers in parenthesis show mean *R*^2^ for models built using traces of single neurons (in bold), and models built using traces of all except one neuron.

How distributed is the representation of behavioral features across neurons? Sparse linear regression ranks neurons by their relative contribution to each behavioral feature (**Fig. 2D**). In all cases, we found that excluding one top-ranking neuron from the datasets before training had only a small effect on prediction accuracy (**Fig. 2D**). Moreover, models trained using the activities of single top-ranking neurons performed worse than models trained using all neurons for most behavioral features (**Fig. 2D**). Thus, most behavioral dynamics during mating have distributed representations in the brain-wide activity.

### The brain partitions into functional communities for each behavioral motif

To better understand how neurons interact with one another during behavior (**Fig. 3A**), we computed neuron-neuron and neuron-behavior correlations for all pairs of neuronal activities and continuous behavioral features. We assembled a consensus correlation matrix that incorporated data from all experiments, and grouped neurons with similar activities using hierarchical clustering (**Fig. 3B**). Certain clusters were associated with distinct behavioral motifs such as scanning or vulva detection. These clusters indicated a group of principal neurons for each motif. We found that many neurons were strongly correlated with other neurons in more than one cluster, indicating that single neurons often participate in multiple behavioral motifs.

**Fig. 3.**
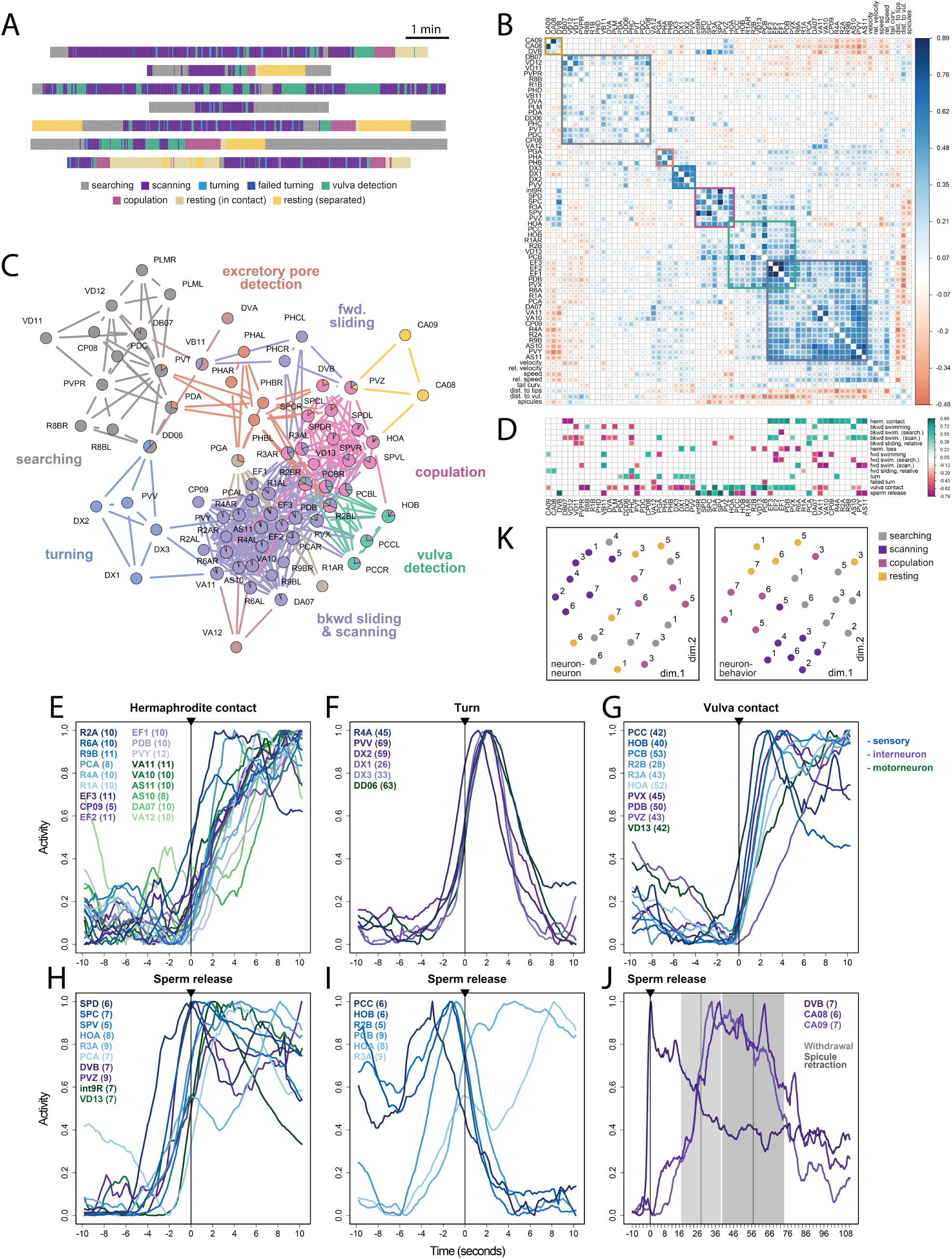
Functional organization of the mating circuit. (**A**) Each male exhibited unique behavioral dynamics represented by distinct ethograms. (**B**) Matrix of pairwise cross-correlations for 57 neuron types and 8 continuous behavioral features generated from the recordings of 18 males. Hierarchical clustering reveals groups of neurons with similar activities and associated with specific behavioral motifs. (**C**) A network of functional connections between neurons. Several neurons belong to multiple partly overlapping functional communities, shown with different colors. (**D**) Many neurons increase (green) or decrease (magenta) their activity at the onset of discrete behavioral events such as turning or stopping at the vulva. Neurons with significant activity changes are shown (FDR-corrected *p*-values < 0.05). (**E** to **J**) Activities of selected neurons from all males aligned to the onset of hermaphrodite contact, turning, vulva contact, and sperm release. The number of events observed for each neuron are in parenthesis. Colors indicate sensory, inter-, and motoneurons. (**I**) At the onset of copulation, vulva-tuned sensory neurons PCC, HOB, R2B, and PCB decrease their activity, although contact with the vulva remains. (**J**) Inter/motoneurons CA8, CA9, and DVB are active after copulation and during resting. Withdrawal from the vulva and spicule retraction events ± SD are shown. (**K**) Unsupervised arrangement of neuron-neuron and neuron-behavior correlation patterns for each animal and behavioral state. The correlations are more strongly grouped by behavioral state (distinguished by color) than by animal (distinguished by the dataset number)

To identify circuits for different motifs, we used *link clustering*, a community detection algorithm where each node in a network can belong to multiple communities (**20, 21**). Link clustering applied to the consensus matrix partitioned the mating circuit into overlapping communities for searching, scanning, turning, vulva detection, excretory pore detection, and copulation (**Fig. 3C**). Many neurons belonged to more than one community, most often between temporally adjacent motifs like scanning, vulva detection, and copulation.

### Unique activity patterns for each neuron

We compared the contributions of neurons to different behavioral motifs by aligning their activities to the onset of discrete events such as hermaphrodite contact and turning. Many neurons were consistently activated or inhibited by a different combination of behavioral events (**Fig. 3D**). When considered relative to the full set of behavioral events used for alignment, 80% of neurons exhibited unique activity patterns.

When we compared the activities of each group of neurons triggered by each behavioral event, we often observed diverse temporal dynamics (**Fig. 3E–J**). In many cases, we observed highly consistent sequences of peak activities among neurons from the start of a behavioral event, suggesting different and step-wise functional contributions for each neuron. Considering the activity of each neuron relative to all steps of mating behavior and at high temporal resolution, virtually every neuron in the mating circuit emerges as functionally unique.

### Functional correlations between neurons are specific to behavioral context

Not only do different and partly overlapping combinations of neurons contribute to different behavioral events, but the correlations exhibited by overlapping neurons can be different for different events. For example, the HOA and HOB hook sensory neurons have long been viewed as contributing similarly to detecting and stopping at the vulva. We confirmed that both HOA and HOB are co-active during vulva detection, but we discovered that HOA activity increases and HOB activity decreases during copulation (**Fig. 3D, G, I**). Thus, motif-specific correlations diversify brain-wide activity during mating.

To compare circuit-wide functional correlations across motifs, we partitioned each dataset into four states: searching, scanning, copulation, and post-copulatory rest. We calculated a separate correlation matrix for each state for each dataset (**Fig. S3, Fig. S4**). Neuron-neuron correlations as well as neuron-behavior correlations were more similar when we compared the same states exhibited by different animals than when we compared different states exhibited by the same animal. When arranged using multidimensional scaling, the correlation matrices were more strongly grouped by behavioral state than by animal (**Fig. 3K**).

### Synaptic interactions between functionally-connected neurons and functional communities

The wiring diagram of the male tail has been reconstructed by serial-section electron microscopy (**13, 14**). If this wiring diagram of chemical and electrical synapses supports the full diversity of brain-wide activity patterns across mating, functional connectivity will not be fully predicted by synaptic connectivity, or vice-versa. We estimated the functional connectivity between individual neurons using the correlation between their activity traces throughout mating (**Fig. 3B**). Activity correlations across all pairs varied from negative to strongly positive with a mean of 0.12 ± 0.22. Among pairs of neurons with positive correlations above the mean (> 0.12), 19% are connected by synapses. This probability is marginally higher than the 15% probability that two randomly sampled neurons are connected by synapses (**Fig. 4A**).

**Fig. 4.**
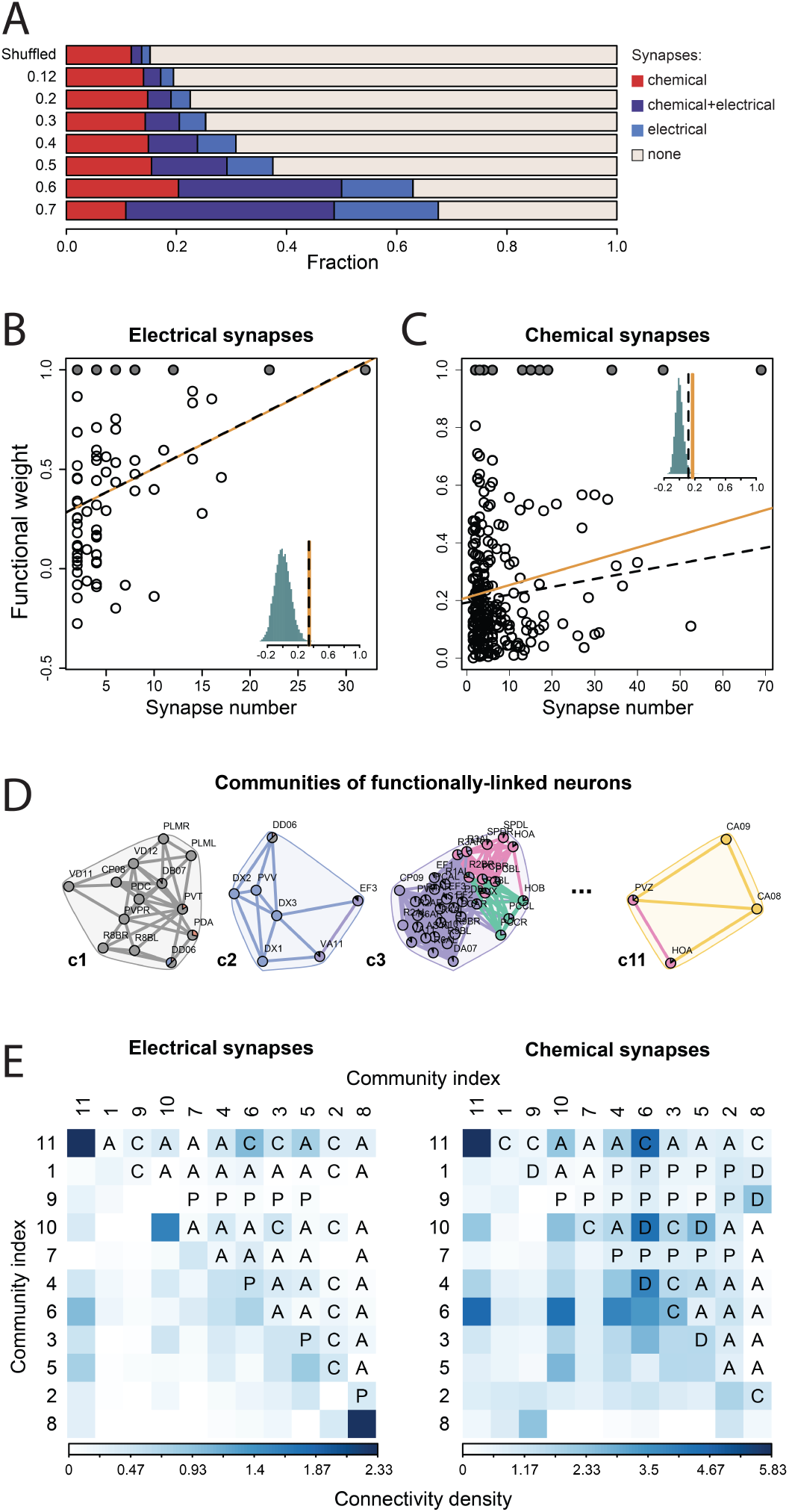
Mapping functional and synaptic connectivity. (**A**) Most functionally-linked neurons are not directly connected by synapses. Direct synapses, particularly electrical synapses, are more likely to occur between neurons that are more strongly functionally-linked. Functional weight as a function of the number of electrical (**B**) and chemical (**C**) synapses between neurons. Solid circles indicate imputed correlations between the left-right neuron pairs. Orange solid lines and black dashed lines indicate observed Pearson and partial correlations respectively and the histograms show the distribution of correlation coefficients when the functional weights are shuffled. (**D**) Overlapping communities of functionally-linked neurons extracted with link clustering. (**E**) In-group and out-group connectivity density for the functional communities. Assortative (“A”), disassortative (“D”) and core-periphery (“C”, “P”) community interactions involving electrical and chemical synaptic networks are shown.

Among neuron pairs with stronger positive functional correlations (> 0.3, > 0.5, and > 0.7), larger fractions were connected by synapses (25%, 38%, and 68%, respectively). When we considered more strongly correlated pairs, we observed larger fractions of connections with only electrical synapses or with both chemical and electrical synapses. In contrast, the fraction of connections with only chemical synapses did not change. For neurons connected by synapses, we also quantified the dependence of functional connectivity on the number of electrical or chemical synapses. Functional connectivity was positively correlated with the number of electrical synapses (Pearson correlation *r* = 0.35, *p* = 0.001, partial correlation *r* = 0.34, *p* = 0.0007), and less positively correlated with the number of chemical synapses (Pearson correlation *r* = 0.17, *p* < 0.0013, partial correlation *r* = 0.12, *p* = 0.021) (**Fig. 4B, C, Table S4**). Thus, functional connectivity between individual neurons is prevalent among neurons without synaptic connectivity. Among synaptically connected neurons, functional connectivity is better predicted by the number of shared electrical synapses than chemical synapses.

Functional correlations partition the mating circuit into overlapping communities for different behavioral motifs (**Fig. 3C, Fig. 4D**). How does synaptic connectivity contribute to signaling within and between the circuits for different steps of mating behavior? For each pair of functional communities, we calculated the density of synaptic connections within each community (*in-group connectivitiy*) and between the two communities (*out-group connectivitiy*). An interaction between two communities is called assortative when in-group connectivity for each community is stronger than the out-group connectivity between them (**22**). For electrical synapses, 61% of all pairwise interactions between functional communities were assortative. For chemical synapses, 40% of all pairwise interactions were assortative (**Fig. 4E**). Assortative interactions were rare when we randomly shuffled synaptic connectivity (18% ± 17). In contrast, disassortative interactions, when out-group connectivity between communities is stronger than each in-group connectivity, were prevalent when connectivity was randomly shuffled (51%). Disassortative pairwise interactions between functional communities were rare for both electrical (0%) and chemical synapses (12%) (**Fig. 4E**). In summary, both electrical and chemical connectivity play substantial roles in internal signaling within functional communities. Chemical synapses play the larger role in signaling between functional communities, perhaps coordinating the display of different behavioral motifs.

### Circuit-level implementation of behavioral motifs

The mechanisms that diversify brain-wide activity patterns and confer unique functional properties to each neuron are driven by stimulus, feedback, and motor patterns that only occur in the context of natural behavior. Here, we explore several of these mechanisms by dissecting circuits during distinct motifs of mating.

#### Scanning

The largest functional community within the mating circuit is associated with scanning along the hermaphrodite to locate the vulva. During scanning, the male recognizes the hermaphrodite, distinguishes different locations of her body, and uses feedback to keep track of his own movements.

#### Sensorimotor initiation of backward movement

Scanning begins with the initiation of backward movement when the tail makes ventral contact with a hermaphrodite. Earlier studies identified neurons required to initiate scanning including ray sensory neurons (**Fig. 1B**) and the PVY interneuron that receives synapses from the rays (**15, 23, 24**). Our brain-wide imaging revealed many more neurons for scanning involving 30% of the mating circuit from sensory neurons (R1AL, R2A, R4A, R6A, R9B, PCA) to interneurons (PVY, PDB, EF1-EF3, CP9) to motoneurons (AS10, AS11, VA10-VA12, DA7) (**Fig. 3E**, S1).

We found that many neurons are functionally correlated during scanning in an unfolding sequence of neuronal activity that triggers backward movement and suppresses forward movement upon hermaphrodite recognition. The first neurons to be activated when the male contacts a hermaphrodite are the R2A and R6A sensory neurons (**Fig. 3E**). The EF1-3 GABAergic interneurons, postsynaptic to R2A and R6A, become active shortly after contact. The activities of R2A, R6A, and EF1-3 are strongly correlated with each other and with backward scanning (**Fig. 3B**). The PVY interneuron also becomes active after contact, and is only active during backward movement (**Fig. 3D**, S1).

EF1-3 make GABAergic (presumably inhibitory) synapses to the AVB premotor interneuron in the head that drives forward locomotion (**25**). PVY makes excitatory synapses to the AVA premotor interneuron in the head, which is itself pre-synaptic to the A-type motoneurons that drive backward locomotion (**23, 26, 27**) and AS motoneurons. Indeed, the A-type motoneurons in the tail (VA10-VA12, DA7) and the AS10 and AS11 motoneurons become active with backward scanning. In this manner, hermaphrodite recognition triggers a switch from forward movement during search to backward movement during scanning through a synaptically-connected chain of neurons.

#### Sensory feedback during backward and forward sliding

The PCA postcloacal sensilla neuron is active during scanning. Close examination of its neuron-behavior correlations revealed a role in sensory feedback rather than in hermaphrodite recognition: PCA activity is strongly associated with the relative sliding between the male and hermaphrodite (*t*(4) = 3.53, *p* < 0.025), and is specifically active during backward sliding (**Fig. 3D**, S1).

Unlike other postcloacal sensilla neurons, PCA expresses the TRP-4 mechanosensory receptor and has striated rootlets like A-type sensory ray neurons (**12, 28**). PCA may detect backward sliding using mechanosensation. Unlike the ray sensory neurons, the synaptic outputs of PCA are to neurons for behavioral motifs other than scanning. Multiple synaptic targets of PCA are involved in vulva detection. PCA is correlated with other neurons involved in scanning, but its functional role is to monitor backward sliding to provide feedback to neurons in other motifs.

Scanning involves long backward movements that are interrupted by short forward movements. At the onset of forward movements, A-type motor neurons that drive backward movement are inactivated and B-type motor neurons that drive forward movement are activated (**Fig. 3D**, S1). Forward movements are often initiated when the male overshoots the vulva or the ends of the hermaphrodite (see below). Forward movement during scanning appears to be a form of sensory-triggered course correction.

The mating circuit monitors relative forward sliding between the male and hermaphrodite by mechanosensory feedback using the PHC sensory neuron, the sensory counterpart of PCA. Like PCA, PHC has striated rootlets for mechanosensation (**12, 29, 30**). Whereas PCA is correlated with backward sliding relative to the hermaphrodite, PHC is correlated with forward sliding relative to the hermaphrodite (**Fig. 3D**, S1).

PHC has sex-specific connectivity (13, 14, 31). In the male, one of the main synaptic targets of PHC is the PVY interneuron which drives backward movement through AVA. PHC is anticorrelated with PVY (**Fig. 3B**, S1). Thus, a feedback signal from PHC that detects forward sliding may be used to sustain forward sliding by inhibiting PVY. PHC also makes synapses to neurons involved in vulva detection. Thus, both PCA and PHC provide feedback about sliding to modulate other behavioral motifs.

#### Context-dependent posture control

The tail exhibits rhythmic undulatory movement when not in contact with a hermaphrodite. During mating, the tail stops rhythmic movement and assumes different prolonged postures with motif-specific purposes. Different locations on the hermaphrodite evoke the different sensorimotor transformations needed to assume these postures. These circuits control posture in different ways by context-dependent orchestration of the activity of overlapping sets of motoneurons and muscle cells (see below).

#### Ventral bending during turning

When the tail reaches either end of the hermaphrodite, the male must turn around her body with a sharp prolonged ventral bending of the tail to maintain contact and continue scanning (**Fig. 5A**). When turning is successful, the male continues scanning on her other side. When unsuccessful – for example, when the male overshoots before turning – the tail loses contact and the male switches to forward swimming, presumably trying to reestablish contact in a form of course correction.

**Fig. 5.**
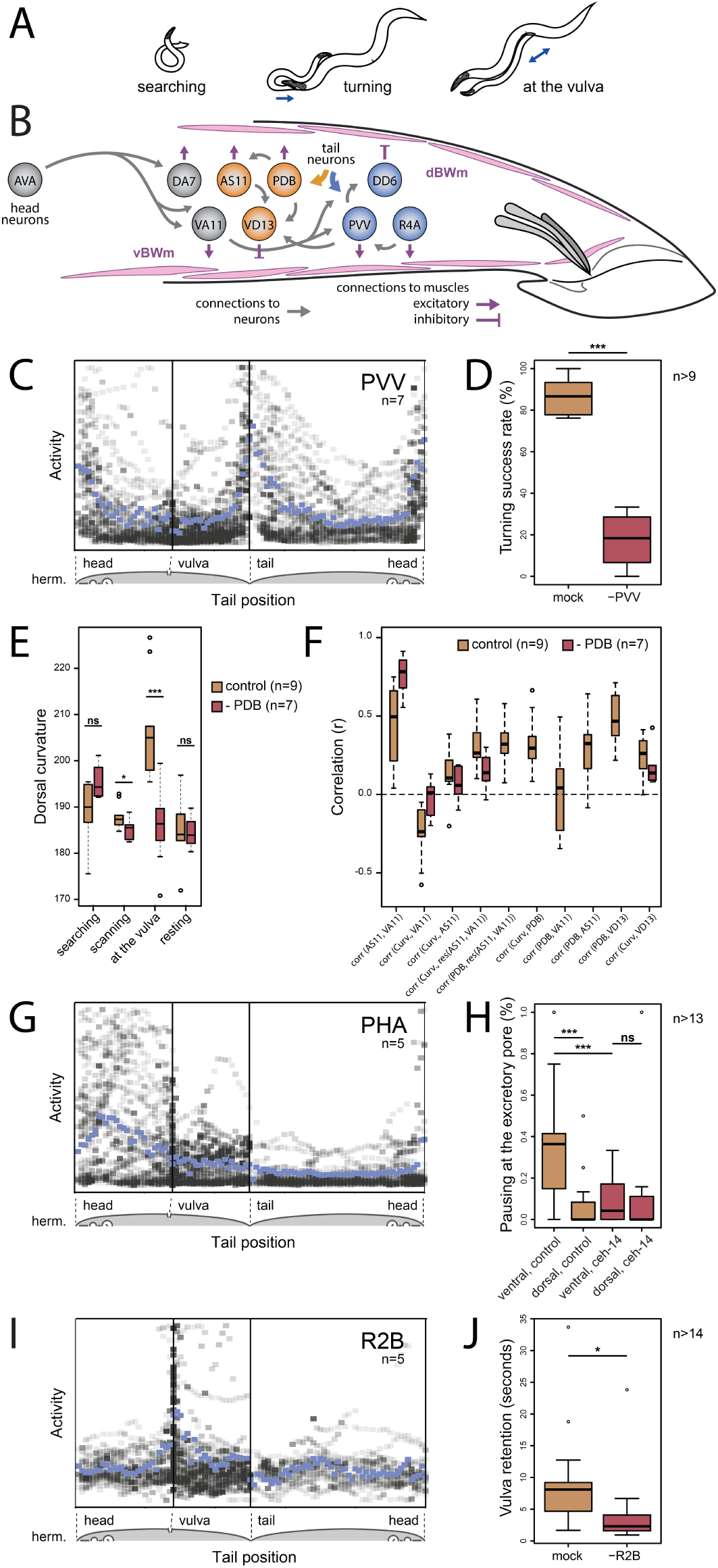
Circuit dissection of stimulus-triggered behavioral motifs. (**A**) The male adjusts his tail posture to keep in contact with the hermaphrodite using motif-specific circuits (**B**). Neurons involved in PVV-mediated turning are shown in blue and neurons involved in PDB-mediated curvature control during scanning are shown in orange. (**C**) The PVV interneuron becomes active when the tail flexes to turn around the ends of the hermaphrodite body. PVV activities from seven males relative to tail position on the hermaphrodite reveal high specificity of PVV activation. Mean activity is shown in blue. (**D**) PVV ablation compromises turning. (**E**) PDB-ablated males show less dorsal curvature when in contact with the hermaphrodite, particularly at the vulva. (**F**) Correlation coefficients between curvature and neuronal activity for 9 control and 7 PDB-ablated males. (**G**) Chemosensory PHA activates when the tail is near the hermaphrodite excretory pore. (**H**) Control males discriminate between the ventral and dorsal sides of the vulvaless hermaphrodites and pause near the excretory pore. *ceh-14* mutants, in which PHA and PHB fail to develop normally, do not pause at the excretory pore and move past without interruption. (**I**) The ray sensory neuron R2B is activated upon vulva contact. (**J**) Males in which R2B is ablated often lose contact with the vulva. *p<0.05, **p<0.01, ***p<0.001, ns – not significant

Several neurons become active specifically during turns including the R4A sensory neuron, the PVV interneuron, the DX1, DX2, and DX3 inter/motoneurons, and the DD6 motoneuron (**Fig. 3B, C, F, Fig. 5B, C**). Some neurons, like PVV, are active during both successful and unsuccessful turns, suggesting roles in turn initiation. Other neurons, like R4A and DD6, are only active after successful turns, suggesting roles in turn completion (S1).

PVV ablation severely disrupted turning behavior. Without PVV, males were usually unable to turn successfully at the ends of the hermaphrodite and often lost contact (*t*(16.43) = 14.44, *p* < 0.001) (**Fig. 5D**). PVV receives inputs from ray sensory neurons and has direct synaptic output to the ventral cord motoneurons and ventral body wall muscles (**13, 14**). PVV operates as a trigger for turning behavior by collecting sensory cues associated with the tapering ends of the hermaphrodite and directly signaling to neurons and muscles for ventral bending.

The R4A sensory neuron and DD6 motoneuron reach their peak activity when the male reaches the other side of the hermaphrodite. R4A directly innervates PVV and ventral body wall muscles. DD6 is a GABAergic motoneuron that relaxes dorsal body wall muscles. Thus, R4A and DD6 may prolong the ventral bending that allows the male to reach the other side of the hermaphrodite.

#### Dorsal bending near the vulva

The tail often makes a sharp dorsal bend when it reaches the vulva, presumably to help transition to copulation (**32**) (**Fig. 5A**). A subset of neurons are associated with dorsal bending, including sensory neurons involved in vulva detection, the PDB inter/motor neuron, and the AS11 and DA7 motoneurons (**Fig. 2D, Fig. 3B**).

PDB receives inputs from many different neurons including CP7-9, HOA, EF1-3, and ray neurons, and is thus sensitive to different context-dependent signals for regulating tail posture. PDB-ablated males showed no difference from normal males in dorsal bending during searching and resting (**Fig. 5E**). However, PDB-ablated males showed less dorsal bending during scanning and vulva detection (*t*(13.94) = 2.18, *p* = 0.046 and *t*(13.98) = 4.14, *p* = 0.001) (**Fig. 5E**). PDB may directly evoke dorsal bending through its synaptic innervation of dorsal body wall muscles, as well as by activating AS11 (which synapses onto dorsal muscles) and VD13 (which relaxes ventral muscles) (**27**) (**Fig. 5B**). Thus, PDB is poised to change the functional correlations between motoneurons when driving dorsal bending in a context-dependent manner.

The AS11 motoneuron is positively correlated with backward scanning, consistent with its receiving 30% of its synaptic input from the AVA premotor interneuron. AS11 is thus positively correlated with A-type motoneurons for backward movement including VA11 which activates ventral muscles (**Fig. 3B, D, Fig. 5B, F**). Although both AS11 and VA11 are correlated with backward movement, they have contrasting relationships to tail curvature. AS11 is positively correlated with dorsal bending whereas VA11 is negatively correlated with dorsal bending (*r* = 0.12 ± 0.15 and *r* = −0.13 ± 0.15). In PDB-ablated males, AS11 and VA11 became more correlated with each other and less correlated with curvature. Thus, PDB appears to contribute to the curvature-dependent portion of AS11 activity (**Fig. 5F**).

AS11 receives 15% of its synapses from PDB. Because PDB is correlated with dorsal curvature (*r* = 0.29±0.19) (**Fig. 5F**, S1) but not velocity (*r* = −0.03±0.23), PDB contributes the curvature-dependent portion of AS11 activity but not its direction-dependent portion. PDB has a similar effect on the VD13 motoneuron, which relaxes ventral muscles. During scanning, VD13 is correlated with PDB and with curvature. PDB ablation lessens the correlation between VD13 and curvature (**Fig. 5F**).

Both PDB and PVV provide mechanisms for context-dependent posture control. These interneurons collect sensory input that provide spatial cues about the vulva and hermaphrodite ends, and target overlapping sets of motoneurons to control dorsal and ventral bending for different purposes.

#### Excretory pore detection

When the tail reaches the hermaphrodite’s excretory pore while scanning, the PHA and PHB phasmid sensory neurons become active (**Fig. 5G, Fig. S5A**, S1). The phasmid neurons have sex-specific synaptic connectivity (**13, 14, 33**). The hermaphrodite uses her phasmid neurons for chemotactic responses (**34**). The male might use his phasmid neurons to detect excretory pore secretions to prevent mistaking the pore for the vulva (**35**) or as a spatial cue to guide him to the vulva.

The excretory pore is anterior to the vulva on the ventral side of the hermaphrodite. If the excretory pore is a guidance cue, its detection should affect movement decisions on the ventral side of the hermaphrodite. Because the vulva is the dominant feature that affects movement decisions on the ventral side, we examined male behavior with vulvaless hermaphrodites (*let-23* mutants, **Fig. S5B, C**). During scanning with vulvaless hermaphrodites, males pause during 36% of scans on the anterior ventral side but only 8% of scans on the anterior dorsal side (*t*(28.26) = 3.67, *p* < 0.001) (**Fig. 5H**). Pauses near the excretory pore are sometimes followed by forward slides that returns the tail to where the vulva should be. Thus, the excretory pore provides a guidance cue for course correction when the male overshoots the vulva.

Mutant *ceh-14* males have defects in phasmid neuron development (**36, 37**). These males do not respond to the excretory pore, pausing in only 9% of backward scans on either side of the hermaphrodite (*t*(24.77) = 0.03, *p* = 0.97) (**Fig. 5H**). The phasmid neurons thus likely contribute to triggering pauses. The PGA interneuron is also specifically active when the tail is near the excretory pore, and is thus functionally correlated with the phasmid neurons (**Fig. 3B, C**, S1). The wiring diagram does not have direct synaptic connections from the phasmid neurons to PGA, but the circuit for excretory pore detection extends from sensory neurons to interneurons.

#### Vulva recognition

When the male reaches the vulva, he stops scanning and transitions to another behavioral motif, vulva recognition. During vulva recognition, the male actively reduces his relative movement with respect to the hermaphrodite while pushing his spicules toward the vulva in repeated thrusts. Successfully breaching the vulva leads to copulation. Often, however, the male is unable to insert spicules and loses contact with the vulva. When this happens, he transitions to scanning. Mating can involve many transitions between scanning and vulva recognition.

The PCB and PCC postcloacal sensilla neurons and the HOA and HOB hook neurons are strongly associated with detecting the vulva (**Fig. 1B, D**). These neurons were previously shown via ablation experiments to help stopping at the vulva (**15**). Our brain-wide imaging revealed many additional neurons for vulva recognition including the R2B sensory neuron, the PVZ and PVX interneurons, the PDB inter/motoneuron, and the VD13 motoneuron (**Fig. 3B–D, G, Fig. 5I**). As discussed above, PDB and VD13 contribute to dorsal bending near the vulva. When the tail is aligned with the vulva for copulation, the second ray pair is closest to the vulva suggesting that R2B might be specialized to detect the vulva and trigger the transition to vulva recognition behavior (**Fig. S6**). Indeed, R2B ablation reduced the likelihood and duration of stopping at the vulva (*t*(24.29) = 2.54, *p* = 0.02, *t*(21.61) = 2.2, *p* = 0.04) (**Fig. 5J**).

#### HOA mediates a behavioral switch between scanning and vulva recognition

Previous ablation studies suggested that HOA contributes to both scanning and vulva recognition but in different ways. Near the vulva, HOA contributes to stopping and spicule insertion. Away from the vulva, HOA is thought to suppress other vulva-sensing neurons because its ablation causes the male to spuriously pause anywhere on the hermaphrodite and initiate spicule thrusts (**15, 38**). To determine how HOA contributes to switching between scanning and vulva recognition, we combined cell-specific neuron ablation with brain-wide imaging (**Table S3**).

In control animals, a group of vulva-detecting neurons – PCB, PCC, R2B, and PVX – is specifically active when the tail is at the vulva (**Fig. 6A, B**). In HOA-ablated males, these neurons exhibited frequent bursts of correlated activity. These bursts coincided with spurious pauses and spicule thrusts away from the vulva (**Fig. 6C, Fig. S7**). Spurious pauses interfered with reaching the vulva. When HOA-ablated animals did reach the vulva, they were often unable to maintain contact. During scanning, HOA is needed to inhibit other vulva-detecting neurons. During vulva recognition, HOA is needed to maintain contact and initiate copulation. How does HOA enable distinct functional correlations within the mating circuit during different behavioral motifs?

**Fig. 6.**
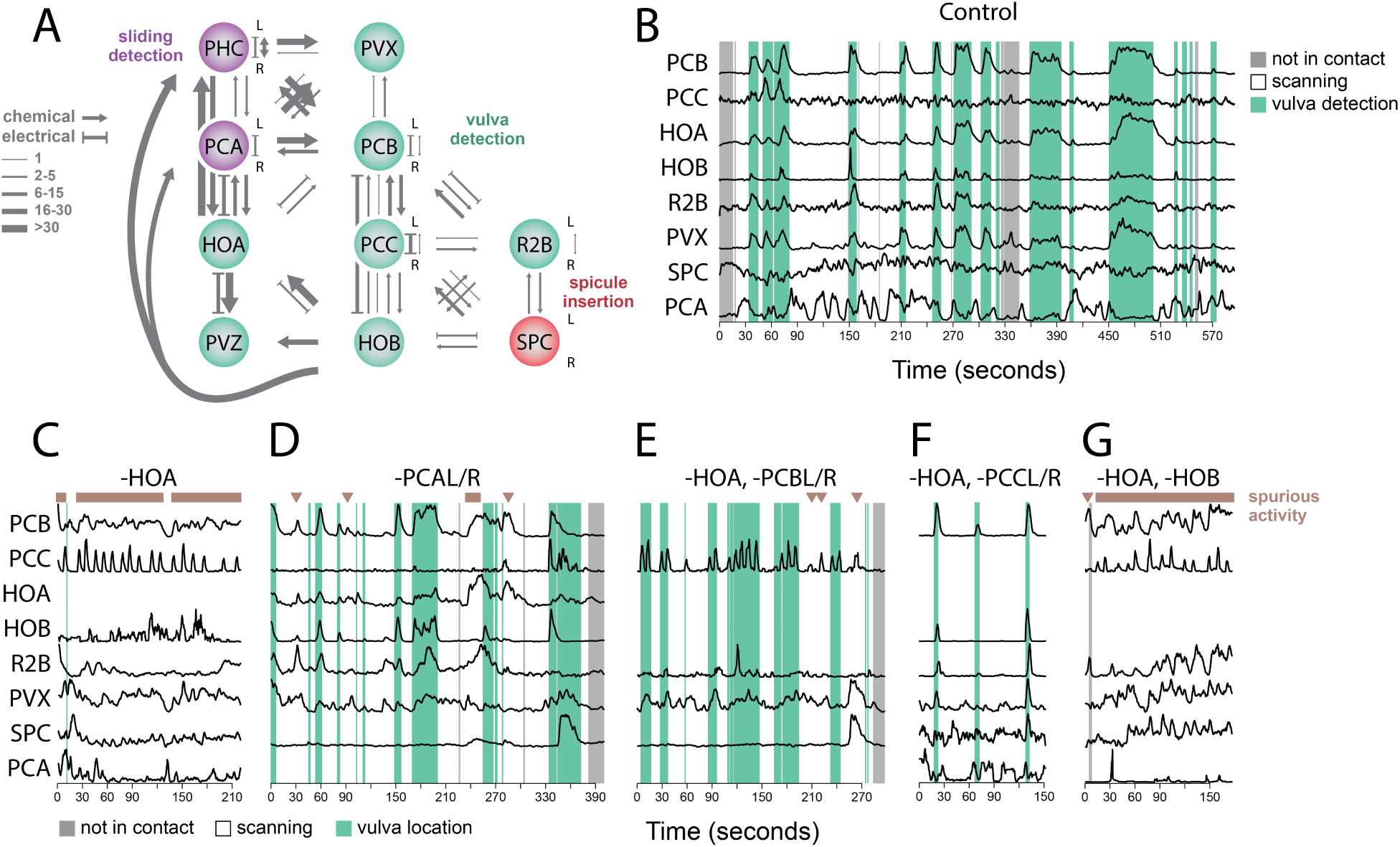
Signal amplification and inhibition within and across circuits control switching between behavioral motifs. (**A**) Vulva detection involves a circuit with mostly sensory neurons that are recurrently connected by electrical and chemical synapses (shown in green). (**B**) In the unoperated males, PCB, PCC, HOA, HOB, PVX, and R2B are correlated and their activation occurs when the tail is at the vulva. PCA shows the opposite pattern and is active during backward scanning. (**C**) Ablation of HOA leads to spurious activation of neurons involved in vulva detection. Repeated bursts of activation of the vulva-detecting circuit coincide with the spicule insertion attempts away from the vulva compromising scanning and reaching the vulva itself. (**D**) Ablation of PCA leads to spurious activation of the vulva-tuned neurons, but less frequently than HOA-ablated males. (**E** to **G**) Double ablations of HOA and PCB and PCC but not HOB attenuate the HOA ablation phenotype and revert scanning and vulva detecting behavior to nearly normal.

HOA makes both electrical and chemical synapses to the PCB vulva-detecting neuron (**Fig. 6A**). Thus, both hyperpolarization or depolarization of HOA may propagate to PCB and hence to other other vulva-detecting neurons. Glutamatergic inhibition from PCA has been shown to hyperpolarize HOA (**38**). We found that the PCA is active during backward scanning away from the vulva. PCA may thus hyperpolarize HOA during backward scanning. Near the vulva, HOA is directly activated by sensory cues from the vulva. The mating circuit thus contains mechanisms by which HOA excites other vulva-detecting neurons near the vulva and inhibits the same neurons away from the vulva.

To test whether PCA is part of a context-dependent switch that regulates HOA, we examined PCA-ablated animals. The phenotypes caused by HOA and PCA ablations were qualitatively similar but different in magnitude. Like HOA-ablated animals, PCA-ablated animals exhibited spurious activity in the neurons for vulva-recognition (**Fig. 6D, Fig. S7**). This spurious activity was more common with HOA-ablation than with PCA-ablation, suggesting that other neurons might also inhibit HOA during scanning. The PHC sensory neuron has the second strongest synaptic input to HOA (**Fig. 6A**). Like PCA, PHC is glutamatergic but is active during forward movement.

We conclude that HOA is both a positive and a negative regulator of the vulva-detecting circuit. HOA changes its effect on synaptic partners in a context-dependent manner. Near the vulva, HOA is activated by vulva detection, and further activates other vulva-detecting neurons. During scanning, PCA and PHC provide glutamatergic inhibition to HOA, which leads to hyper-polarization of other vulva-detecting neurons through electrical synapses.

#### Recurrently-connected sensory neurons and a self-reinforced activity state

After HOA ablation, the vulva-detecting sensory neurons – PCB, PCC, HOB, and R2B – exhibited correlated bursts of activity during spurious pauses and spicule thrusts. Because this activity occurs away from the vulva, it might be triggered by the hermaphrodite cuticle itself. One possibility is that the cuticle fully activates one or more of the vulva-detecting neurons away from the vulva. Alternatively, the cuticle might not fully activate one neuron, but, when some neurons become activated at a low level, recurrent excitation amplifies sub-threshold signals into a burst. The wiring diagram reveals many synapses initiate defecation (**38, 40**). In the male, defecation and ejacula-between vulva-detecting neurons that might allow recurrent excitation (**13, 14**) (**Fig. 6A**).

To test whether recurrent excitation occurs, we released the vulva-detecting circuit from inhibition away from the vulva by ablating HOA. If recurrent excitation plays a role, removing additional neurons should reduce the spurious activation of others. We found that additionally ablating either PCB or PCC reduced spurious activation of the rest of the vulva-detecting circuit (**Fig. 6E, F, Fig. S7**) – remaining sensory neurons were less activated by the cuticle, but were activated by the vulva. However, additionally ablating HOB did not suppress the spurious activity of remaining neurons (**Fig. 6G, Fig. S7**). PCB and PCC appear to contribute to recurrent excitation, but not HOB.

Recurrent excitation and signal amplification may facilitate the all-or-none switch of circuit activity from scanning to vulva-recognition. To confine its activity to the vulva, the vulva-detecting circuit must be under strong context-dependent inhibition. Integrated mechanisms for recurrent excitation and context-dependent inhibition enable sensitive, robust, and abrupt switching between temporally-adjacent behaviors.

#### Copulation

Breaching the vulva leads to the transition from vulva recognition to copulation behavior. Copulation involves full spicule insertion, sperm release, and withdrawal. Previous experiments showed that several sensory and motoneurons contribute to the different steps of copulation (**15, 18, 39, 40**). Here, we directly visualized the temporal sequence in neuronal activity during copulation (**Fig. 3H–J**). Several mechanisms, including chemosensory and mechanosensory feedback, shape the functional correlations and temporal progression of neuronal activities during copulation.

The first neuron involved in copulation, SPD, is activated before full spicule insertion. SPD has sensory endings at the spicule tips. The next neurons to be activated are the SPC sensory/motoneuron and DVB inter/motoneuron. SPC sends sensory endings to the base of the spicules, suggesting a role in detecting spicule movement. SPC is also a motoneuron that innervates the spicule protractor muscles and the gonad. The spicule protractor muscles are innervated by both SPC and DVB. Both SPC and DVB exhibit a tightly-correlated spike-like activation upon spicule protraction (**Fig. 3H, J**), and are required for spicule insertion (**15, 39**).

Following spicule protraction, the next neuron to be activated is SPV, another sensory neuron with endings at the spicule tips. The timing of SPV activation is consistent with its proposed roles in detecting the internal environment of the uterus and controlling sperm release. SPV ablation interferes with the precise timing of sperm release (**15, 18**).

HOA is active during vulva detection, but exhibits peak activity shortly after spicule insertion. The PVZ interneuron has similar activity to HOA but with slower dynamics, active during vulva detection and peaking after spicule insertion. The wiring diagram predicts multiple synapses between HOA and PVZ (**13, 14**).

Approximately two seconds after spicule insertion, calcium transients occur in multiple intestinal cells including Int9R. Anterograde calcium transients across intestinal cells are known to initiate defecation (39, 41). In the male, defecation and ejaculation motor sequences have shared components and circuitry (**39, 41**). We hypothesize that intestinal calcium transients might have dual roles in initiating ejaculation and defecation in the male (but see (**39**)). The PCA postcloacal sensilla neuron becomes active following these intestinal calcium transients. The exact role of PCA in copulation is unclear, but it might be involved in the movements of the gubernaculum, a sclerotized structure involved in mating. PCA directly innervates the gubernaculum erector and retractor muscles (**13, 14, 18**).

The activities of the CA8 and CA9 inter/motoneurons start increasing several seconds after spicule insertion, peak roughly when the male detaches from the vulva, and decrease with spicule retraction (**Fig. 3J**). This activity pattern is consistent with the proposed role of CA neurons in initiation and continuation of sperm transfer (**42**).

Four vulva-detecting sensory neurons – PCB, PCC, HOB, and R2B – decrease their activity upon sperm release even when maintaining contact with the vulva (**Fig. 3I**). As the male transitions to ejaculation, several neurons that are involved in an earlier step in the temporal sequence are deactivated by the circuit. These vulva-detecting neurons might interfere with later steps in copulation requiring their deactivation. The pathway for this deactivation might involve dopamine release from dopaminergic socket cells, which has been implicated in the switch between pre- and postejaculation behaviors (**18**).

In summary, brain-wide imaging has revealed sequential patterns of circuit activity that are driven by sensory inputs and feedback along the step-wise progression of copulation behavior.

#### Resting

After copulation, the male rests for several minutes, during which his motor activity is suppressed (**18**). Resting coincides with decreased global activity of the mating circuit (**Fig. S8**). Few neurons are active during resting, but these have distinct activity patterns. DVB, which becomes active when copulation begins, is the only neuron with sustained activity throughout rest (**Fig. 3J**). The PVZ and CA8-9 interneurons, which become active later during copulation, remain active until spicule retraction.

#### Searching

After resting, the male searches for another partner. Most neurons previously associated with searching are in the head (**16, 18, 43, 44**). We discovered that several neurons in the posterior brain are also associated with searching (**Fig. 3C**). Almost all ray sensory neurons are associated with motifs involving physical contact with the hermaphrodite. However, one pair of ray neurons, R8B, shows higher activity without contact. Several interneurons – PDA, PVPR, and PVT – are also active during searching, as well as several ventral cord motoneurons – DB7, VD11, and VD12 – that drive forward swimming (**27**).

Interestingly, the PLM mechanosensory neuron is more active during searching than during contact with a hermaphrodite. PLM is ordinarily required to initiate forward movement in an escape response when either males or hermaphrodites are touched at the posterior (**45**). We found that PLM activity drops sharply when scanning begins. PLM activity does not immediately rise when the male loses contact. PLM may be suppressed when the male contacts a hermaphrodite, allowing mating behavior to be sustained by inactivating the circuit for escape response behavior.

## Discussion

Natural multi-step behaviors challenge the brain to perform diverse tasks across the range of what the brain evolved to do (**2, 46**). Here, we analyze, with single-neuron resolution, how the entire posterior brain drives the full performance of mating by the male *C. elegans*.

Each mating between a male and hermaphrodite unfolds differently because of the complex and dynamic interactions of two animals. The sequence, timing, and detailed performance of different steps of mating differ for each trial. However, continuous brain-wide recordings reveal that the neuronal activities associated with the many recurring steps of mating are strikingly consistent. The brain operates in the same way between instances of the same component behavior, whether recurring in one animal or occurring in different animals. Neuronal activity that is specific to each step of behavior appears at all levels of signal processing from sensory to inter-to motoneurons.

When the full behavior is taken into account, we find little redundancy in the activities of different neurons. Nearly every neuron has a different set of correlations with respect to the rest of the circuit and to behavioral dynamics. Some neurons are specialized for single behavioral motifs. Other neurons participate in multiple motifs. Each motif engages a unique set of neurons, but different motifs often engage overlapping neurons. During the execution of each motif, neurons may be activated in a temporal sequence, revealing step-wise functional contributions as behavior unfolds. When brain-wide activity is considered in the context of an entire natural behavior, nearly every neuron exhibits a unique activity pattern.

Functional correlations between neurons in the brain are not fixed, but change with shifting contexts as a natural behavior unfolds. These functional correlations are produced by the dynamic interactions between the animal, the nervous system, and the environment. A prominent mechanism that produces context-dependent functional correlations is the stimulus-evoked recruitment of sensorimotor pathways. The hermaphrodite presents different stimulus patterns to the male with different parts of her body and at different steps during mating. These stimulus patterns create context-dependent correlations among the sensory neurons of the posterior brain. The diversity of these correlations underlies the diversity of sensorimotor responses in an entire behavior (**Fig. 7**).

**Fig. 7.**
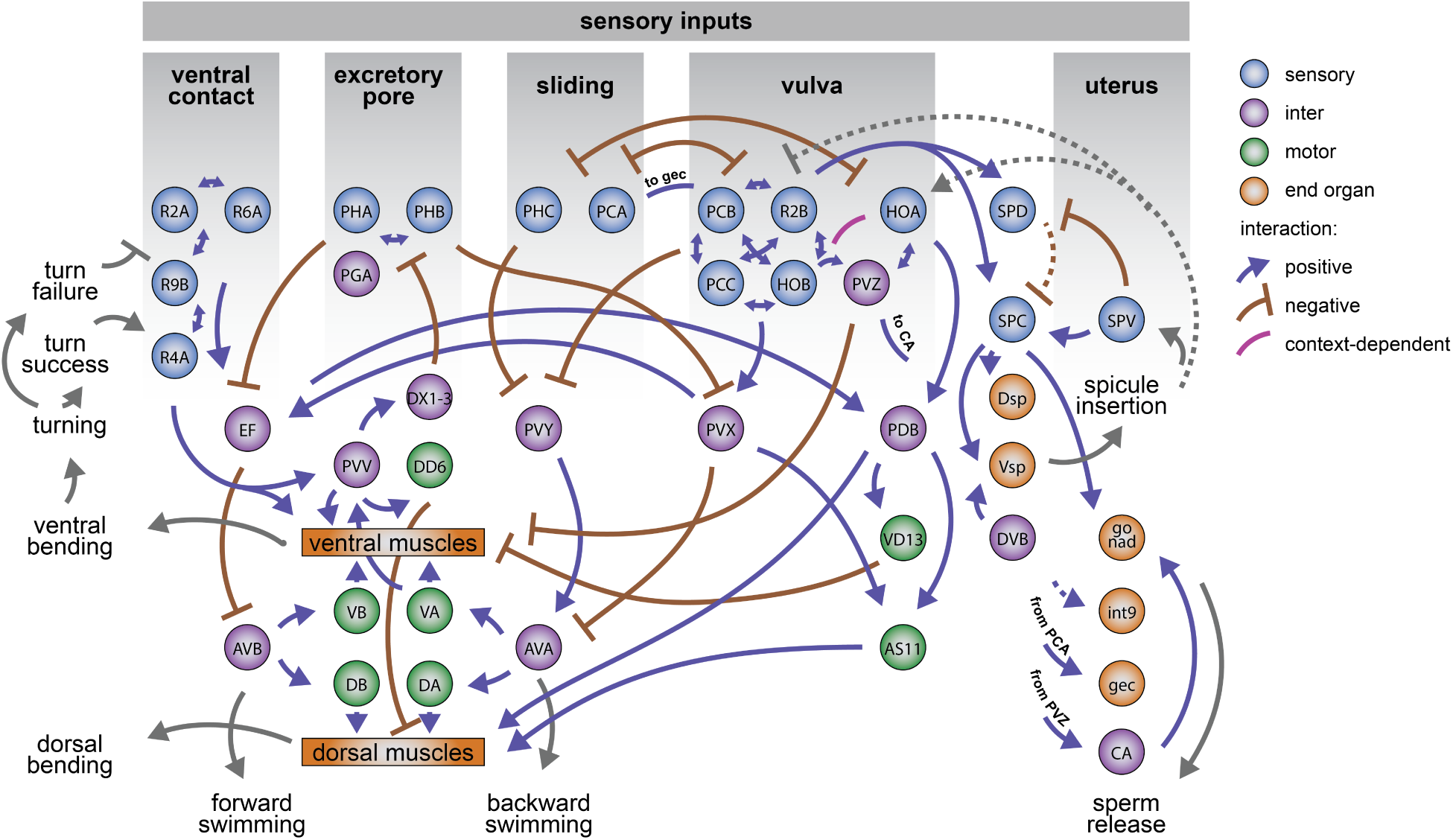
Conceptual diagram of stimulus-triggered information flow in the posterior brain. Distinct sensory patterns from the hermaphrodite at different steps of mating are used by the male to execute different motor actions and behavioral motifs. Stimulus patterns from the hermaphrodite act on diverse and specialized sensory neurons. Sensory perception requires multisensory integration in the brain that is carried out by direct interactions between sensory neurons and by downstream circuits. Multiple circuits use overlapping sets of interneurons to effect the decision to move forward vs. backward. Circuits for motif-specific posture control use a common set of motor neurons and muscles. Circuits for different motifs interact with one another to control behavioral transitions. Sensory feedback is used to keep track of behavioral dynamics and the steps of mating behavior, enabling error correction and the proper sequence of behavioral events. The diagram illustrates known and hypothetical positive, negative, and context-dependent interactions. Dashed lines indicate hypothetical interactions with undetermined synaptic mechanisms.

The richness of natural sensory context increases the variety of stimulus patterns that translate into diverse task-specific sensorimotor transformations. In recent studies of *C. elegans* that focused on a small behavioral repertoire and small space of fictive or real motor output, brain-wide dynamics exhibited low diversity (**3, 19, 47, 48**). But across the entirety of mating behavior, the brain orchestrates many different components of behavior. Each component involves different brain dynamics even when using the same motoneurons and muscle cells to produce movement. One example of diverse context-dependent circuit function is the control of tail posture. During mating, tail posture is controlled in different ways in different contexts. This is accomplished by motif-specific computations by partly overlapping circuits that converge on a common set of motoneurons and musculature, thereby producing dorsal bending, ventral bending, or undulatory movement at different times.

One set of functional correlations does not describe the diversity of brain-wide activity across the many steps of mating. Because one wiring diagram supports many patterns of functional correlation, the relationship between the connectome and brain-wide activity cannot be one-to-one. Throughout the brain, neurons have synaptic connectivity both to other neurons that participate in the same components of behavior as well as connectivity to neurons that participate in other components. Some of the significance of in-group and out-group connectivity is revealed by temporally adjacent motifs like scanning and vulva-detection. For example, in-group connectivity among vulva-detecting neurons (**13**) provides recurrent excitation that amplifies circuit activity during copulation attempts (**Fig. 6**). Signal amplification within the vulva-detecting circuit presumably enhances its all-or-none sensitivity, but then requires context-dependent inhibition to prevent spurious activation. This context-dependent inhibition is provided by out-group connectivity from the scanning circuit. Circuits for different steps of behavior are not isolated, but instead continuously interact to stabilize individual behaviors or mediate transitions between behaviors.

We conclude that diverse brain-wide dynamics emerge from the complex interactions between a male and his mating partner (**Fig. 7**). The mating circuit recognizes different sets of sensory inputs that only arise in their natural context. These inputs are manifested in brain-wide functional correlations, starting from the diverse properties of sensory neurons. Diverse sensory patterns act on the wiring diagram to recruit different sensorimotor pathways and different communities of neurons for different steps of behavior. Brain-wide dynamics during a natural behavior are not a phenomenon of an isolated nervous system, but are shaped by the flow of sensory inputs and motor actions as behavior unfolds. Behavioral dynamics must be integrated with neuronal dynamics and wiring to understand how the brain does what it evolved to do.

## Methods

### Growing conditions

All *C. elegans* strains (**Table S5**) were grown at 23°*C* on nematode growth medium (NGM) plates seeded with *E. coli* OP50. All strains were allowed to recover from starvation and freezing for at least two generations before experiments.

### Molecular biology

A *C. elegans* imaging strain ZM9624 was designed to co-express GCaMP6s and mNeptune in all neuronal nuclei. For this, *lin-15*(*n765*) hermaphrodites were co-injected with lin-15 rescuing plasmid and *pJH*3971(*Prgef* − 1 :: *GCaMP* 6*s* :: 3*xN LS* :: *mN eptune*). A transgenic line (hpEx3880) with consistent expression of GCaMP6s and mNeptune was selected for the UV-mediated transgene integration. The integrated line was out-crossed four times with N2.

ADS1002 was created by co-injecting *lin-15*(*n765*) with lin-15 rescuing plasmid and *pJH*34038(*Prgef* − 1 :: *GCaMP* 6*s* :: 3*xNLS*). A stable transgenic line (hpEx3931) was generated and outcrossed three times with N2.

### Brain-wide recordings of neuronal activity

We sought to capture multiple motifs of mating behavior within each recording session. The number and order of behavioral motifs varied from trial to trial. For comprehensive analyses, we selected seven datasets capturing a wide range of motifs. Additionally, we collected 11 datasets focusing on specific neurons and behavioral motifs and 23 datasets using ablated males (see below). It was impossible to image both the head and tail simultaneously. In this study, we focused on the neurons in the tail, most of which are male-specific and form a posterior brain that drives mating (**12**–**14**).

Virgin L4 males were selected and kept on a separate plate for 24 to 40 hours before imaging. This allowed GCaMP6s and mNeptune to accumulate in the neurons of the tail, many of which are born during the L4 stage (**12**). For the imaging experiments, a single adult male was placed on an NGM agar plate with 2-5 hermaphrodites and a small amount of OP50. We used hermaphrodites that expressed red fluorescent markers in either their muscles or epidermis for tracking (**Table S1**). The assay plate was covered with a coverslip for oil immersion. Animals could swim freely under the coverslip.

Imaging was performed using a custom spinning-disc confocal microscope described in Venkatachalam et al. (2016), with some modifications (**19**). Briefly, the emitted light from the samples was split into the red and green channels and images were captured using two Andor Zyla 4.2 sCMOS cameras. Each camera recorded 256 × 512 pixel area of interest at 200 Hz and with the system pixel size of 0.45 µm. Volumetric imaging was done using a 40x, 0.95 NA Nikon Plan Apo Lambda objective mounted on a piezoelectric stage. We sampled 10 brain volumes per second, each volume consisting of 20 optical sections approximately 1.75 µm apart. The tail of the freely-swimming male was tracked continuously by adjusting the microscope stage position with a stage controller.

### Extracting activity traces

Neuronal activity traces were extracted from raw image volumes following image pre-processing and registration. All red channel volumes were registered to the green channel through rigid transformation. A difference of Gaussian filter was applied to both channels to suppress the background noise. The datasets were subsampled to include every other volume. The volumes were converted into 5D big-data-viewer arrays and used for neuronal segmentation and tracking with MaMuT 0.27, a Fiji plugin (**49, 50**). Segmentation and tracking were done manually, with some help of the MaMuT’s automated tracking. We tracked all visible neurons for six out of seven main datasets. For the seventh dataset, we tracked all identifiable neuron types (see below). For the 11 additional and 23 ablation datasets, only neurons of interest were tracked. Fluorescent intensities (*F*^(*red*)^ and *F*^(*green*)^) were extracted by computing mean pixel values for the 2.25 × 2.25 × 3.5 µm volumes surrounding the center of each nucleus in the green and red channels. Savitzky-Golay filtering (polynomial order of 1 and frame length 13) was applied to intensity traces from each channel for noise-reduction (**51**). We computed neuronal activity as the ratio *F*^(*green*)^*/F*^(*red*)^ to minimize the effects of correlated noise and motion artifacts. To mitigate remaining motion artifacts, a singular value decomposition was applied to the main datasets retaining 2/3 of components.

### Quantifying behavior

We parameterized several continuous features of male behavior. These included swimming velocity and speed, sliding velocity and speed (tail velocity and speed relative to the hermaphrodite body when in contact), tail curvature, distance of the male tail to the head and tail of the hermaphrodite and to the vulva, and spicule protraction. To parameterize velocity, we computed the centroid of the tail from coordinates of three tail neurons (AS11, PLM, EF1). Centroid positions were extracted for each volume and converted into velocity of the worm in the egocentric coordinates (**Fig. S9**). Negative velocity values correspond to swimming forwards, and positive values to swimming backwards. Tail curvature was estimated using the angle between two rays diverging from the center of PVY and intersecting centers of AS11 and PLM (**Fig. S9**). When the tail is relaxed, the angle formed by the rays is nearly straight. Spicule protraction was estimated indirectly by measuring the distance between SPC, which moves with the spicules, and PCB. As the spicules become protracted, the distance between SPC and PCB sharply decreases.

To map the position of the tail to the hermaphrodite, we extracted maximum intensity projections (MIPs) of the volumes in which large parts of the hermaphrodite were visible. The images were tiled to cover the entire body and aligned with rigid rotations and translations to produce a straightened representation of the hermaphrodite (**Fig. S10A–C**). For each data volume, we mapped the position of the tail to the straightened hermaphrodite by matching uniquely identifiable MIP features (**Fig. S10D**). Finally, we estimated the best fit for an ellipse of aspect ratio 10 given the x and y coordinates of the tail position on the hermaphrodite, generating a representation of the tail trajectory on an idealized elliptical hermaphrodite (**Fig. S10E**). Using this map, we calculated the male’s scanning velocity and speed and its distance to the vulva and to the head and tail of the hermaphrodite.

The distance of the tail to the vulva is the arc length of the shortest path between the tail and the vulva along a section of the ellipse representing the hermaphrodite. The distance of the tail to the hermaphrodite tips is the arc length of the shortest path between the male tail and either the head or the tail of the hermaphrodite along a section of the ellipse.

### Neuron identification

Neurons were identified based on a combination of features, including their position, morphology, and expression of specific fluorescent markers. **Table S2** lists key criteria for assigning IDs to specific neurons. Anatomical drawings in Sulston et al. (1980) were a primary reference for the neurons’ relative positions and morphology (**12**). Neuronal positions were additionally cross-checked with the annotated electron microscopy data for the N2Y male shared by Scott Emmons. Among other sources, we used fluorescent microphotographs in Serrano-Saiz et al. (2017) which provided a comprehensive account of the expression patterns of multiple sparse markers (**52**). In many instances, to confirm neuronal IDs, we crossed existing well-characterized marker strains with panneuronal ADS1002 or ZM9624. This approach worked well for the regions with stereotyped cell positions. Positions of some neurons are variable (**12**). To confirm identities of such neurons we performed panneuronal imaging using sparsely labeled strains, and tested if neuron-specific activity patterns matched. In total, we were able to identify 76 neurons across all datasets.

### Cross-correlation analysis and hierarchical clustering

Left-right symmetric pairs of sensory neurons showed positively correlated activity (*r* = 0.73 ±0.19), with one exception – R1A (**Fig. S2**). We considered each left-right correlated pair to represent a distinct neuronal type and compiled datasets to include the activity traces of 57 neuron types and one non-neuronal cell (int9R). We used 58 activivty traces and eight continuous behavioral parameters for cross-correlation analyses. Pairwise cross-correlations between time series were calculated for each dataset, with the lag parameter set to 5 (corresponding to one second). For each pairwise comparison, we calculated mean cross-correlations at each time lag, and the highest mean cross correlation value was included in the consensus cross-correlation matrix. This consensus correlation matrix was used to calculate pairwise Euclidian distances for hierarchical clustering with the Ward’s minimum variance method (**53**).

### Event-triggered activity averages

We tested how neurons of the mating circuit responded to the start of specific behavioral events (**Fig. 3D**, S1). Events included were: ventral contact with the hermaphrodite (both the initial contact and any contact preceded by more than 6 seconds of separation); hermaphrodite loss lasting for more than 6 seconds; turning; failed turning attempts; vulva contact, copulation (as identified by increased calcium signals in the intestinal cells); switching from forward to backward swimming (tabulated for the entire behavior, and separately for searching, and scanning); switching from backward to forward swimming (tabulated for the entire behavior, and separately for searching, and scanning). Switching from forward to backward sliding (movement of the tail relative to the hermaphrodite); switching from backward to forward sliding. Neuronal signals were extracted for a window spanning 6 seconds before and after the events, and dataset-averaged response curves were calculated for each neuron and event. We then calculated cross-correlations between these response curves and binarized event traces, with the lag parameter set to 5. The lag with the maximum absolute mean correlation was selected. One-sample t-test was used to test whether the correlation estimate between the neurons’ response curves at that lag and behaviors was greater than zero. *p*-values were adjusted using the FDR method to correct for multiple tests. The FDR adjusted significance cut-off was set at 0.05. Activity changes at the onset of discrete behavioral events (i.e. significant increases, decreases, no changes) were used to calculate the fraction of neurons with unique activity patterns.

### Decoding behavior from neuronal activity

To predict behavior from neuronal activity traces, we used generalized linear models with elastic-net regularization (**54**). The procedure was adapted from Scholz et al. (2018) with modifications (**6**). We sought predictions across different individuals. Activity traces of all neurons were centered and scaled to have a zero sample mean and unit sample variance, and concatenated for all datasets. We also concatenated time series of all behavioral features. Traces of several neurons were absent from some datasets. Missing traces were imputed if they were missing from single datasets. Activity traces missing from more than one dataset were excluded. To instantiate short-term memory, we used time-lags in the neuronal activity traces. Three time lags were used (*t* = 0, *t* = 0.6, and *t* = 1.2 seconds), that maximized the accuracy of predictions.

We partitioned concatenated time series into training and testing sets. For each modeling iteration, we selected six datasets for training and one for testing the resulting model’s out-of-sample prediction accuracy. The analysis was repeated to include all possible combinations of training and testing data. Model training and evaluation was performed using R packages ‘Caret’ version 6.0-84 (**55**) and ‘glmnet’ version 2.0-18 (**54, 56**). The algorithm fits a generalized linear model via penalized maximum likelihood. Tuning parameters include the penalty term *λ*, controlling the penalty strength, and the elastic-net mixing parameter α (where *α* = 0 is the ridge penalty and *α* = 1 is the lasso penalty). We tuned the model’s parameters by searching 101 α values (*α* ∈ [0, 1]) and 101 logarithmically-spaced λ values (*λ* ∈ [10^−5^, 10^1^]). The best tuning parameters were selected based on the root mean square error and 5-fold cross-validation. The models’ prediction accuracy was estimated by calculating the mean *R*^2^ between out-of-sample predictions and target variables.

To better understand contributions of individual neurons, we ranked neurons by their weight, averaging the weight of all included time lags for a given neuron. We also evaluated additional models which were trained leaving out traces of single neurons, as well as models trained on traces of individual neurons and their time lags.

### State-specific correlations

To compare correlation structure across behavioral states, we calculated neuron-neuron correlation matrices for each individual and mating state (searching, scanning, copulation, and resting) (**Fig. 5**). We calculated the Euclidian distances between all pairs of correlation matrices and applied non-metric multidimensional scaling using Sammon’s non-linear mapping to generate a 2-dimentional representation of the similarity/ dissimilarity between correlation matrices (**57**). We also compared correlations between neurons and continuous behavioral features (velocity, curvature, and spicules position) across behavioral states and datasets.

### Comparisons between functional and synaptic connectivity

As a measure of functional weight, we used consensus cross-correlations (**Fig. 3B**). When both members of a left-right neuron pair were included in the analyses, their activity was assumed to be perfectly correlated, except in the case of R1A. As a measure of synaptic connectivity, we used the number of synapses between neurons from https://wormwiring.org/ (**13, 14**). Identifying connections between neurons, particularly gap junctions, can involve a degree of subjectivity (**58**), so we additionally validated gap junctions of the male tail by screening EM photographs of all annotated gap junctions in the N2Y male. We assigned a confidence score from 1 to 5 to each gap junction. Synapses with confidence scores 3 and higher were used to compile a validated gap junction connectivity matrix. To minimize the effects of observational noise, we also symmetrized the connectivity for all neurons that are bilaterally symmetrical, except R1A. Only connections involving more than one synapse were used for all analyses.

To test for the correlation between functional and synaptic connectivity, we calculated Pearson and partial correlation coefficients between the functional weight and the number of shared electrical and chemical synapses. To test for the correlation between the functional weight and the number of chemical synapses, we used the absolute value of functional connectivity. We tested for significance using 10000 shuffled functional connectivity matrices. To test if the results were sensitive to a particular choice of the connectivity hyperparameters, the analyses were performed using 16 variants of connectivity matrices. These matrices were generated using (i) synapse number or weight as a measure of connectivity ((**13**)), (ii) validated or original gap junctions, (iii) symmetrized or original connectivity, and (iv) both neurons of left-right pairs or only one neuron of each pair. The results of these analyses are summarized in **Table S4**.

### Functional communities and community interaction types

To extract overlapping community structure we used link clustering (**20, 21**). For link clustering, functional correlations higher than 0.3 were selected from the the consensus cross-correlation matrix; left-right pairs of laterally symmetrical neurons were included. Extracted link communities were clustered themselves with the Ward’s minimum variance method (**53**) and a deprogram cut-off parameter set at 0.85, resulting in eleven meta-communities. Community interaction motifs were identified as described in Betzel et al., 2018 (**22**). Synaptic connectivity density was computed using the number of chemical and electrical synaptic connections, including connections between nodes present in both communities.

### Neuronal ablations

All ablations were performed in young adult males, 1-4 hours after they completed their final molt. Ablations were done using a MicroPoint pulsed laser system (Andor) mounted on a Nikon microscope equipped with Nomarski optics and a 100x 0.95 NA Nikon Plan Apo objective. Males were placed on a 10% agar pad and immobilized with 0.01 µm latex beads (Polysciences, Inc). Target neurons were identified based on the expression of fluorescent markers. The neuronal nuclei were targeted until a visible damage appeared. Mock ablations were performed by placing males on a slide with latex beads for 5-10 minutes. Males were left to recover overnight and behavioral assays and imaging experiments were performed on the next day.

### Mating assays

Behavioral arenas were 6 cm NGM agar plates seeded with a thin layer of *E. coli* OP50 forming a spot approximately 0.5 cm in diameter. One to three young adult hermaphrodites and one male were placed onto each assay plate. Behavior of the male was recorded at 5 frames per second for 30 to 60 minutes using a Grasshopper3 camera and Spinnaker software (Point Grey). Treatment and control trials were recorded on the same day. The recordings were analyzed blindly in MaMuT. Two-sided t-tests were used to compare group means.

For the PVV ablations, we tabulated successful turns and failed turning attempts. A turn was considered failed when the turn initiation was followed by either a permanent or temporary loss of contact.

To quantify pausing at the excretory pore, we performed mating assays with vulvaless *let-23* hermaphrodites. We recorded when the male stopped backward scanning for more than 0.2 seconds on the anterior ventral side opposite the terminal bulb of the pharynx. This location corresponds to the excretory pore opening (**Fig. S5A**). For comparison, we tabulated pausing on the anterior dorsal side. Pausing was recorded during backward scanning towards the tip of the hermaphrodite’s head (**Fig. S5B**).

For the R2B ablations, we tested whether vulva detection and retention were compromised. We calculated the fraction of successful vulva detection events. Vulva detection was considered successful if the male paused scanning at the vulva for more than 0.2 seconds. We also calculated the average vulva retention time. Assays with more than five vulva detection events were included; vulva contacts during copulation were excluded.

## Acknowledgements

We thank Alkmini Chalofti and Noah Milstein for help with neuronal tracking, Albert Lin for creating hpEx3931, Oliver Hobert for sharing strains, Scott Emmons for sharing EM photographs of the male nervous system. We thank Scott Linderman, Paul Sternberg, and members of the Samuel lab for their comments on the manuscript. Some strains were provided by the CGC, which is funded by NIH Office of Research Infrastructure Programs (P40 OD010440). This work was supported by NSF (IOS-1452593), NIH (R01 NS082525, R01 GM130842-01, and U01-NS111697), Canadian Institutes of Health Research (Foundation Scheme 154274), Natural Sciences and Engineering Research Council of Canada (RGPIN-2017-06738), and Burroughs Wellcome Fund CASI (VV).

## Supplementary Materials

**Table S1.**
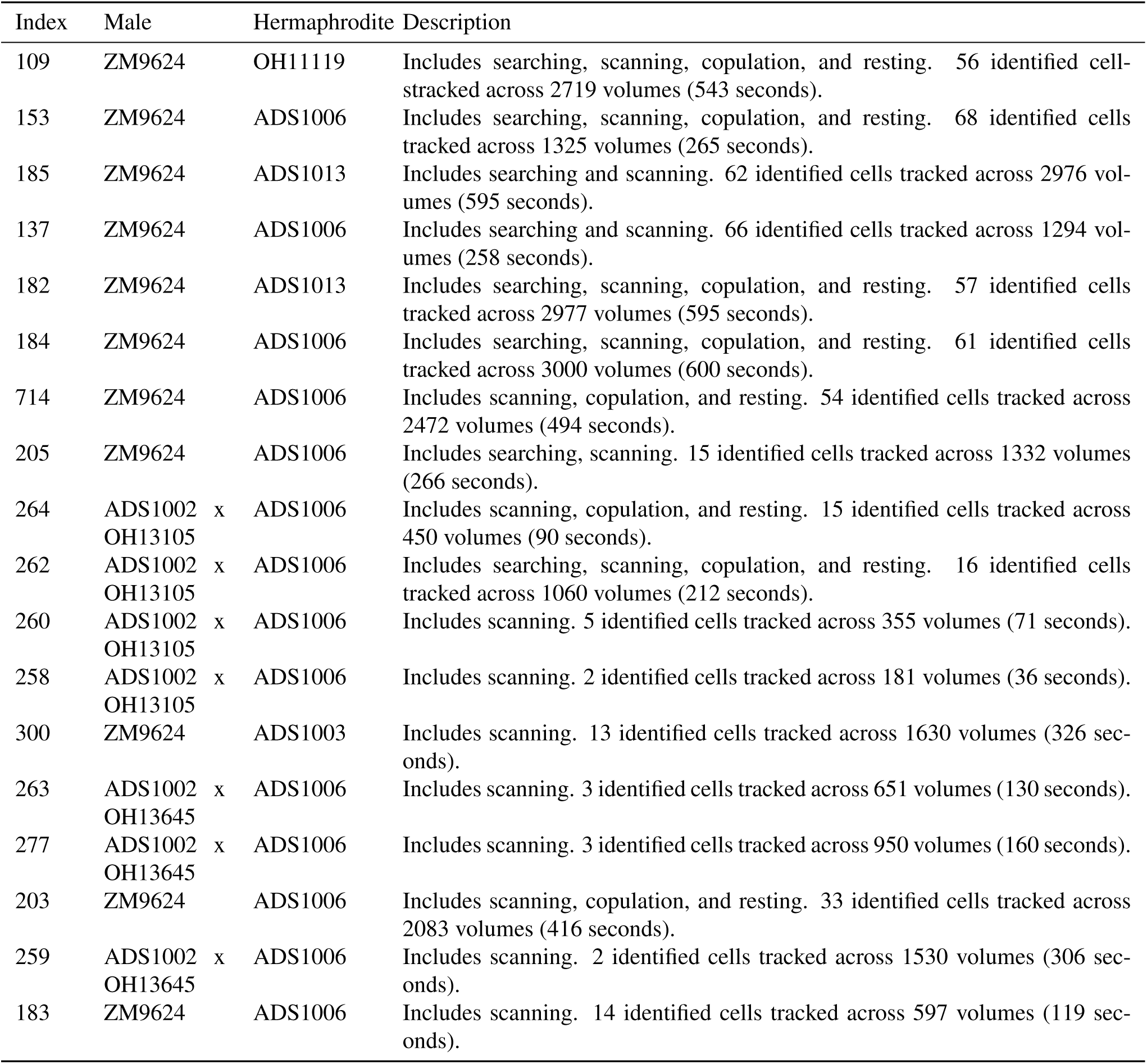
Datasets. Functional analyses were performed on seven main and eleven additional datasets. For the main datasets, activity traces were extracted for all identifiable neurons. For the eleven additional datasets, only neurons of interest were tracked.

**Table S2.**
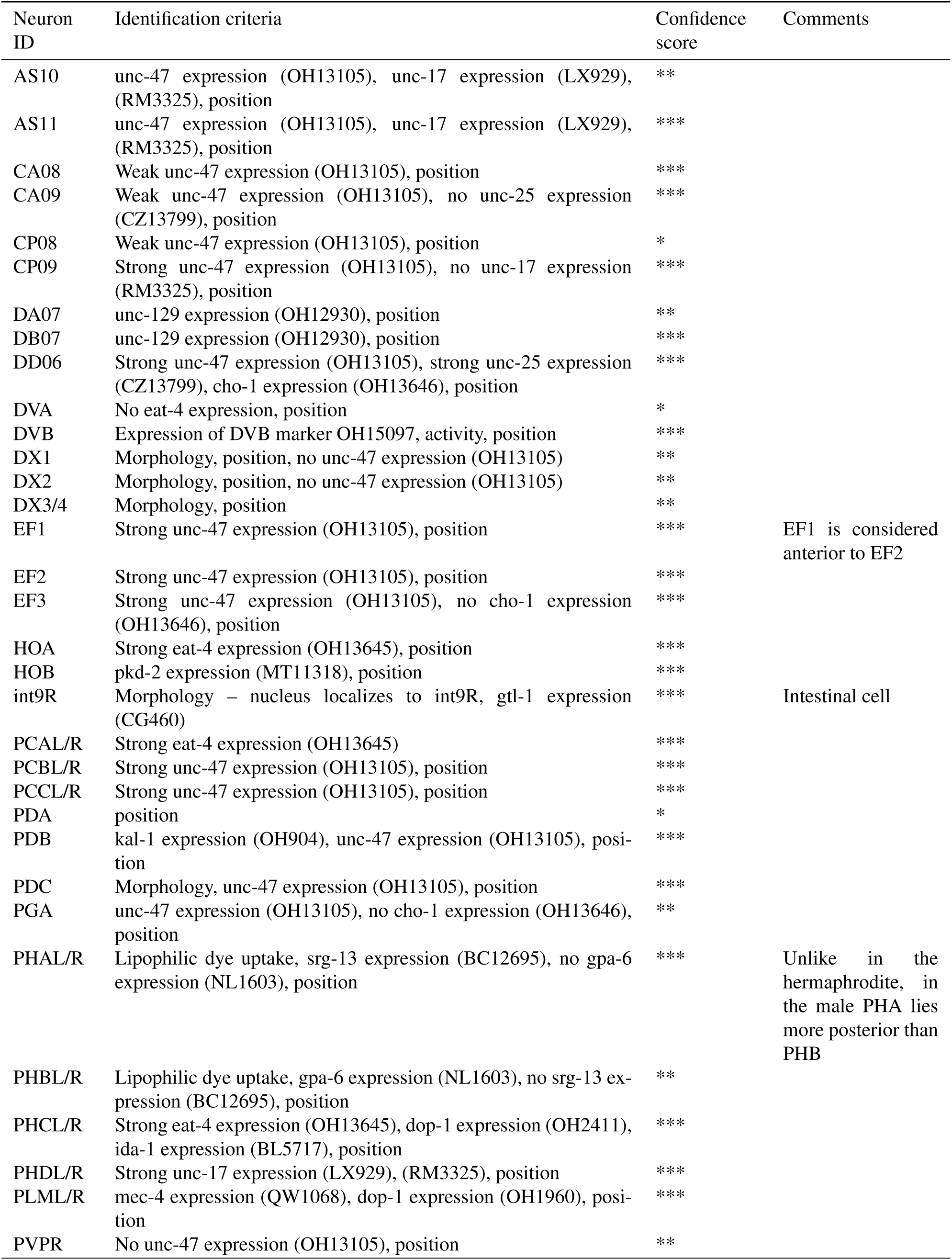
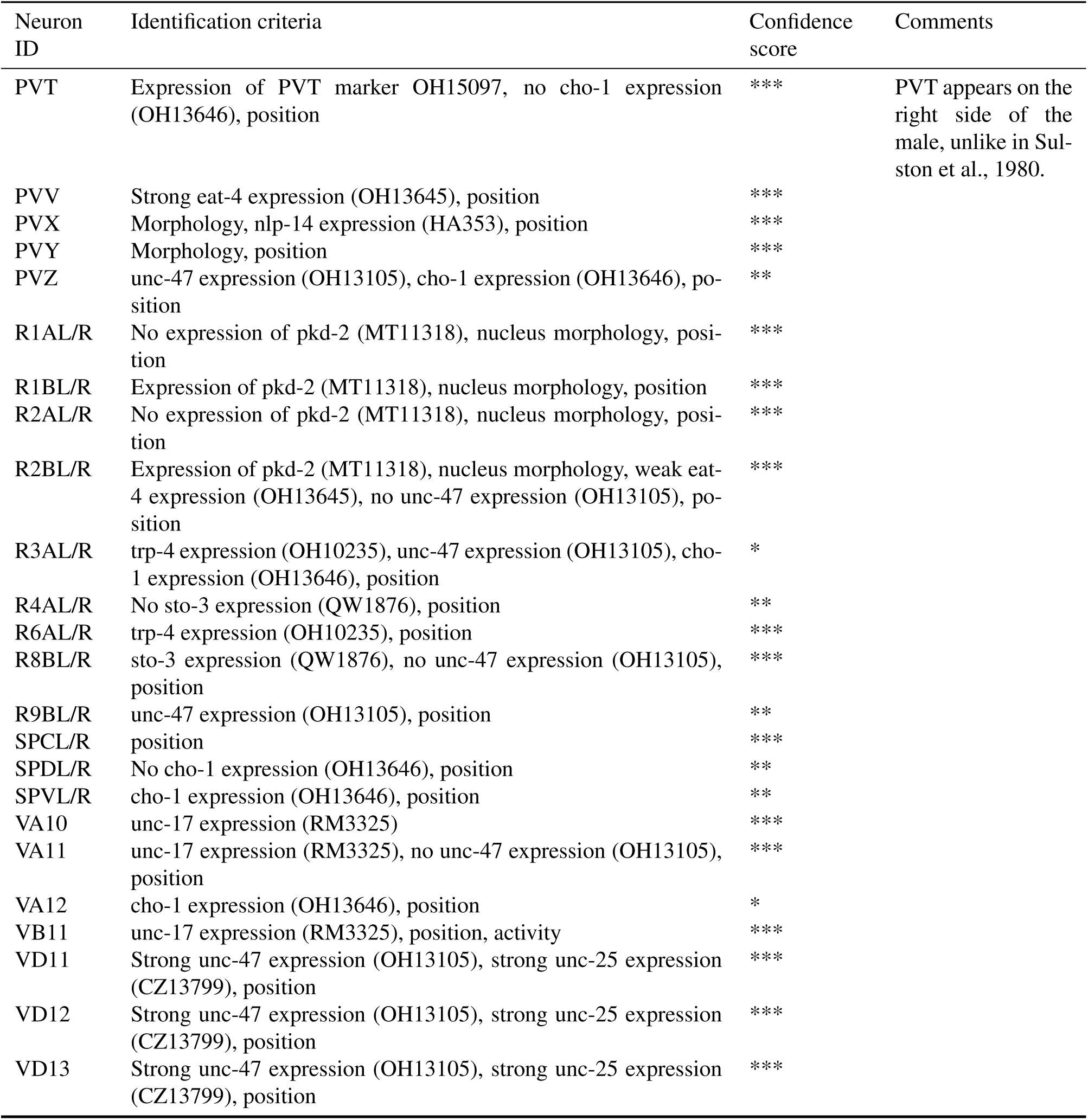
Neuron identification. 57 neuron types and the intestinal cell nucleus int9R were identified based on their morphology, position, and expression of specific fluorescent markers. Confidence scores: *** certain, ** confident, * probable.

| Neuron ID | Identification criteria | Confidence score | Comments |
| --- | --- | --- | --- |
| AS10 | unc-47 expression (OH13105), unc-17 expression (LX929), (RM3325), position | ** |  |
| AS11 | unc-47 expression (OH13105), unc-17 expression (LX929), (RM3325), position | *** |  |
| CA08 | Weak unc-47 expression (OH13105), position | *** |  |
| CA09 | Weak unc-47 expression (OH13105), no unc-25 expression (CZ13799), position | *** |  |
| CP08 | Weak unc-47 expression (OH13105), position | * |  |
| CP09 | Strong unc-47 expression (OH13105), no unc-17 expression (RM3325), position | *** |  |
| DA07 | unc-129 expression (OH12930), position | ** |  |
| DB07 | unc-129 expression (OH12930), position | *** |  |
| DD06 | Strong unc-47 expression (OH13105), strong unc-25 expression (CZ13799), cho-1 expression (OH13646), position | *** |  |
| DVA | No eat-4 expression, position | * |  |
| DVB | Expression of DVB marker OH15097, activity, position | *** |  |
| DX1 | Morphology, position, no unc-47 expression (OH13105) | ** |  |
| DX2 | Morphology, position, no unc-47 expression (OH13105) | ** |  |
| DX3/4 | Morphology, position | ** |  |
| EF1 | Strong unc-47 expression (OH13105), position | *** | EF1 is considered anterior to EF2 |
| EF2 | Strong unc-47 expression (OH13105), position | *** |  |
| EF3 | Strong unc-47 expression (OH13105), no cho-1 expression (OH13646), position | *** |  |
| HOA | Strong eat-4 expression (OH13645), position | *** |  |
| HOB | pkd-2 expression (MT11318), position | *** |  |
| int9R | Morphology – nucleus localizes to int9R, gtl-1 expression (CG460) | *** | Intestinal cell |
| PCAL/R | Strong eat-4 expression (OH13645) | *** |  |
| PCBL/R | Strong unc-47 expression (OH13105), position | *** |  |
| PCCL/R | Strong unc-47 expression (OH13105), position | *** |  |
| PDA | position | * |  |
| PDB | kal-1 expression (OH904), unc-47 expression (OH13105), position | *** |  |
| PDC | Morphology, unc-47 expression (OH13105), position | *** |  |
| PGA | unc-47 expression (OH13105), no cho-1 expression (OH13646), position | ** |  |
| PHAL/R | Lipophilic dye uptake, srg-13 expression (BC12695), no gpa-6 expression (NL1603), position | *** | Unlike in the hermaphrodite, in the male PHA lies more posterior than PHB |
| PHBL/R | Lipophilic dye uptake, gpa-6 expression (NL1603), no srg-13 expression (BC12695), position | ** |  |
| PHCL/R | Strong eat-4 expression (OH13645), dop-1 expression (OH2411), ida-1 expression (BL5717), position | *** |  |
| PHDL/R | Strong unc-17 expression (LX929), (RM3325), position | *** |  |
| PLML/R | mec-4 expression (QW1068), dop-1 expression (OH1960), position | *** |  |
| PVPR | No unc-47 expression (OH13105), position | ** |  |
| PVT | Expression of PVT marker OH15097, no cho-1 expression (OH13646), position | *** | PVT appears on the right side of the male, unlike in Sulston et al., 1980. |
| PVV | Strong eat-4 expression (OH13645), position | *** |  |
| PVX | Morphology, nlp-14 expression (HA353), position | *** |  |
| PVY | Morphology, position | *** |  |
| PVZ | unc-47 expression (OH13105), cho-1 expression (OH13646), position | ** |  |
| R1AL/R | No expression of pkd-2 (MT11318), nucleus morphology, position | *** |  |
| R1BL/R | Expression of pkd-2 (MT11318), nucleus morphology, position | *** |  |
| R2AL/R | No expression of pkd-2 (MT11318), nucleus morphology, position | *** |  |
| R2BL/R | Expression of pkd-2 (MT11318), nucleus morphology, weak eat-4 expression (OH13645), no unc-47 expression (OH13105), position | *** |  |
| R3AL/R | trp-4 expression (OH10235), unc-47 expression (OH13105), cho-1 expression (OH13646), position | * |  |
| R4AL/R | No sto-3 expression (QW1876), position | ** |  |
| R6AL/R | trp-4 expression (OH10235), position | *** |  |
| R8BL/R | sto-3 expression (QW1876), no unc-47 expression (OH13105), position | *** |  |
| R9BL/R | unc-47 expression (OH13105), position | ** |  |
| SPCL/R | position | *** |  |
| SPDL/R | No cho-1 expression (OH13646), position | ** |  |
| SPVL/R | cho-1 expression (OH13646), position | ** |  |
| VA10 | unc-17 expression (RM3325) | *** |  |
| VA11 | unc-17 expression (RM3325), no unc-47 expression (OH13105), position | *** |  |
| VA12 | cho-1 expression (OH13646), position | * |  |
| VB11 | unc-17 expression (RM3325), position, activity | *** |  |
| VD11 | Strong unc-47 expression (OH13105), strong unc-25 expression (CZ13799), position | *** |  |
| VD12 | Strong unc-47 expression (OH13105), strong unc-25 expression (CZ13799), position | *** |  |
| VD13 | Strong unc-47 expression (OH13105), strong unc-25 expression (CZ13799), position | *** |  |

**Table S3.** Functional imaging in ablated males. Neurons of the vulva-detecting circuit were ablated in young adult males, and the panneuronal imaging was performed on the next day. For this, the males were placed on a plate with hermaphrodites and were allowed to mate. Traces of activity were extracted for several neurons of interest. A few neurons of interest could not be seen in some males. These neurons are marked with ‘-’.

| Ablation | Index | PCB | PCC | HOA | HOB | R2B | PVX | SPC | PCA |
| --- | --- | --- | --- | --- | --- | --- | --- | --- | --- |
| -HOA | 102 | + | + | na | + | + | + | + | + |
| -HOA | 105 | + | + | na | + | + | + | + | + |
| -HOA | 101 | + | + | na | + | + | + | + | + |
| -PCA/L | 492 | + | + | + | + | + | + | + | na |
| -PCA/L | 501 | + | + | + | + | - | + | + | na |
| -PCA/L | 505 | + | + | + | + | + | + | + | na |
| -HOA -PCB/L | 450 | na | + | na | + | + | + | + | + |
| -HOA -PCB/L | 453 | na | + | na | - | + | + | + | - |
| -HOA -PCB/L | 1023 | na | + | na | + | - | + | + | - |
| -HOA -PCC/L | 568 | + | na | na | + | + | + | + | + |
| -HOA -PCC/L | 1025 | + | na | na | + | + | + | + | + |
| -HOA -PCC/L | 1032 | + | na | na | + | + | + | + | + |
| -HOA -HOB | 560 | + | - | na | na | - | + | + | + |
| -HOA -HOB | 561 | + | - | na | na | - | + | + | + |
| -HOA -HOB | 562 | + | + | na | na | + | + | + | + |
| -HOA -HOB | 1022 | + | + | na | na | + | + | + | + |

**Table S4.** Correlations between functional and synaptic connectivity. We calculated Pearson and partial corralation coefficients on 16 variants of synaptic connectivity matrices. The different variants of the connectivity matrices were generated using (i) both neurons of the same type or one neuron per neuronal type, (ii) symmetrized or original connectivity, (iii) validated or original gap junctions, and (iv) synapse number or synaptic weight as a measure of connectivity.

| Both laterally symmetrical neurons included | Symmetrized connectivity | Validated gap junctions | Connectivity measure | Chemical |  | Electrical |  |
| --- | --- | --- | --- | --- | --- | --- | --- |
|  |  |  |  | Pearson correlation | Partial correlation | Pearson correlation | Partial correlation |
| yes | yes | yes | Synapse n | 0.17<br>p=.0013 | 0.12<br>p=.021 | 0.35<br>p=.001 | 0.34<br>p=.0007 |
| yes | yes | yes | Synaptic w | 0.11<br>p=.012 | 0.08<br>p=.069 | 0.29<br>p=.0035 | 0.29<br>p=.001 |
| yes | yes | no | Synapse n | 0.17<br>p=.001 | 0.13<br>p=.0097 | 0.33<br>p=.0001 | 0.33<br>p=0 |
| yes | yes | no | Synaptic w | 0.11<br>p=.014 | 0.08<br>p=.087 | 0.37<br>p=0 | 0.37<br>p=0 |
| yes | no | yes | Synapse n | 0.14<br>p=.0075 | 0.10<br>p=.069 | 0.37<br>p=.0001 | 0.36<br>p=0 |
| yes | no | yes | Synaptic w | 0.11<br>p=.023 | 0.08<br>p=.081 | 0.30<br>p=.0024 | 0.30<br>p=.0012 |
| yes | no | no | Synapse n | 0.14<br>p=.0072 | 0.11<br>p=.034 | 0.36<br>p=0 | 0.35<br>p=0 |
| yes | no | no | Synaptic w | 0.11<br>p=.022 | 0.08<br>p=.088 | 0.38<br>p=0 | 0.37<br>p=0 |
| no | yes | yes | Synapse n | -0.01<br>p=.86 | -0.07<br>p=.32 | 0.46<br>p=.0001 | 0.48<br>p=.0004 |
| no | yes | yes | Synaptic w | -0.02<br>p=.75 | -0.07<br>p=.26 | 0.3<br>p=.015 | 0.34<br>p=.0048 |
| no | yes | no | Synapse n | -0.01<br>p=.86 | -0.06<br>p=.4 | 0.37<br>p=0 | 0.38<br>p=0 |
| no | yes | no | Synaptic w | -0.02<br>p=.74 | -0.07<br>p=.22 | 0.37<br>p=0 | 0.38<br>p=0 |
| no | no | yes | Synapse n | 0.01<br>p=.84 | 0.04<br>p=.61 | 0.38<br>p=.0017 | 0.4<br>p=.0008 |
| no | no | yes | Synaptic w | 0.02<br>p=.72 | 0.03<br>p=.65 | 0.28<br>p=.025 | 0.32<br>p=.008 |
| no | no | no | Synapse n | 0.01<br>p=.85 | 0.02<br>p=.79 | 0.37<br>p=0 | 0.37<br>p=0 |
| no | no | no | Synaptic w | 0.02<br>p=.72 | 0.02<br>p=.75 | 0.38<br>p=0 | 0.38<br>p=0 |

**Table S5.** List of *C. elegans* strains used in this study.

| Reagent | Source |
| --- | --- |
| ADS1002 aeaIs010 [Prgef-1::GCaMP6s::3xNLS + lin-15(+)] | This study |
| ADS1003 otIs377 [myo-3p::mCherry]; let-23(sy1) II; unc-64(e246) III | This study |
| ADS1006 uuEx18 [dcr-1(wild-type) + dpy-30::mCherry]; lin-15 (n765ts) X | This study |
| ADS1013 otIs377 [myo-3p::mCherry]; unc-31(e169) IV | This study |
| BB92 uuEx18 [dcr-1(wild-type) + dpy-30::mCherry] | CGC |
| BC12695 dpy-5(e907) I; sIs12409 [rCes T23F11.5::GFP + pCeh361] | CGC |
| BL5717 inIs179 [ida-1p::GFP] II; him-8(e1489) IV | CGC |
| CB169 unc-31 (e169) IV | CGC |
| CB246 unc-64 (e246) III | CGC |
| CB4088 him-5 (e1490) V | CGC |
| CZ13799 juIs76 [unc-25p::GFP + lin-15(+)] II | CGC |
| HA353 lin-15B&lin-15A (n765) X; rtEx247 [nlp-14p::GFP + lin-15(+)] | CGC |
| LX1986 vsIs177 [ocr-2p::GCaMP5::ocr-2 3'UTR + ocr-2p::mCherry::ocr-2 3'UTR + lin-15(+)] lite-1(ce314) lin-15AB(n765) X | CGC |
| LX929 vsIs48 [unc-17::GFP] | CGC |
| MT11318 nIs133 [pkd-2::GFP] I; ceh-30 (n3714) X | CGC |
| N2 wildtype | CGC |
| NL1603 dpy-20(e1282) IV; pkIs583 [gpa-6::GFP + dpy-20(+)] | CGC |
| OH10235 ceh-43(tm480) III; him-5 (e1490) V; bxIs19 [trp-4p::GFP]; norEx41 [ceh-43 fosmid + dat-1::mCherry] | CGC |
| OH11119 otIs377 [myo-3p::mCherry]; otEx4945 [hsp16-2p::hlh-1::2xFLAG + rol-6(su1006)] | CGC |
| OH11992 otIs452[eat-4p9::GFP + pha-1(+)] | Oliver Hobert |
| OH13083 him-5(e1490) V; otIs576 [unc-17(fosmid)::GFP + lin-44::YFP] | CGC |
| OH13105 him-5 (e1490) otIs564 [unc-47(fosmid)::SL2::H2B::mChopti + pha-1(+)] V | CGC |
| OH13645 pha-1(e2123) III; him-5(e1490) V; otIs518 [eat-4(fosmid)::SL2::mCherry::H2B + pha-1(+)] | CGC |
| OH13646 pha-1 (e2123) III; him-5 (e1490) V; otIs544 [cho-1(fosmid)::SL2::mCherry::H2B + pha-1(+)] | CGC |
| OH15039 pha-1(e2123) III; him-5 (e1490) V; otIs625 [cat-1(fosmid)::SL2::mCherry::H2B + pha-1(+)] | CGC |
| OH15097 him-5(e1490) V; otIs541 [lim-6(intron4)::wCherry] | CGC |
| OH2411 pha-1(e2123) III; otEx233 [dop-1(prom1)::GFP + pha-1(+)] | CGC |
| OH904 otIs33[kal-1::GFP] IV | CGC |
| PS21 let-23 (sy1) II; him-5 (e1490) V | CGC |
| QW1068 lin-15 (n765ts) lite-1 (ce314) X; zfEx436 [mec-4::mCherry::SL2::GCaMP6s] | Mark Alkema |
| QW1876 zfEx898[Psto-3::wrmScarlet + lin-15(+)]; lin-15 (n765ts) X | Susoy et al., 2019 |
| RM3325 pha-1 (e2123) III; mdEx865 [unc-17p::NLS::mCherry + pha-1(+)] | CGC |
| ZM8516 hpIs474[glr-1::NLS::wCherry + lin-15(+)] | Mei Zhen |
| ZM9624 hpIs675 [Prgef-1::GCaMP6s::3xNLS::mNeptune + lin-15(+)] lin-15(n765) X | This study |

**Fig. S1.**
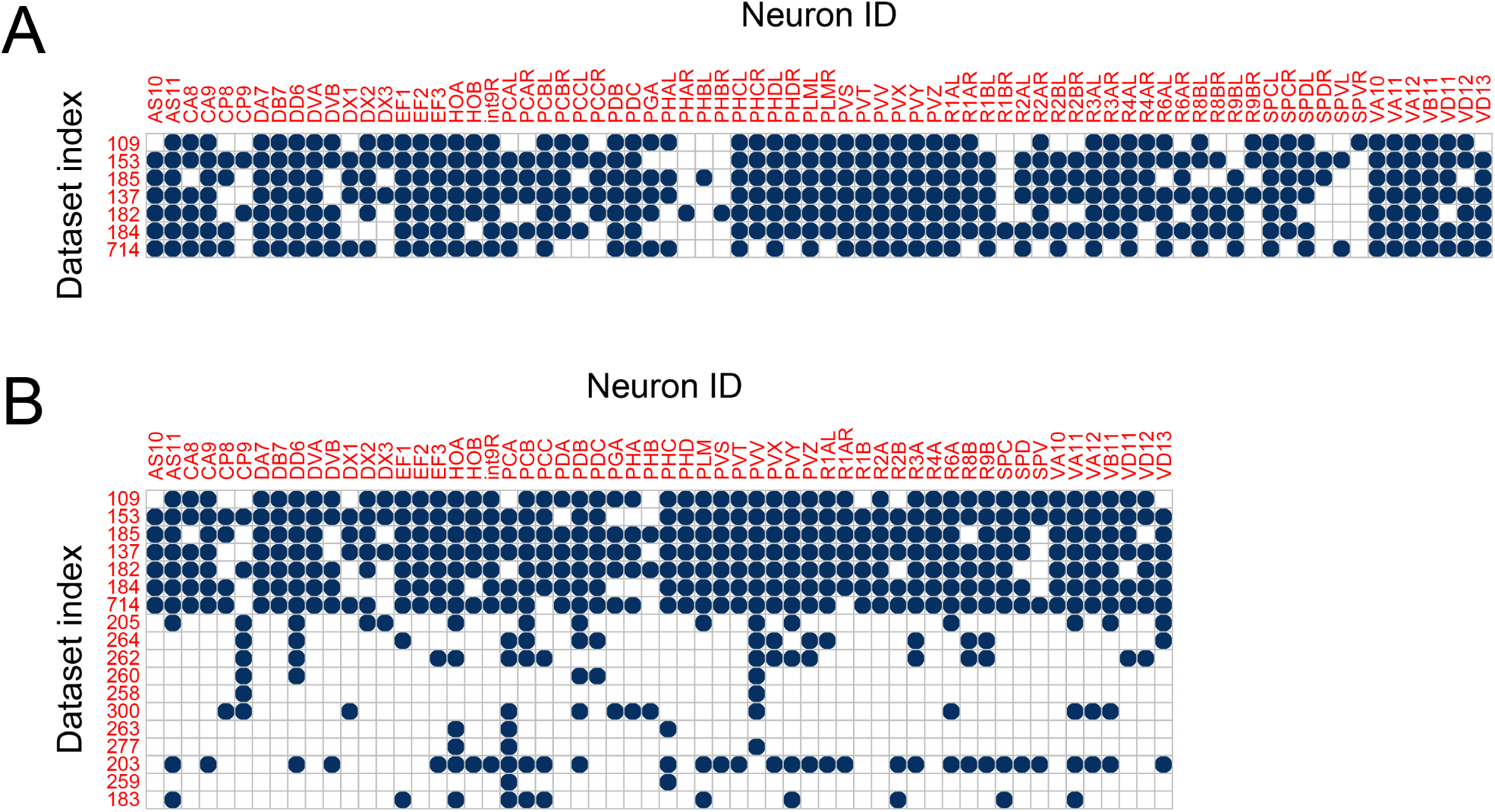
Neurons and datasets. 76 individual neurons and one non-neuronal cell type (int9R) were identified across 7 main datasets (**A**). The neurons were assembled into 57 neuron types for the 7 main and 11 additional datasets (**B**).

**Fig. S2.**
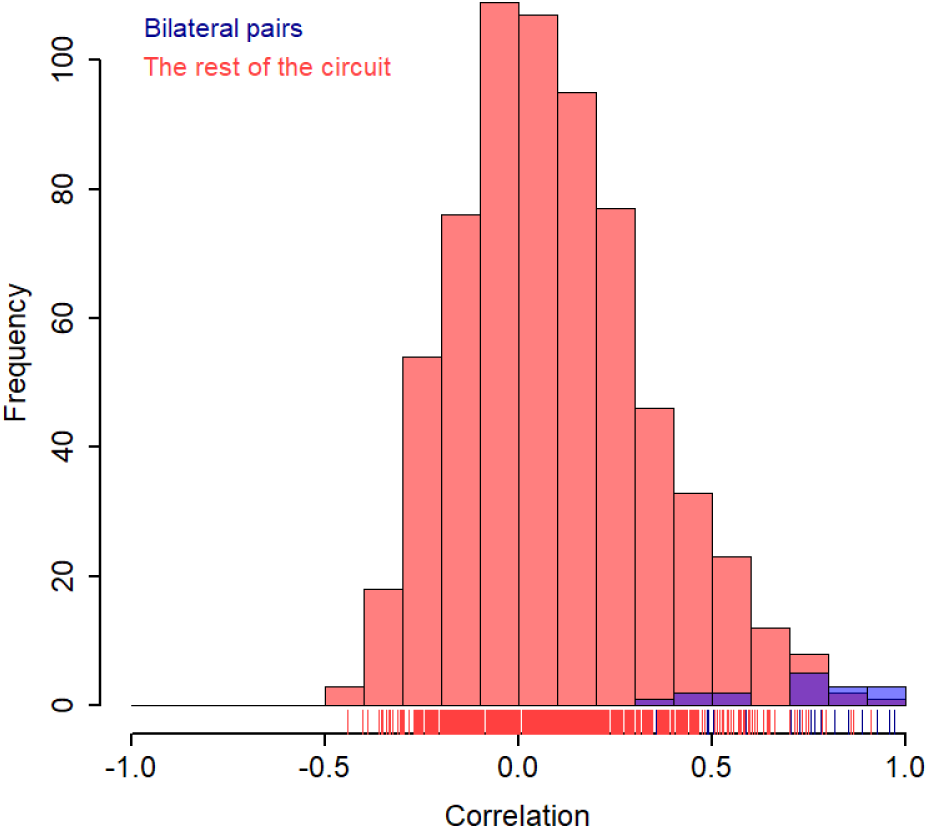
Laterally-symmetrical neurons show correlated activity. Distribution of cross-correlation coefficients between left-right neuron pairs of the same type (blue) and distribution of correlation values between different neuron types (red).

**Fig. S3.**
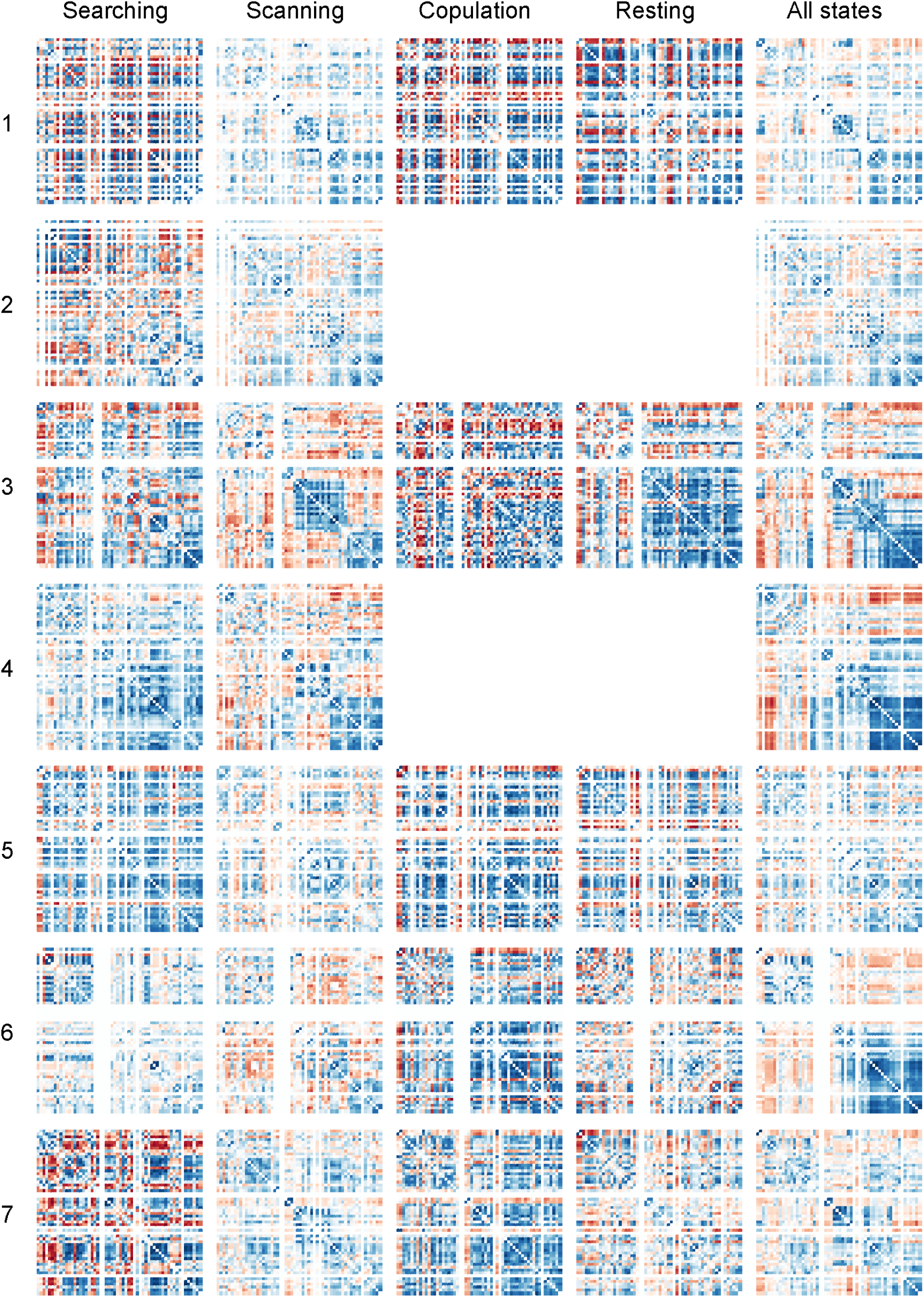
Dataset-specific and state-specific correlations. Cross-correlation matrices for individual datasets and partitioned by four behavioral states: searching, scanning, copulation, and resting.

**Fig. S4.**
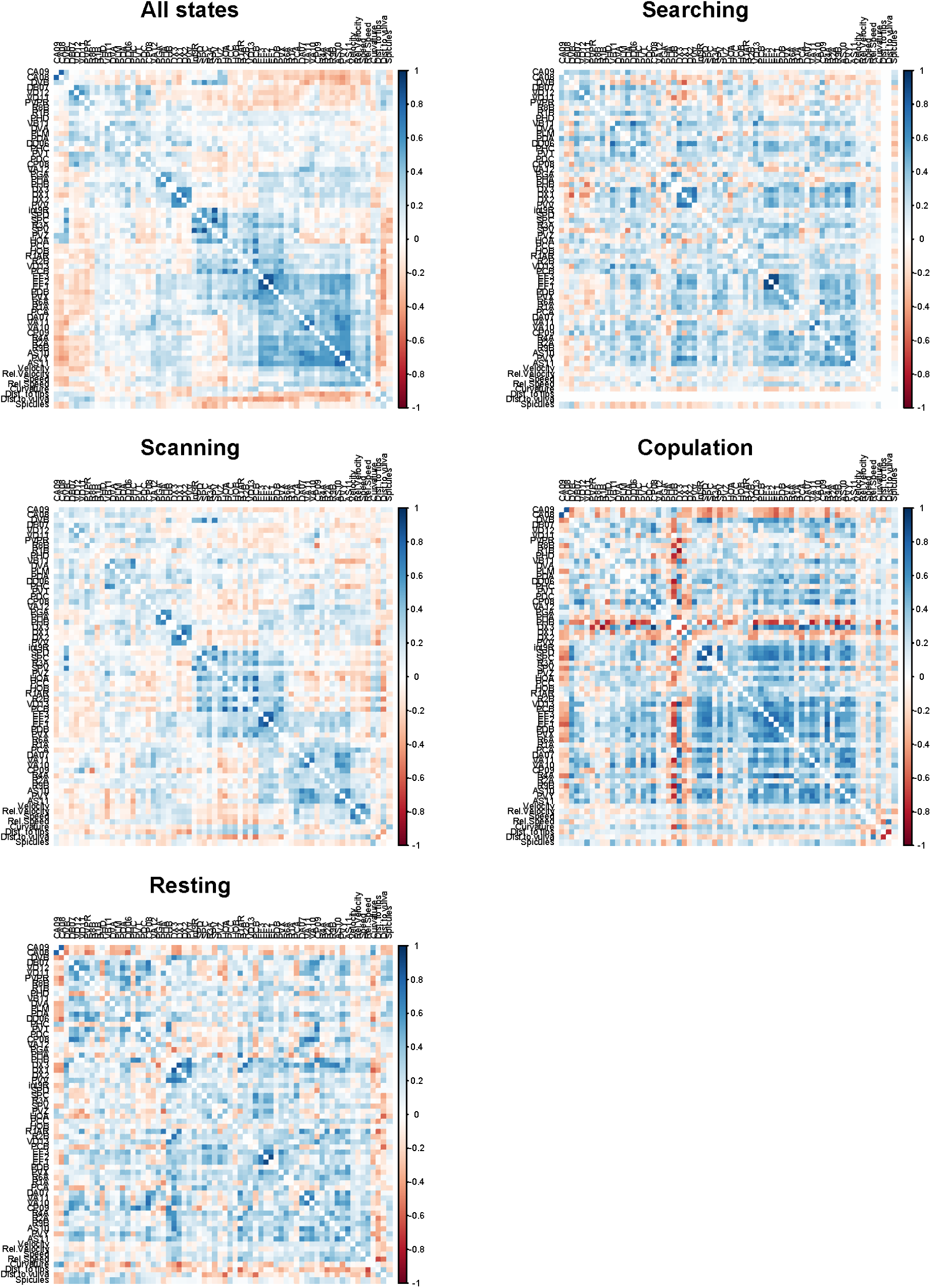
State-specific correlations. Consensus cross-correlation matrices for four major behavioral states: searching, scanning, copulation, and resting.

**Fig. S5.**
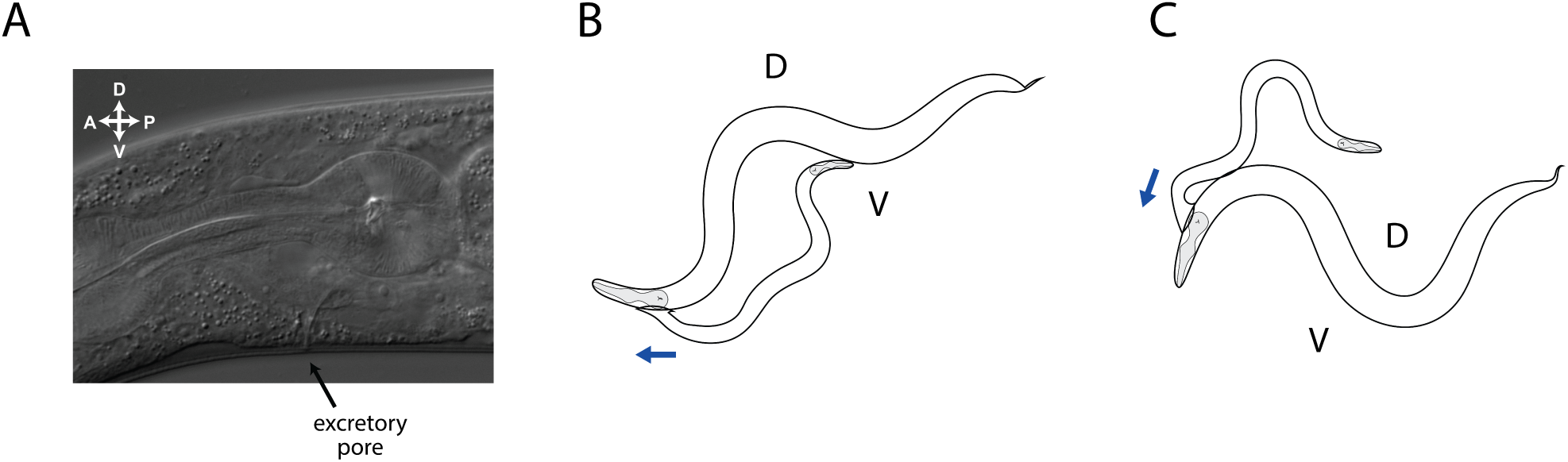
Excretory pore detection. The opening of the excretory pore of the hermaphrodite (**A**) is detected by the male’s phasmid sensory neurons PHA, PHB. We tested whether the male pauses backward scanning near the excretory pore (**B**) more often than in pauses on the corresponding dorsal side (**C**) of the vulvaless *let-23* hermaphrodites.

**Fig. S6.**
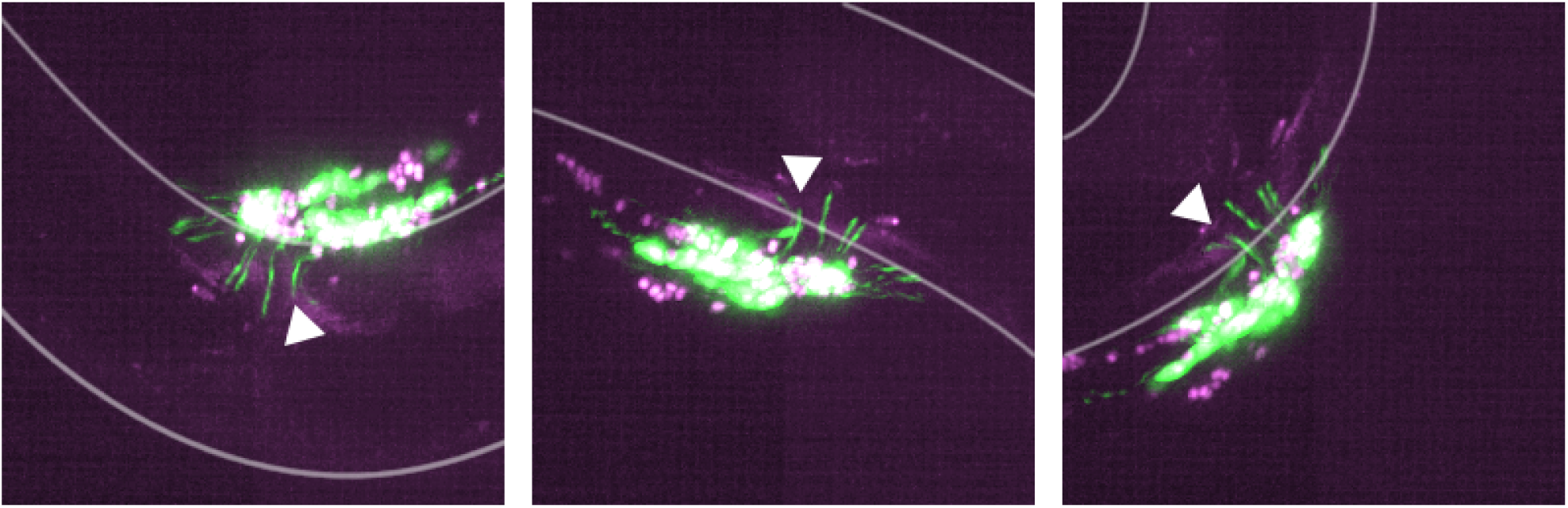
Position of the male rays during vulva detection events. When the male stops at the vulva, some of its rays are positioned opposite of the opening of the vulva. B-type ray neurons expressing pPkd-2::GFP are shown in green. Arrows point at the second ray. Hermaphrodite’s vulva can be seen in magenta. Right lateral aspect. Maximum intensity projections are shown.

**Fig. S7.**
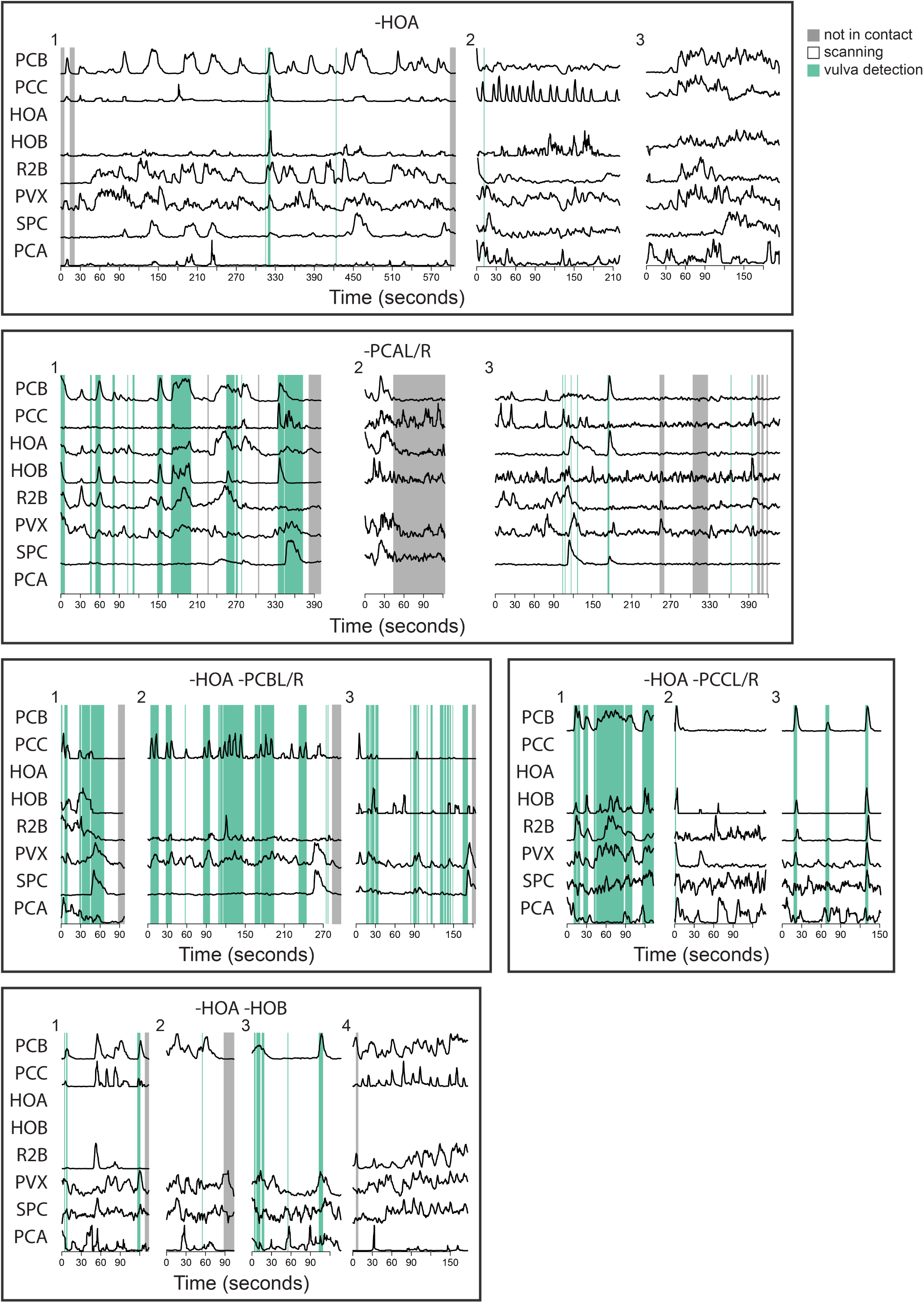
Activities of neurons involved in vulva detection recorded from ablated males during mating. Each neuron or neuron combination was targeted in three to four different males, and the activities of the remaining neurons of interest were recorded.

**Fig. S8.**
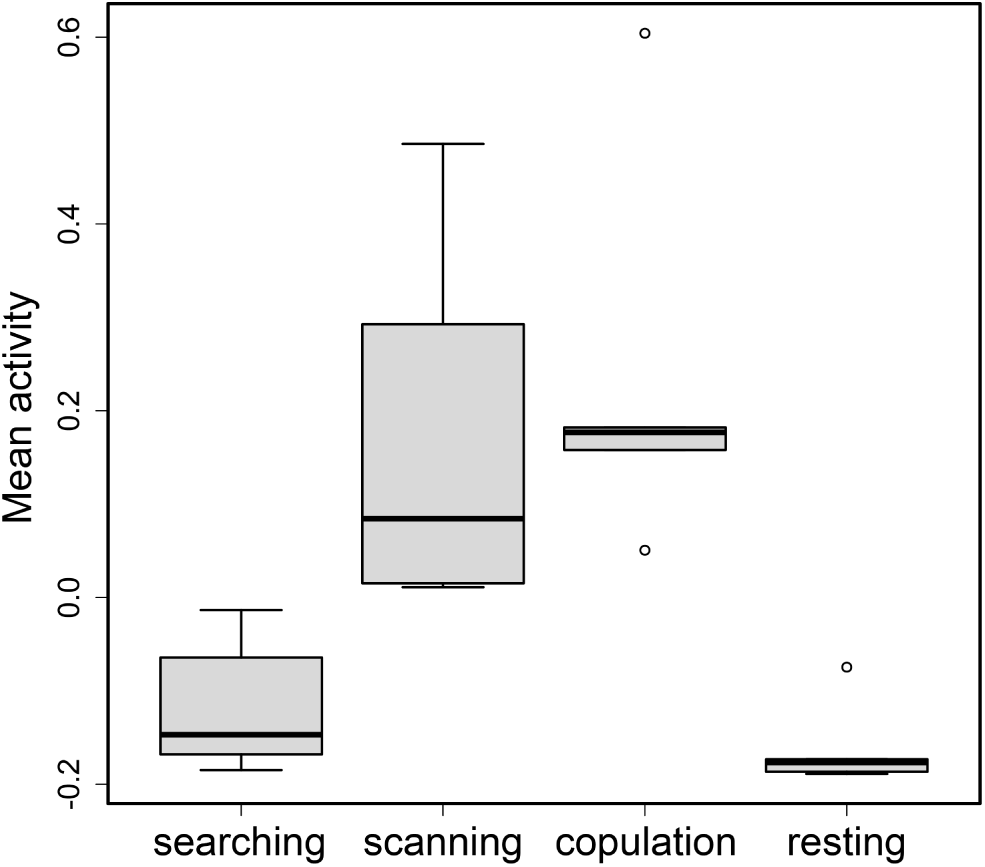
Mean circuit activity for seven males and four different states of mating behavior. Global activity of the circuit was calculated for the four major states of mating behavior – searching, scanning, copulation, and resting using normalized activity traces.

**Fig. S9.**
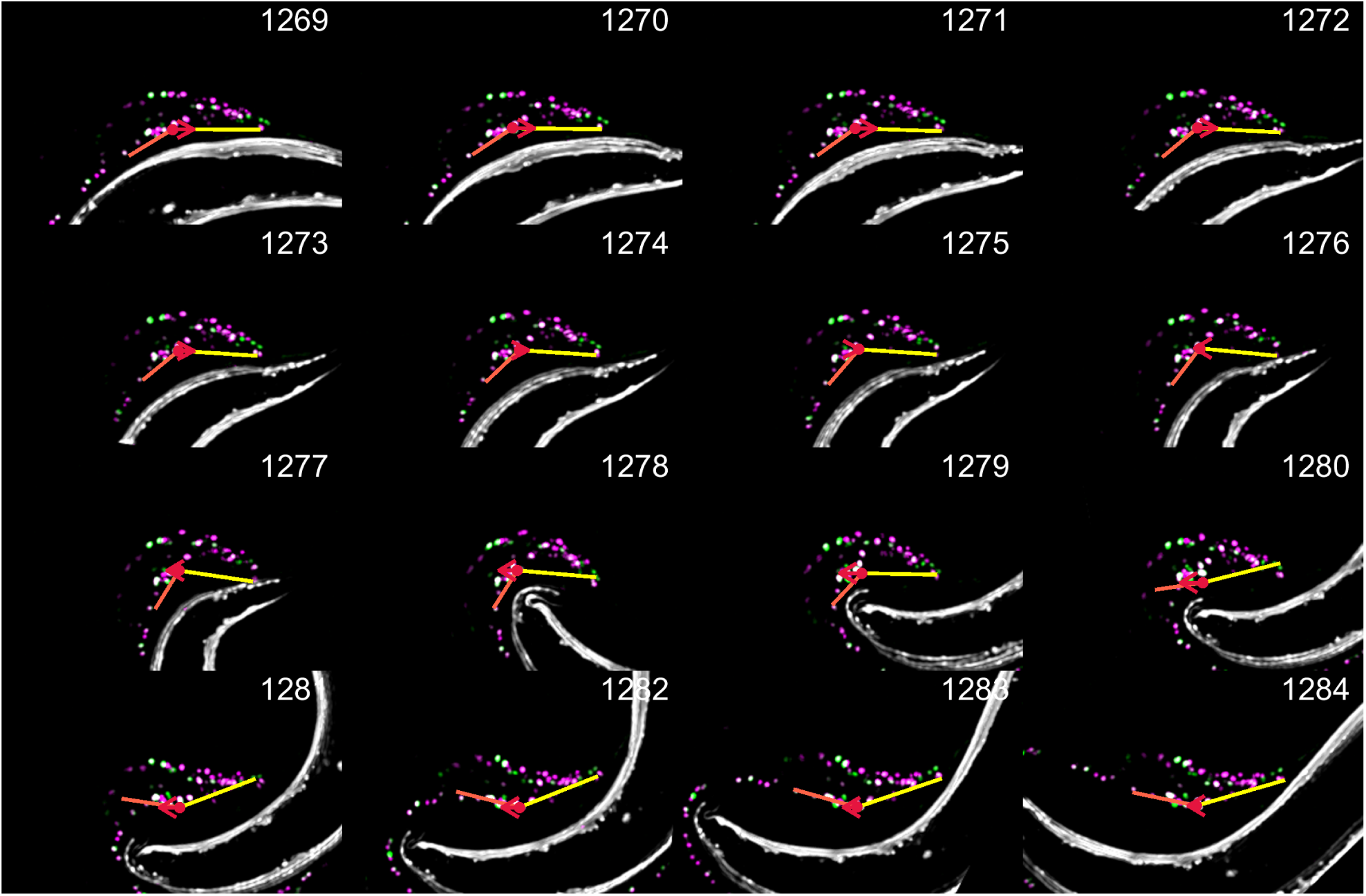
Velocity and tail curvature at turning. A set of continuous features was extracted from the recordings, including velocity (shown with red arrow) and tail curvature (shown with orange and yellow lines).

**Fig. S10.**
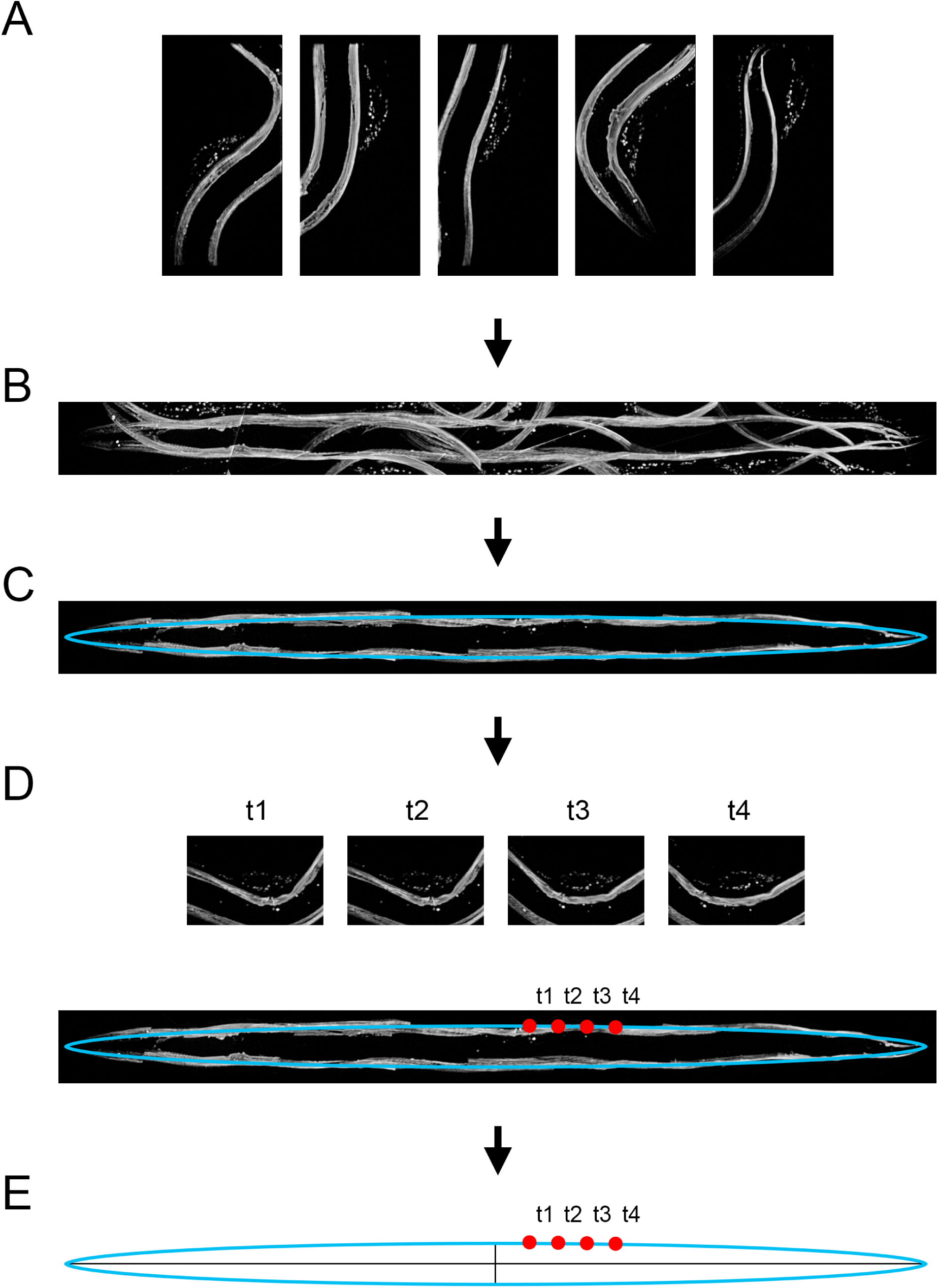
Mapping the tail position on the hermaphrodite. Maximum intensity projections showing large parts of the hermaphrodite were selected (**A**) to cover the entire body of the hermaphrodite. Images were stitched to create a straightened representation of the hermaphrodite (**B**,**C**). For every volume of the recording, the position of the male tail was mapped to the straightened hermaphrodite (**D**) to generate a male tail trajectory on an an idealized elliptical hermaphrodite (**E**)

