## Supplementary material for "Natural sensory context drives diverse brain-wide activity during *C. elegans* mating": S1

#### The PDF file includes:

S1. Tuning properties of individual neurons of *C. elegans* posterior brain.

**S1. Tuning properties of individual neurons of *C. elegans* posterior brain.** For each recorded neuron in the male tail, its activity patterns are shown. **(A)** Neuron activity plotted for an entire mating behavior for a representative dataset. The neuron's location in the posterior brain is shown in red. **(B)** Neuron activities from up to seven males as a function of the tail position on the hermaphrodite (as in **Fig. 5**). Mean activity for all males is shown in blue. **(C)** Neuron activities from all males aligned to the onset of discrete behavioral events. The discrete events include (in the order of appearance) turning, failed turning, vulva contact, sperm release, switching from backward to forward movement (tabulated for the entire behavior), switching from forward to backward movement (tabulated for the entire behavior), switching from backward to forward sliding (relative to the hermaphrodite), switching from forward to backward sliding (relative to the hermaphrodite), switching from backward to forward movement (during searching), switching from forward to backward movement (during searching), switching from backward to forward movement (during scanning), switching from forward to backward movement (during scanning), hermaphrodite contact, hermaphrodite loss, and sperm release (an extended time window is shown). The number of events observed for each neuron is in parenthesis. Red lines show mean activities for individual males. The mean activity for all males is in black and standard deviation is indicated in gray.

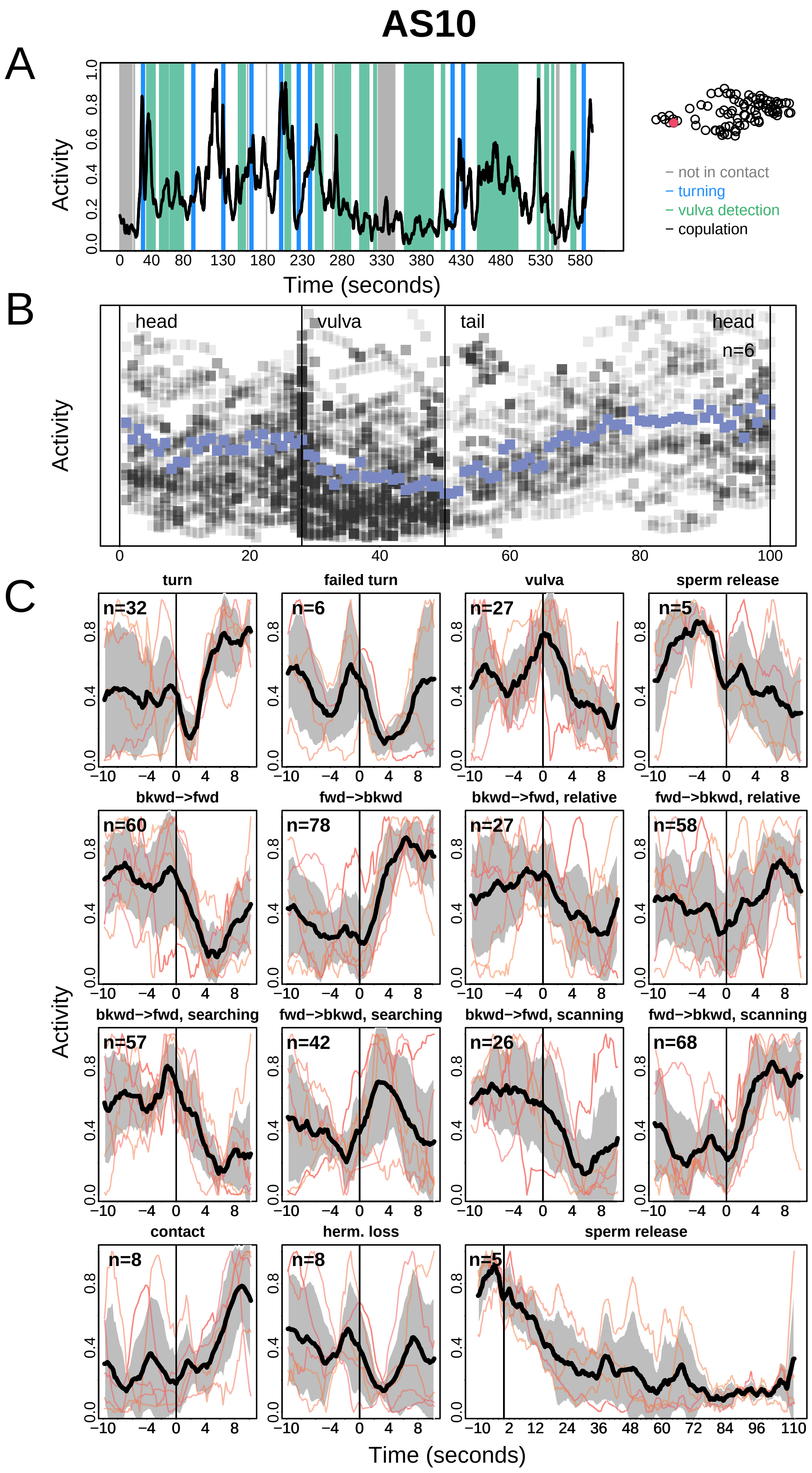
AS10
A
Activity
Time (seconds)
− not in contact
− turning
− vulva detection
− copulation
B
head
vulva
tail
head
n=6
Activity
C
turn
failed turn
vulva
sperm release
bkwd−>fwd
fwd−>bkwd
bkwd−>fwd, relative
fwd−>bkwd, relative
bkwd−>fwd, searching
fwd−>bkwd, searching
bkwd−>fwd, scanning
fwd−>bkwd, scanning
contact
herm. loss
sperm release
Activity
Time (seconds)

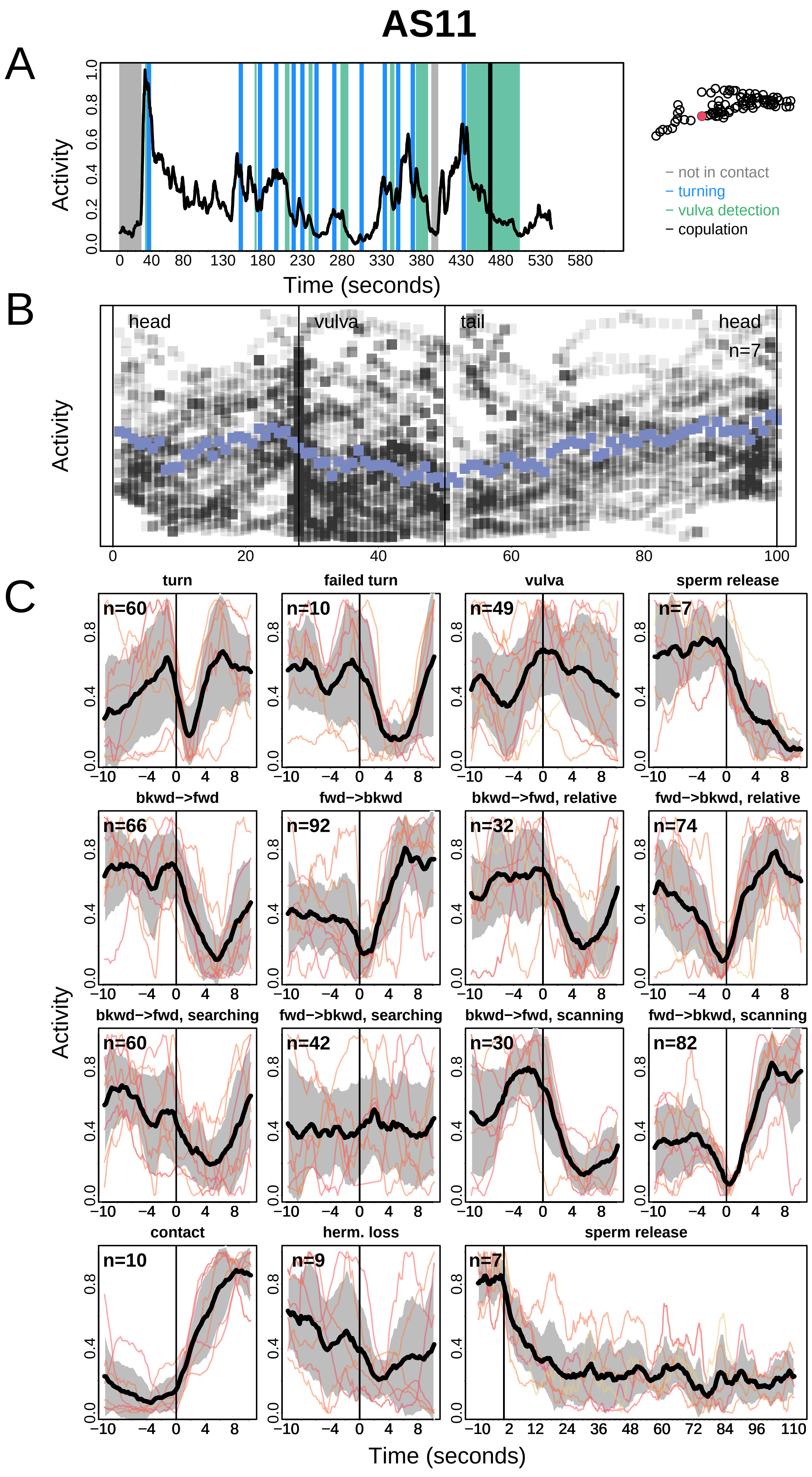
AS11
A
Activity
Time (seconds)
– not in contact
– turning
– vulva detection
– copulation
B
head
vulva
tail
head
n=7
Activity
C
turn
failed turn
vulva
sperm release
bkwd−>fwd
fwd−>bkwd
bkwd−>fwd, relative
fwd−>bkwd, relative
bkwd−>fwd, searching
fwd−>bkwd, searching
bkwd−>fwd, scanning
fwd−>bkwd, scanning
contact
herm. loss
sperm release
Activity
Time (seconds)

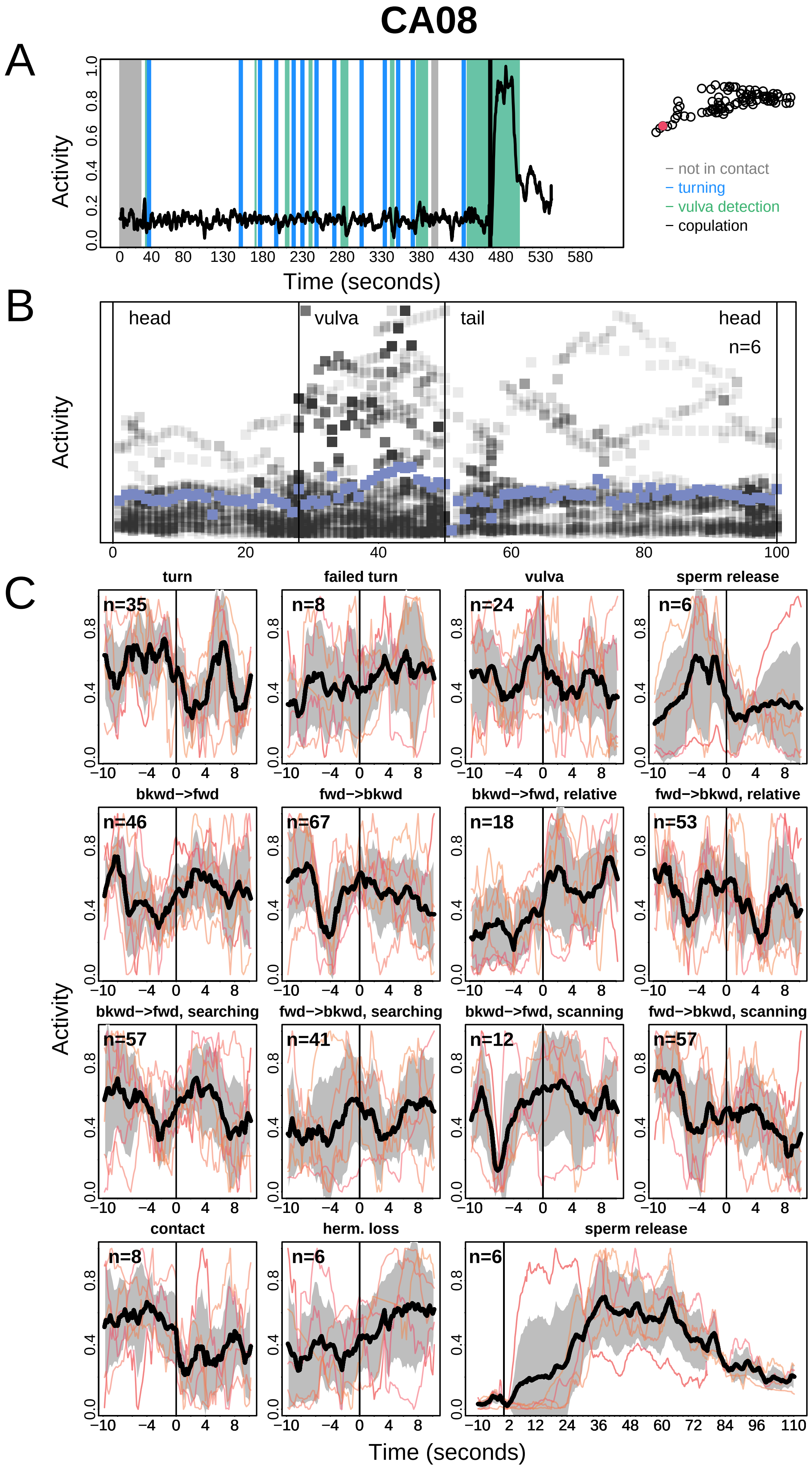
CA08
A
Activity
Time (seconds)
– not in contact
– turning
– vulva detection
– copulation
B
head
vulva
tail
head
n=6
Activity
C
turn
failed turn
vulva
sperm release
bkwd−>fwd
fwd−>bkwd
bkwd−>fwd, relative
fwd−>bkwd, relative
bkwd−>fwd, searching
fwd−>bkwd, searching
bkwd−>fwd, scanning
fwd−>bkwd, scanning
contact
herm. loss
sperm release
Activity
Time (seconds)

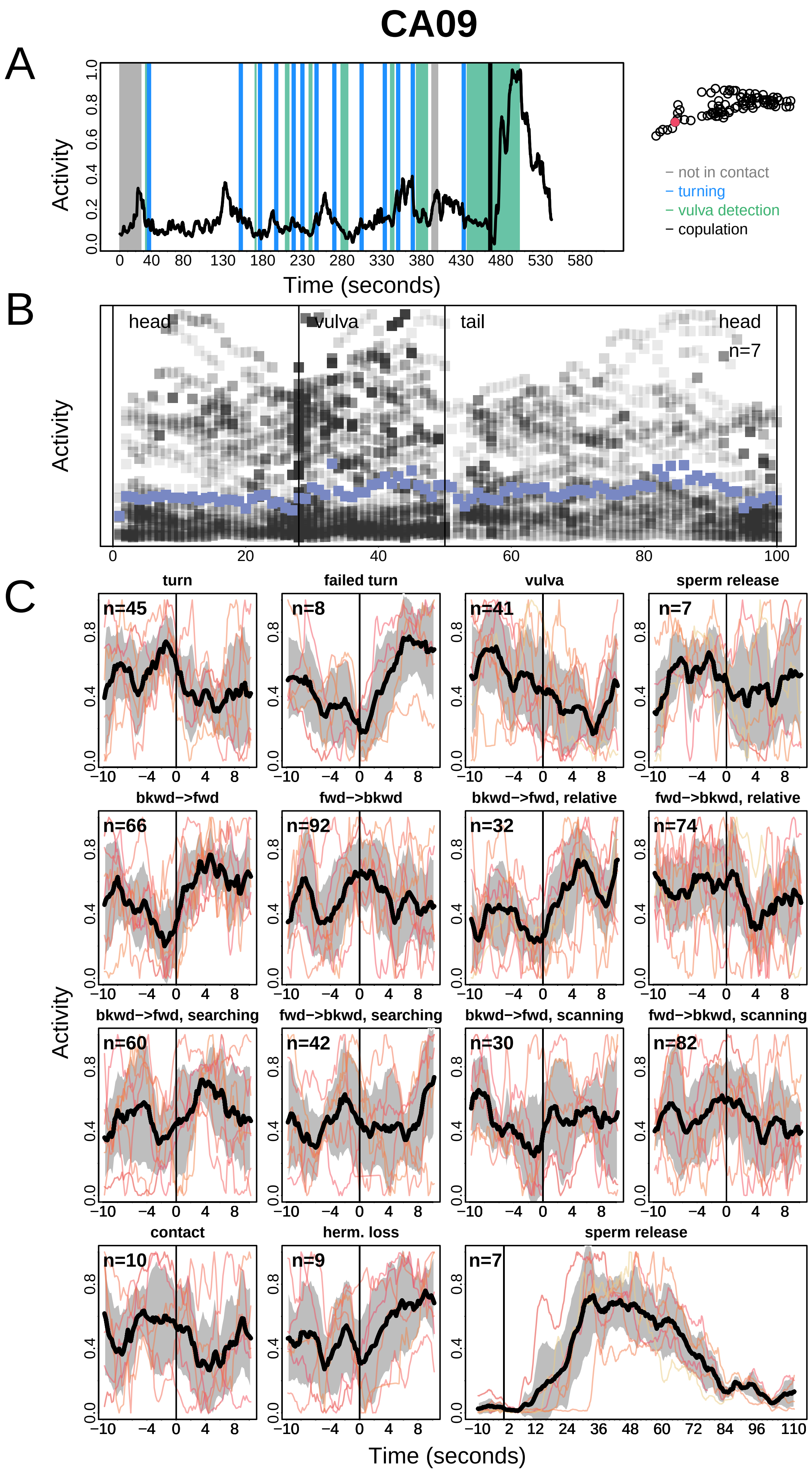
CA09
A
Activity
Time (seconds)
not in contact
turning
vulva detection
copulation
B
head
vulva
tail
head
n=7
C
turn
failed turn
vulva
sperm release
bkwd−>fwd
fwd−>bkwd
bkwd−>fwd, relative
fwd−>bkwd, relative
bkwd−>fwd, searching
fwd−>bkwd, searching
bkwd−>fwd, scanning
fwd−>bkwd, scanning
contact
herm. loss
sperm release
n=45
n=8
n=41
n=7
n=66
n=92
n=32
n=74
n=60
n=42
n=30
n=82
n=10
n=9
n=7
Time (seconds)

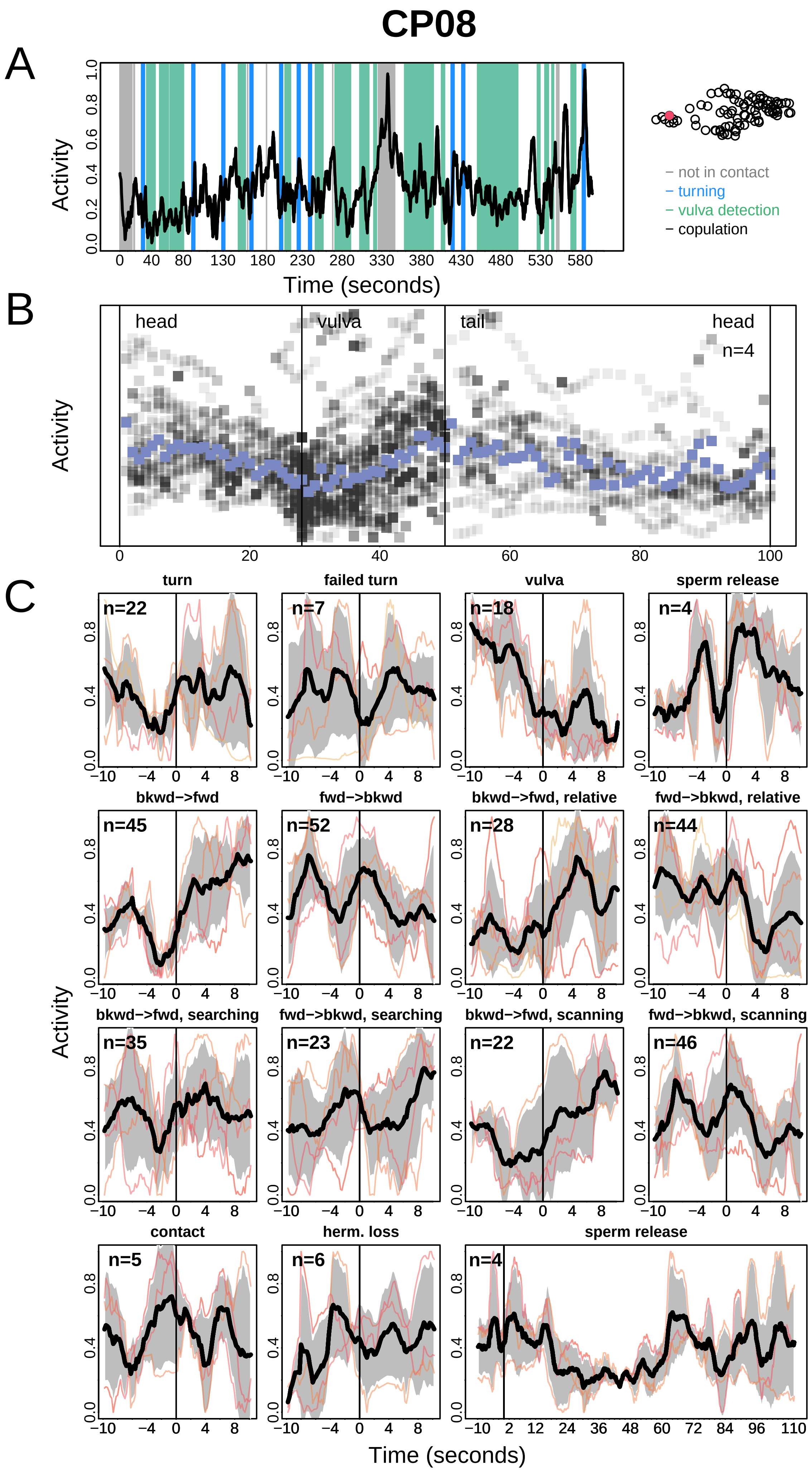
CP08
A
Activity
Time (seconds)
− not in contact
− turning
− vulva detection
− copulation
B
head
vulva
tail
head
n=4
C
turn
failed turn
vulva
sperm release
bkwd−>fwd
fwd−>bkwd
bkwd−>fwd, relative
fwd−>bkwd, relative
bkwd−>fwd, searching
fwd−>bkwd, searching
bkwd−>fwd, scanning
fwd−>bkwd, scanning
contact
herm. loss
sperm release
n=22
n=7
n=18
n=4
n=45
n=52
n=28
n=44
n=35
n=23
n=22
n=46
n=5
n=6
n=4
Time (seconds)

# CP09

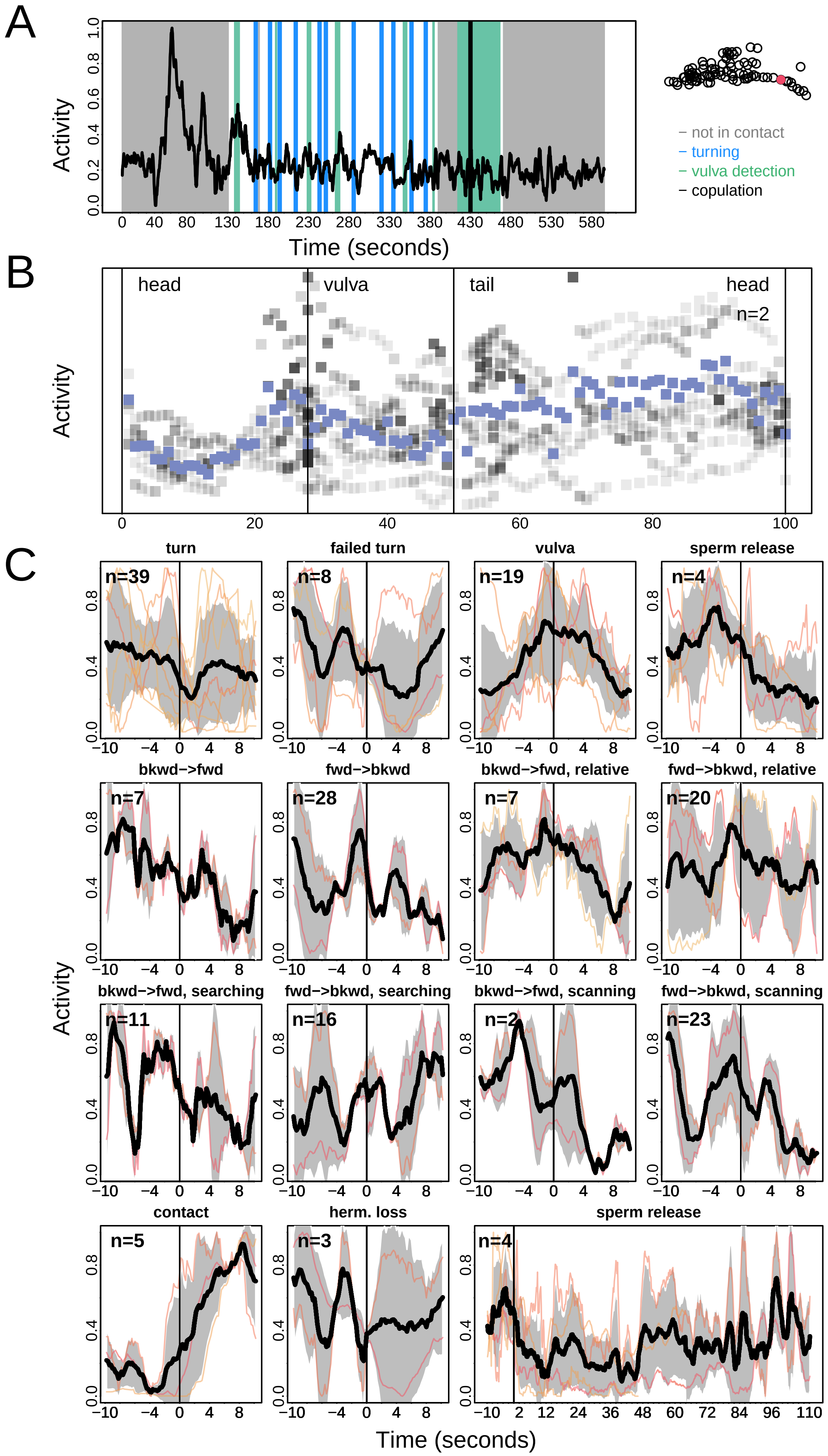
A
Activity
Time (seconds)
– not in contact
– turning
– vulva detection
– copulation
B
head
vulva
tail
head
n=2
Activity
C
turn
failed turn
vulva
sperm release
bkwd−>fwd
fwd−>bkwd
bkwd−>fwd, relative
fwd−>bkwd, relative
bkwd−>fwd, searching
fwd−>bkwd, searching
bkwd−>fwd, scanning
fwd−>bkwd, scanning
contact
herm. loss
sperm release
n=39
n=8
n=19
n=4
n=7
n=28
n=7
n=20
n=11
n=16
n=2
n=23
n=5
n=3
n=4
Activity
Time (seconds)

# DA07

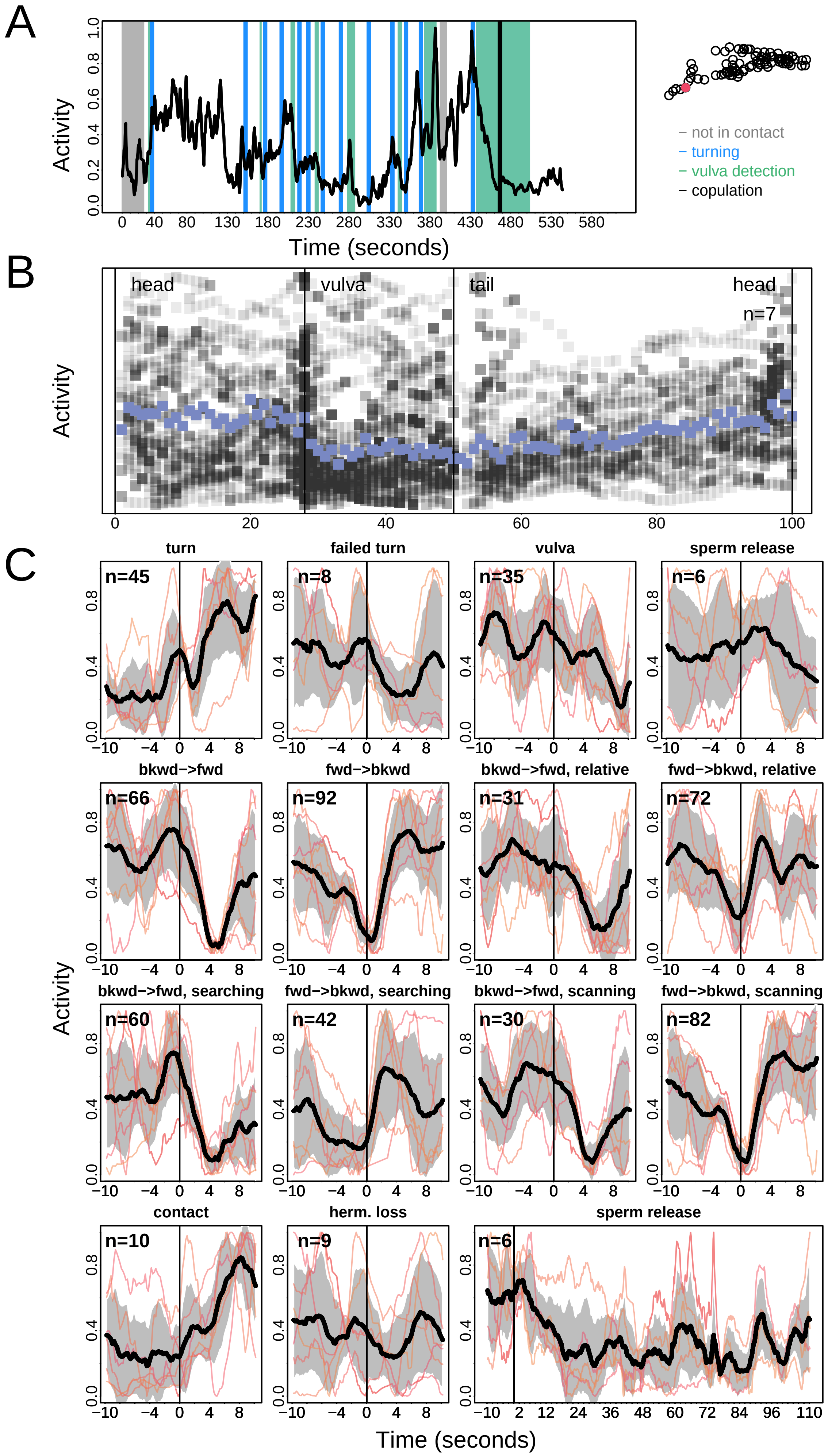

A
Activity
Time (seconds)
not in contact
turning
vulva detection
copulation
B
head
vulva
tail
head
n=7
Activity
C
turn
failed turn
vulva
sperm release
bkwd−>fwd
fwd−>bkwd
bkwd−>fwd, relative
fwd−>bkwd, relative
bkwd−>fwd, searching
fwd−>bkwd, searching
bkwd−>fwd, scanning
fwd−>bkwd, scanning
contact
herm. loss
sperm release
n=45
n=8
n=35
n=6
n=66
n=92
n=31
n=72
n=60
n=42
n=30
n=82
n=10
n=9
n=6
Activity
Time (seconds)

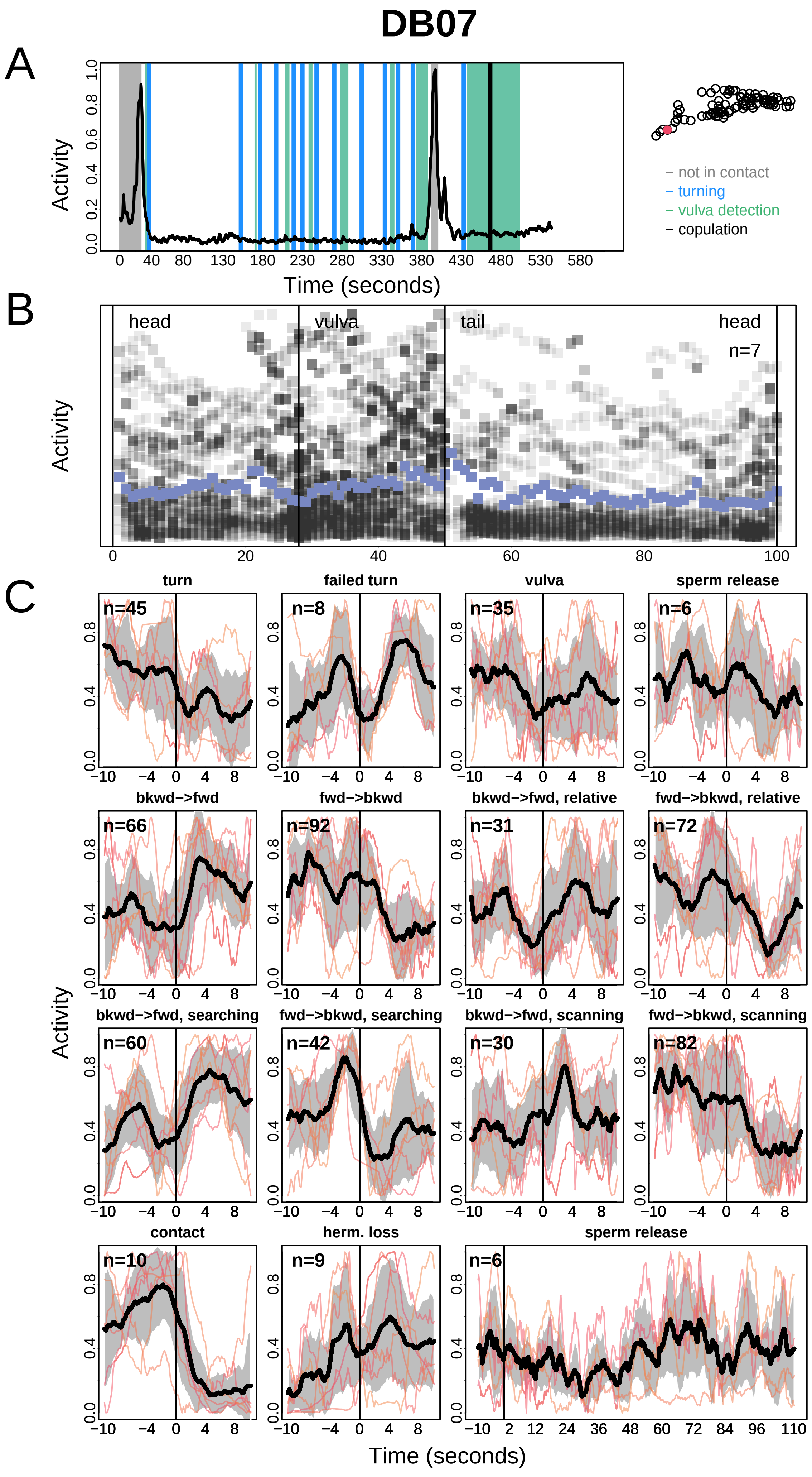
DB07
A
Activity
Time (seconds)
− not in contact
− turning
− vulva detection
− copulation
B
head
vulva
tail
head
n=7
Activity
C
turn
failed turn
vulva
sperm release
bkwd−>fwd
fwd−>bkwd
bkwd−>fwd, relative
fwd−>bkwd, relative
bkwd−>fwd, searching
fwd−>bkwd, searching
bkwd−>fwd, scanning
fwd−>bkwd, scanning
contact
herm. loss
sperm release
Activity
Time (seconds)

# DD06

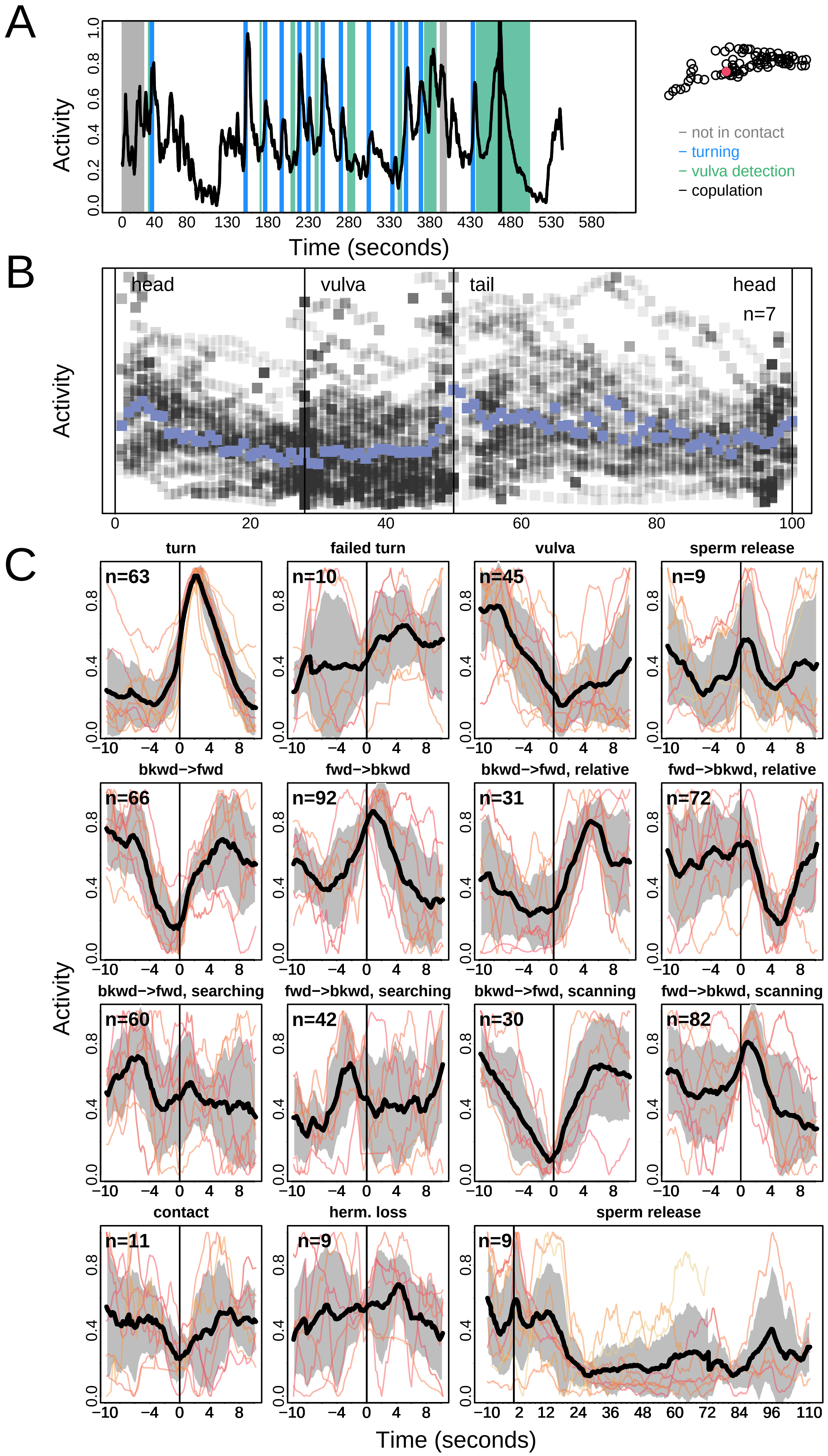
A
Activity
Time (seconds)
– not in contact
– turning
– vulva detection
– copulation
B
head
vulva
tail
head
n=7
Activity
C
turn
failed turn
vulva
sperm release
bkwd−>fwd
fwd−>bkwd
bkwd−>fwd, relative
fwd−>bkwd, relative
bkwd−>fwd, searching
fwd−>bkwd, searching
bkwd−>fwd, scanning
fwd−>bkwd, scanning
contact
herm. loss
sperm release
Activity
Time (seconds)

DVA

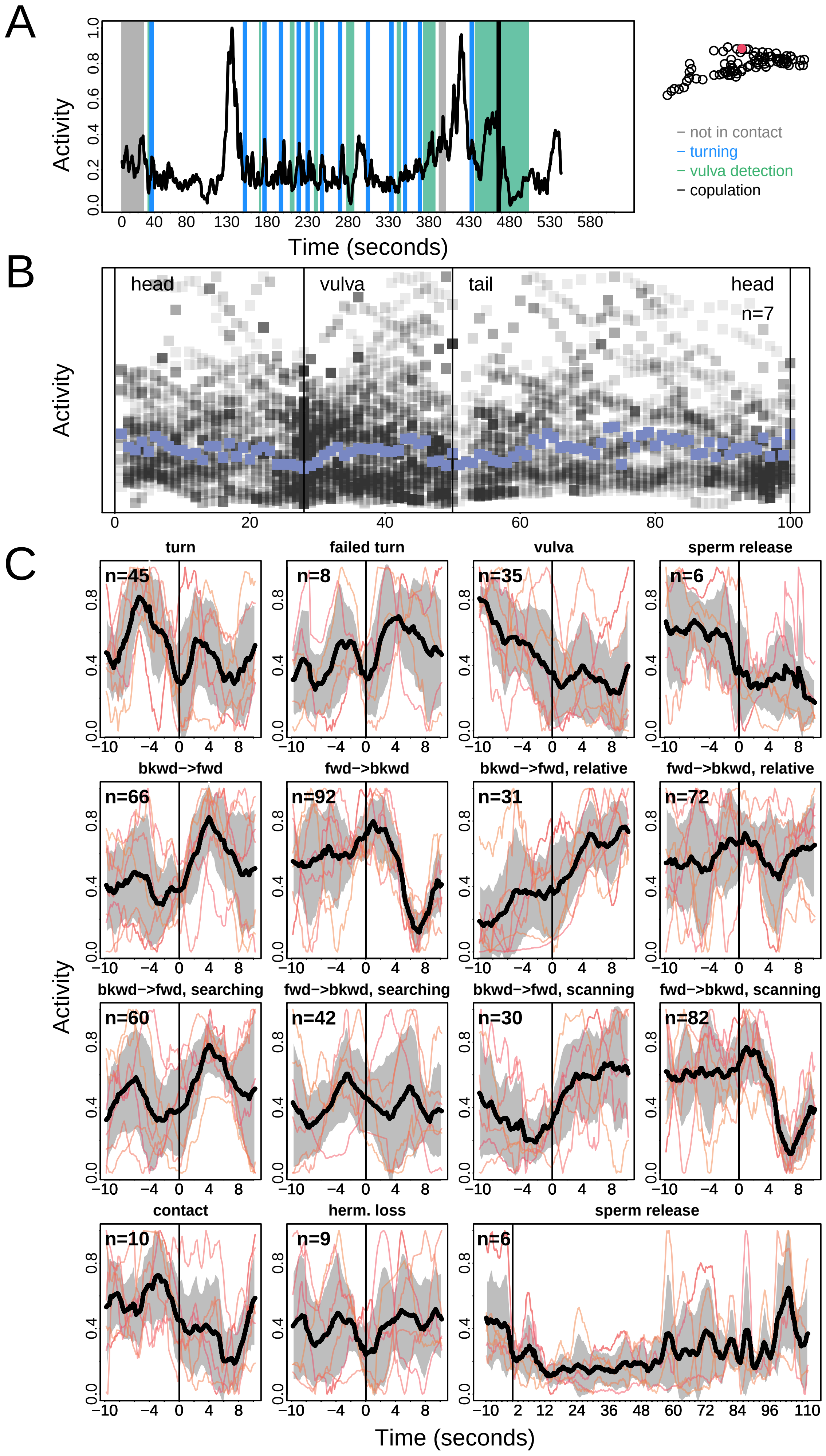
A
Activity
Time (seconds)
– not in contact
– turning
– vulva detection
– copulation
B
head
vulva
tail
head
n=7
Activity
C
turn
failed turn
vulva
sperm release
bkwd–>fwd
fwd–>bkwd
bkwd–>fwd, relative
fwd–>bkwd, relative
bkwd–>fwd, searching
fwd–>bkwd, searching
bkwd–>fwd, scanning
fwd–>bkwd, scanning
contact
herm. loss
sperm release
n=45
n=8
n=35
n=6
n=66
n=92
n=31
n=72
n=60
n=42
n=30
n=82
n=10
n=9
n=6
Activity
Time (seconds)

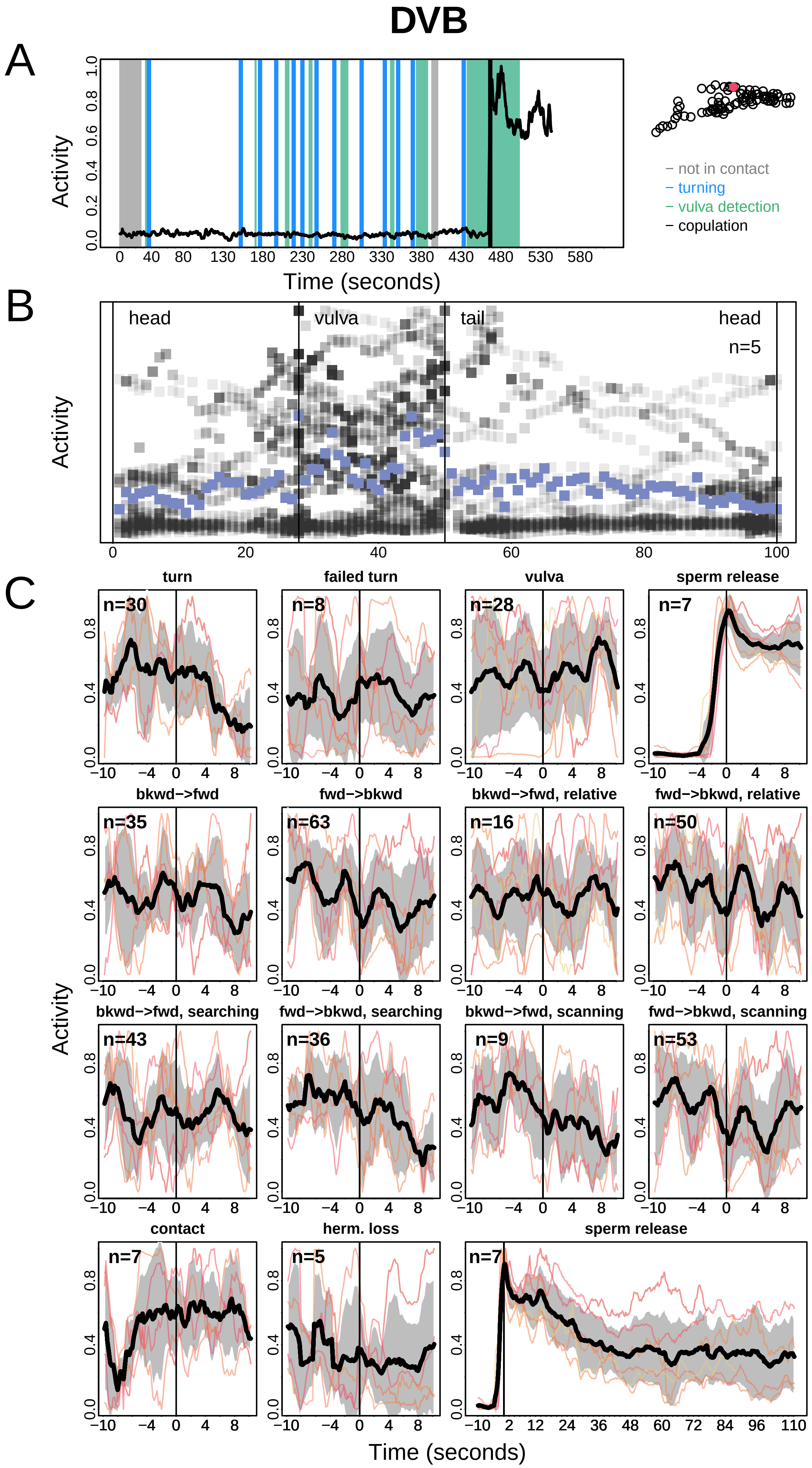

DVB
A
Activity
Time (seconds)
– not in contact
– turning
– vulva detection
– copulation
B
head
vulva
tail
head
n=5
Activity
C
turn
failed turn
vulva
sperm release
bkwd–>fwd
fwd–>bkwd
bkwd–>fwd, relative
fwd–>bkwd, relative
bkwd–>fwd, searching
fwd–>bkwd, searching
bkwd–>fwd, scanning
fwd–>bkwd, scanning
contact
herm. loss
sperm release
Activity
Time (seconds)

# DX1

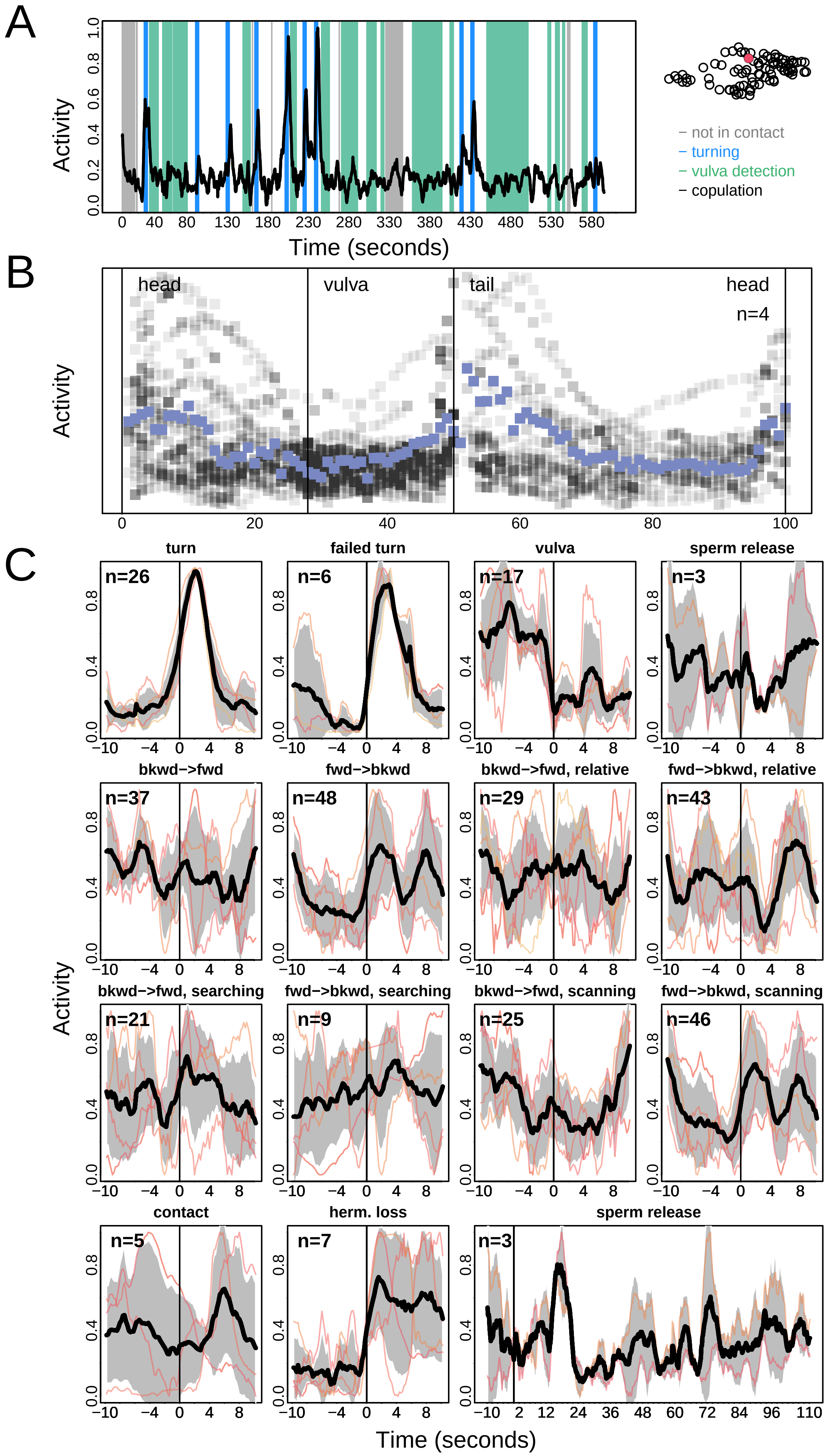
A
B
C
Activity
Time (seconds)
not in contact
turning
vulva detection
copulation
head
vulva
tail
head
n=4
turn
failed turn
vulva
sperm release
bkwd->fwd
fwd->bkwd
bkwd->fwd, relative
fwd->bkwd, relative
bkwd->fwd, searching
fwd->bkwd, searching
bkwd->fwd, scanning
fwd->bkwd, scanning
contact
herm. loss
sperm release
n=26
n=6
n=17
n=3
n=37
n=48
n=29
n=43
n=21
n=9
n=25
n=46
n=5
n=7
n=3

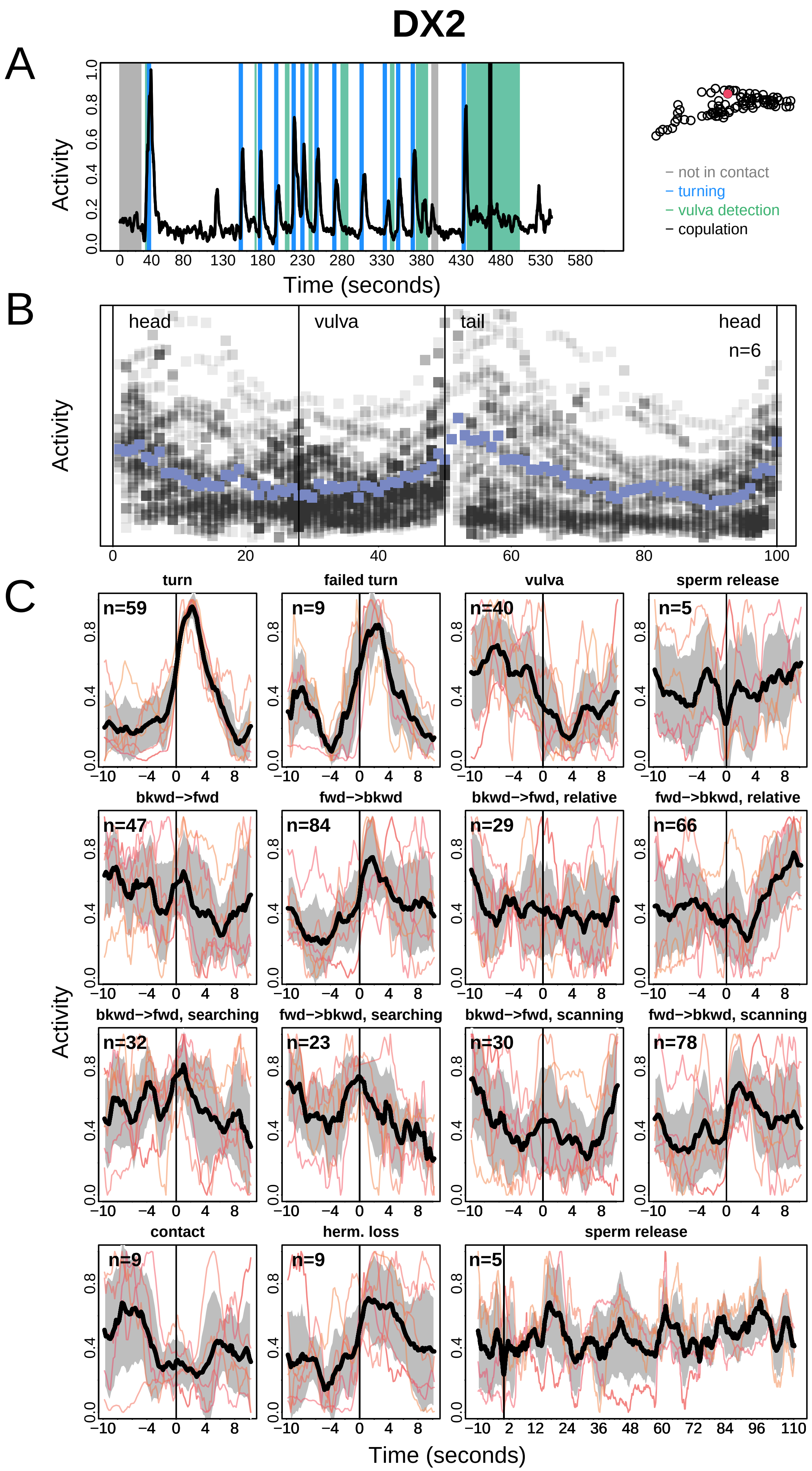
DX2
A
Activity
Time (seconds)
− not in contact
− turning
− vulva detection
− copulation
B
head
vulva
tail
head
n=6
Activity
C
turn
failed turn
vulva
sperm release
bkwd−>fwd
fwd−>bkwd
bkwd−>fwd, relative
fwd−>bkwd, relative
bkwd−>fwd, searching
fwd−>bkwd, searching
bkwd−>fwd, scanning
fwd−>bkwd, scanning
contact
herm. loss
sperm release
n=59
n=9
n=40
n=5
n=47
n=84
n=29
n=66
n=32
n=23
n=30
n=78
n=9
n=9
n=5
Activity
Time (seconds)

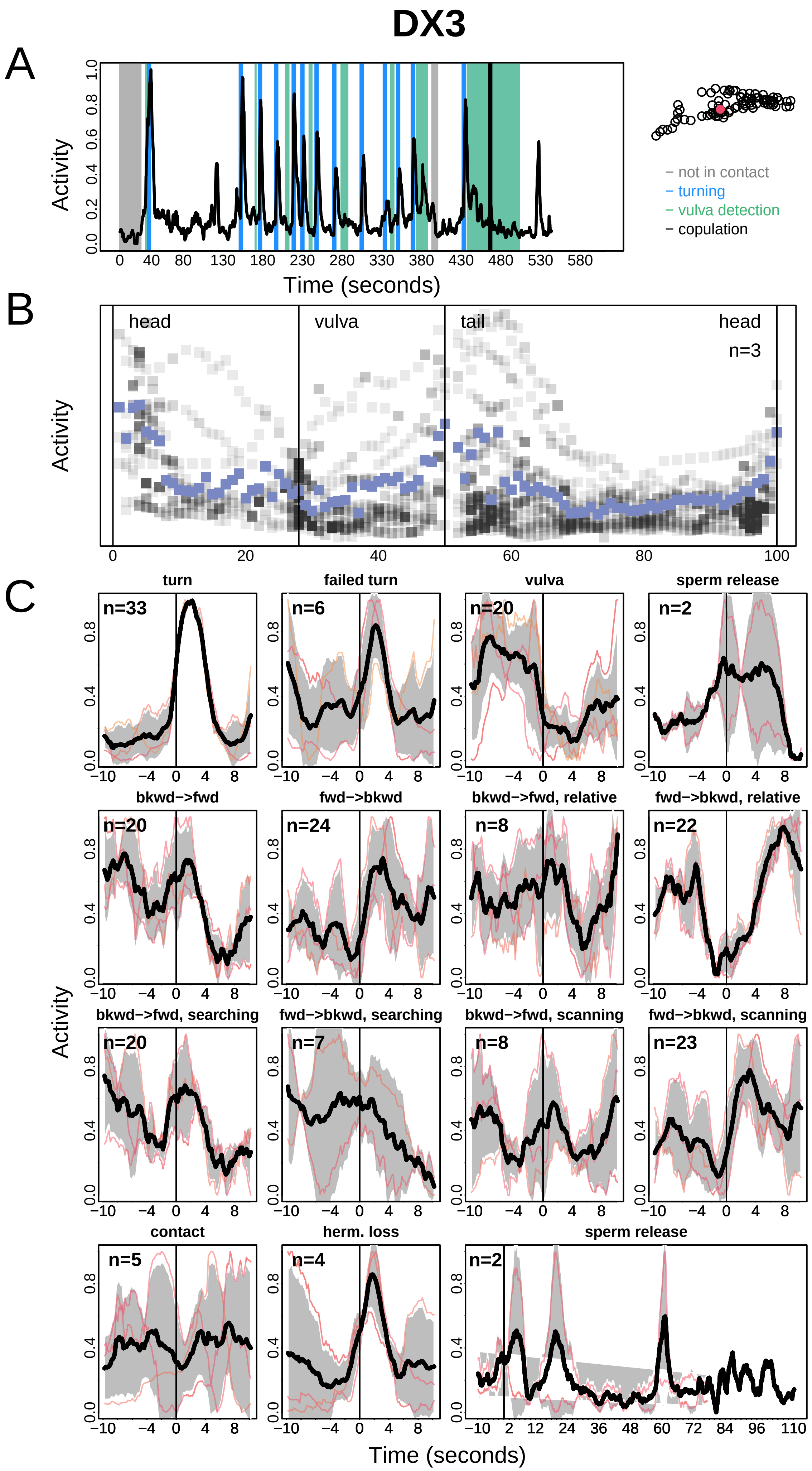

DX3
A
Activity
Time (seconds)
0 40 80 130 180 230 280 330 380 430 480 530 580
0.0 0.2 0.4 0.6 0.8 1.0
– not in contact
– turning
– vulva detection
– copulation
B
head
vulva
tail
head
n=3
Activity
0 20 40 60 80 100
C
turn
n=33
failed turn
n=6
vulva
n=20
sperm release
n=2
bkwd−>fwd
n=20
fwd−>bkwd
n=24
bkwd−>fwd, relative
n=8
fwd−>bkwd, relative
n=22
bkwd−>fwd, searching
n=20
fwd−>bkwd, searching
n=7
bkwd−>fwd, scanning
n=8
fwd−>bkwd, scanning
n=23
contact
n=5
herm. loss
n=4
sperm release
n=2
Activity
−10 −4 0 4 8
−10 2 12 24 36 48 60 72 84 96 110
Time (seconds)

# EF1

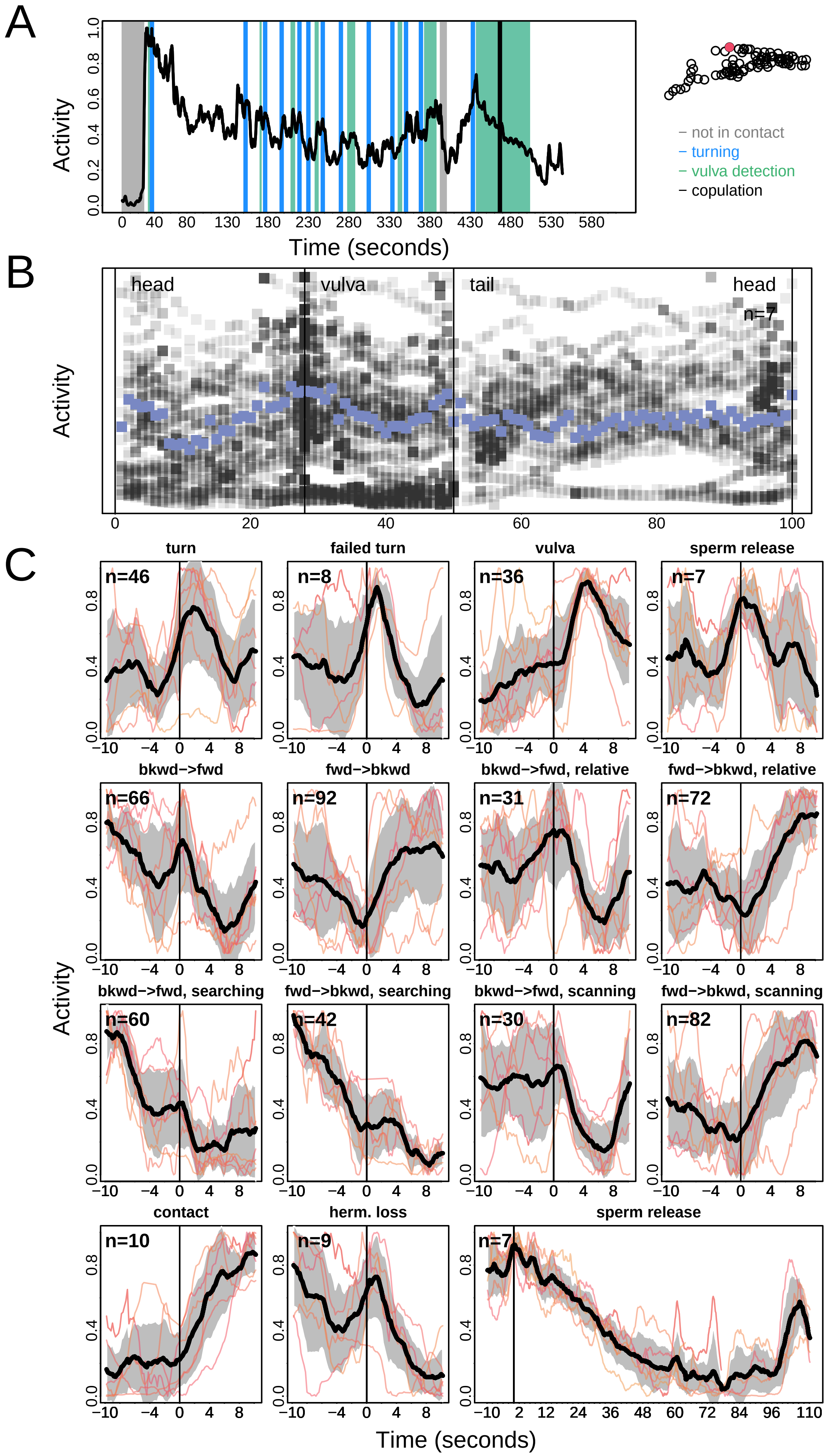
A
Activity
Time (seconds)
not in contact
turning
vulva detection
copulation
B
head
vulva
tail
head
n=7
Activity
C
turn
failed turn
vulva
sperm release
bkwd−>fwd
fwd−>bkwd
bkwd−>fwd, relative
fwd−>bkwd, relative
bkwd−>fwd, searching
fwd−>bkwd, searching
bkwd−>fwd, scanning
fwd−>bkwd, scanning
contact
herm. loss
sperm release
Activity
Time (seconds)

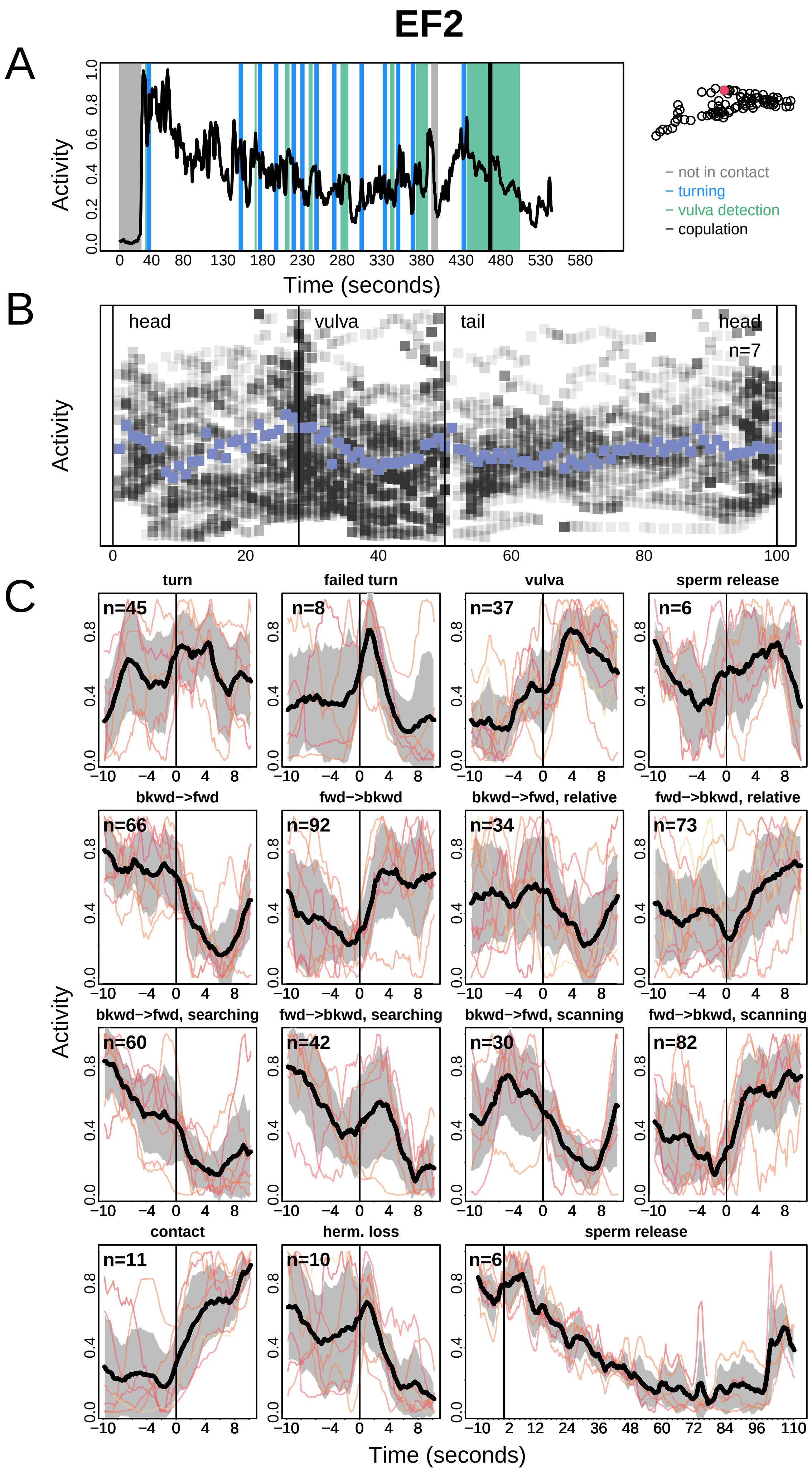

EF2
A
Activity
Time (seconds)
not in contact
turning
vulva detection
copulation
B
head
vulva
tail
head
n=7
Activity
C
turn
failed turn
vulva
sperm release
bkwd−>fwd
fwd−>bkwd
bkwd−>fwd, relative
fwd−>bkwd, relative
bkwd−>fwd, searching
fwd−>bkwd, searching
bkwd−>fwd, scanning
fwd−>bkwd, scanning
contact
herm. loss
sperm release
n=45
n=8
n=37
n=6
n=66
n=92
n=34
n=73
n=60
n=42
n=30
n=82
n=11
n=10
n=6
Activity
Time (seconds)

# EF3

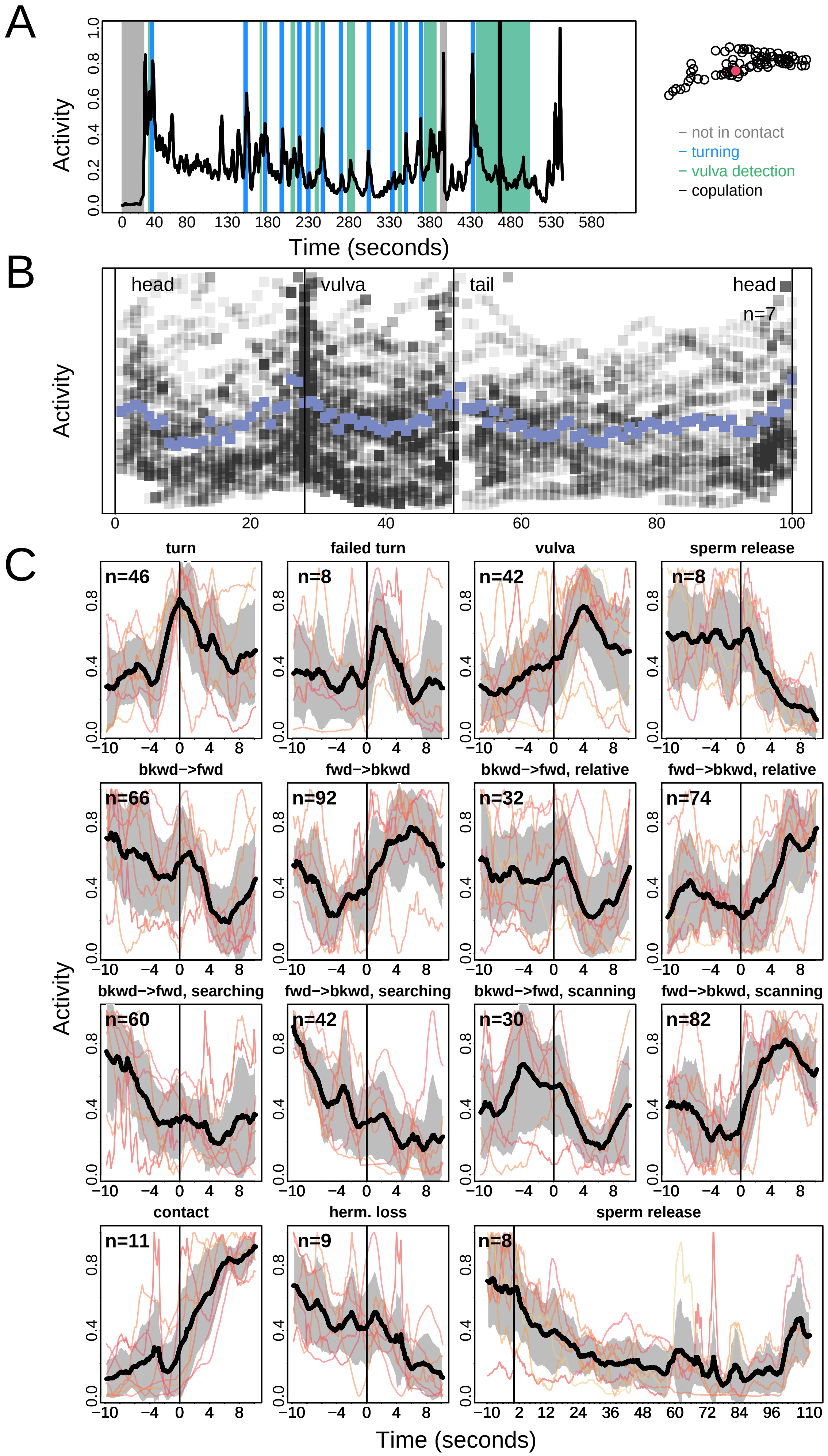
A
Activity
Time (seconds)
– not in contact
– turning
– vulva detection
– copulation
B
head
vulva
tail
head
n=7
Activity
C
turn
n=46
failed turn
n=8
vulva
n=42
sperm release
n=8
bkwd−>fwd
n=66
fwd−>bkwd
n=92
bkwd−>fwd, relative
n=32
fwd−>bkwd, relative
n=74
bkwd−>fwd, searching
n=60
fwd−>bkwd, searching
n=42
bkwd−>fwd, scanning
n=30
fwd−>bkwd, scanning
n=82
contact
n=11
herm. loss
n=9
sperm release
n=8
Activity
Time (seconds)

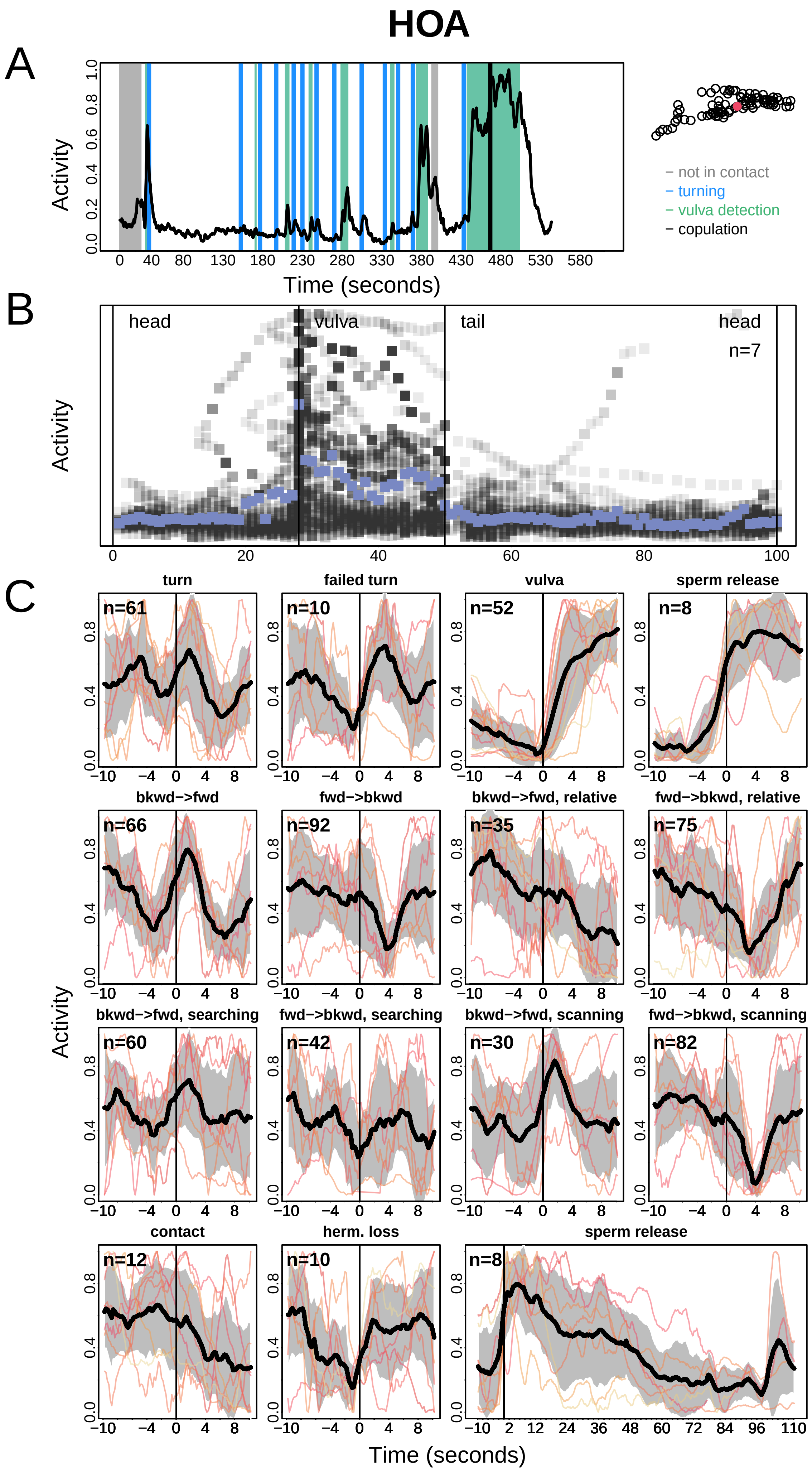
HOA
A
B
C
Activity
Time (seconds)
– not in contact
– turning
– vulva detection
– copulation
head
vulva
tail
head
n=7
turn
failed turn
vulva
sperm release
bkwd–>fwd
fwd–>bkwd
bkwd–>fwd, relative
fwd–>bkwd, relative
bkwd–>fwd, searching
fwd–>bkwd, searching
bkwd–>fwd, scanning
fwd–>bkwd, scanning
contact
herm. loss
sperm release
n=61
n=10
n=52
n=8
n=66
n=92
n=35
n=75
n=60
n=42
n=30
n=82
n=12
n=10
n=8

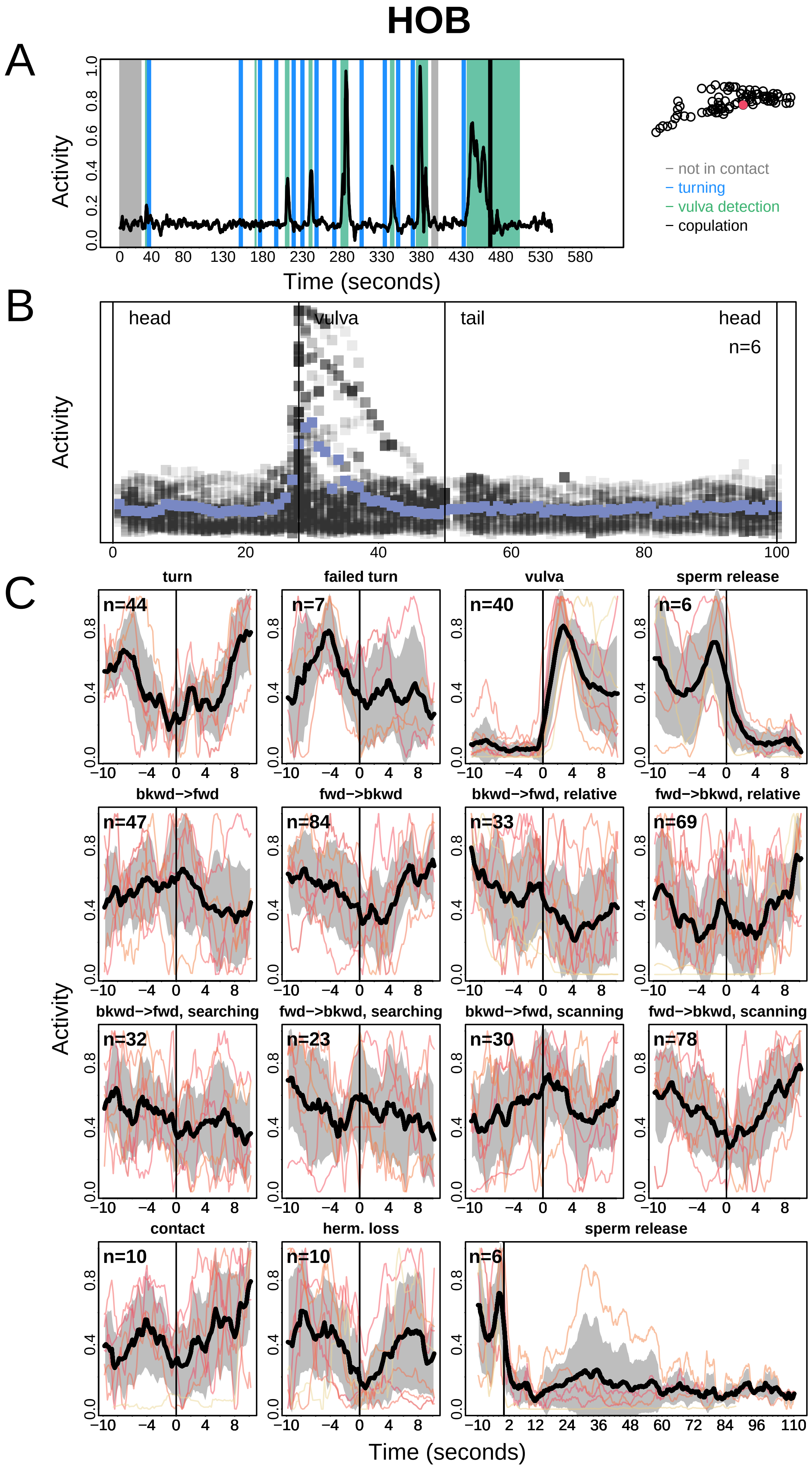
HOB
A
B
C
Activity
Time (seconds)
not in contact
turning
vulva detection
copulation
head
vulva
tail
head
n=6
turn
failed turn
vulva
sperm release
bkwd−>fwd
fwd−>bkwd
bkwd−>fwd, relative
fwd−>bkwd, relative
bkwd−>fwd, searching
fwd−>bkwd, searching
bkwd−>fwd, scanning
fwd−>bkwd, scanning
contact
herm. loss
sperm release

### int9R

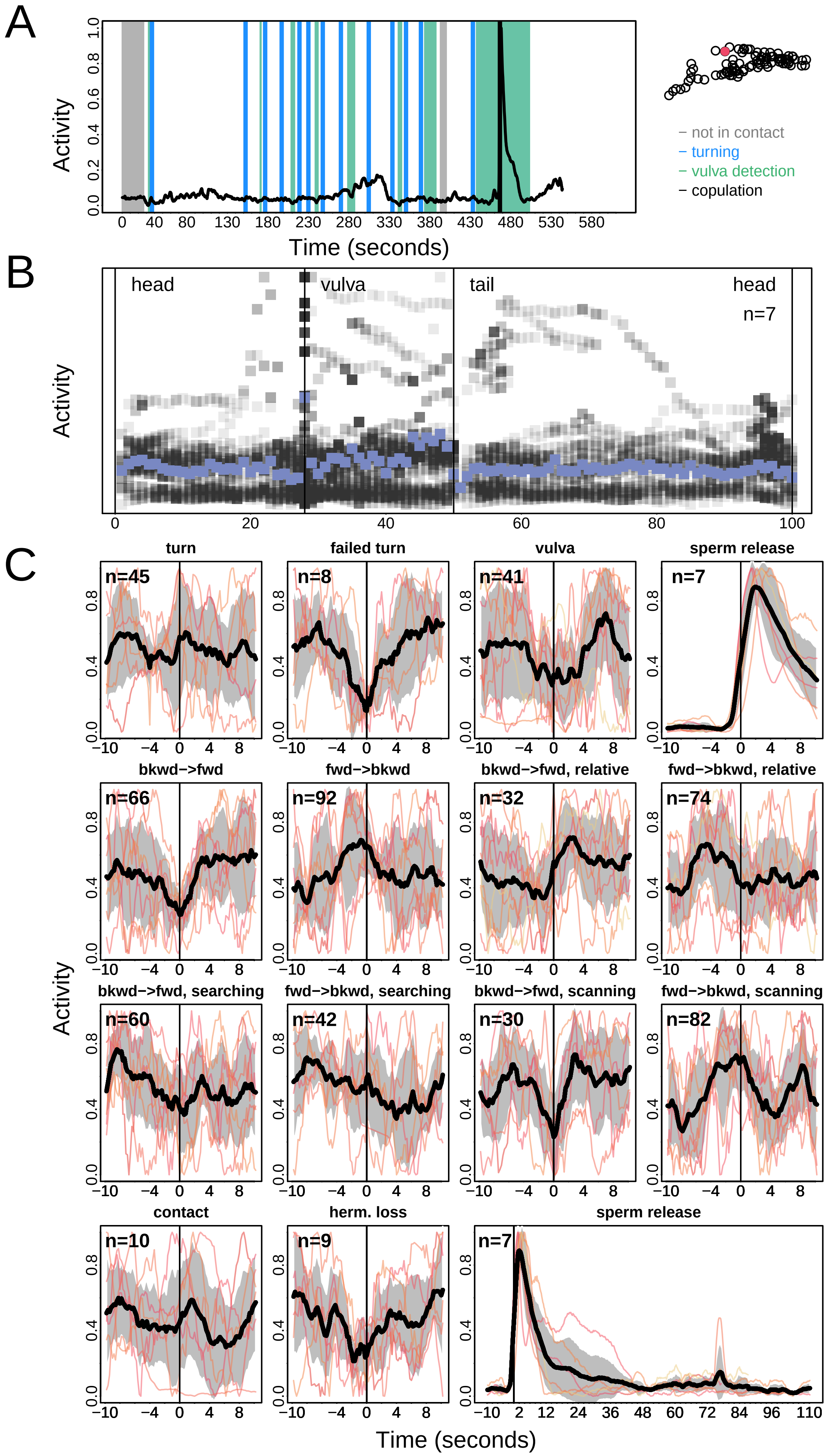
A
Activity
Time (seconds)
not in contact
turning
vulva detection
copulation
B
head
vulva
tail
head
n=7
Activity
C
turn
n=45
failed turn
n=8
vulva
n=41
sperm release
n=7
bkwd−>fwd
n=66
fwd−>bkwd
n=92
bkwd−>fwd, relative
n=32
fwd−>bkwd, relative
n=74
bkwd−>fwd, searching
n=60
fwd−>bkwd, searching
n=42
bkwd−>fwd, scanning
n=30
fwd−>bkwd, scanning
n=82
contact
n=10
herm. loss
n=9
sperm release
n=7
Activity
Time (seconds)

### PCA

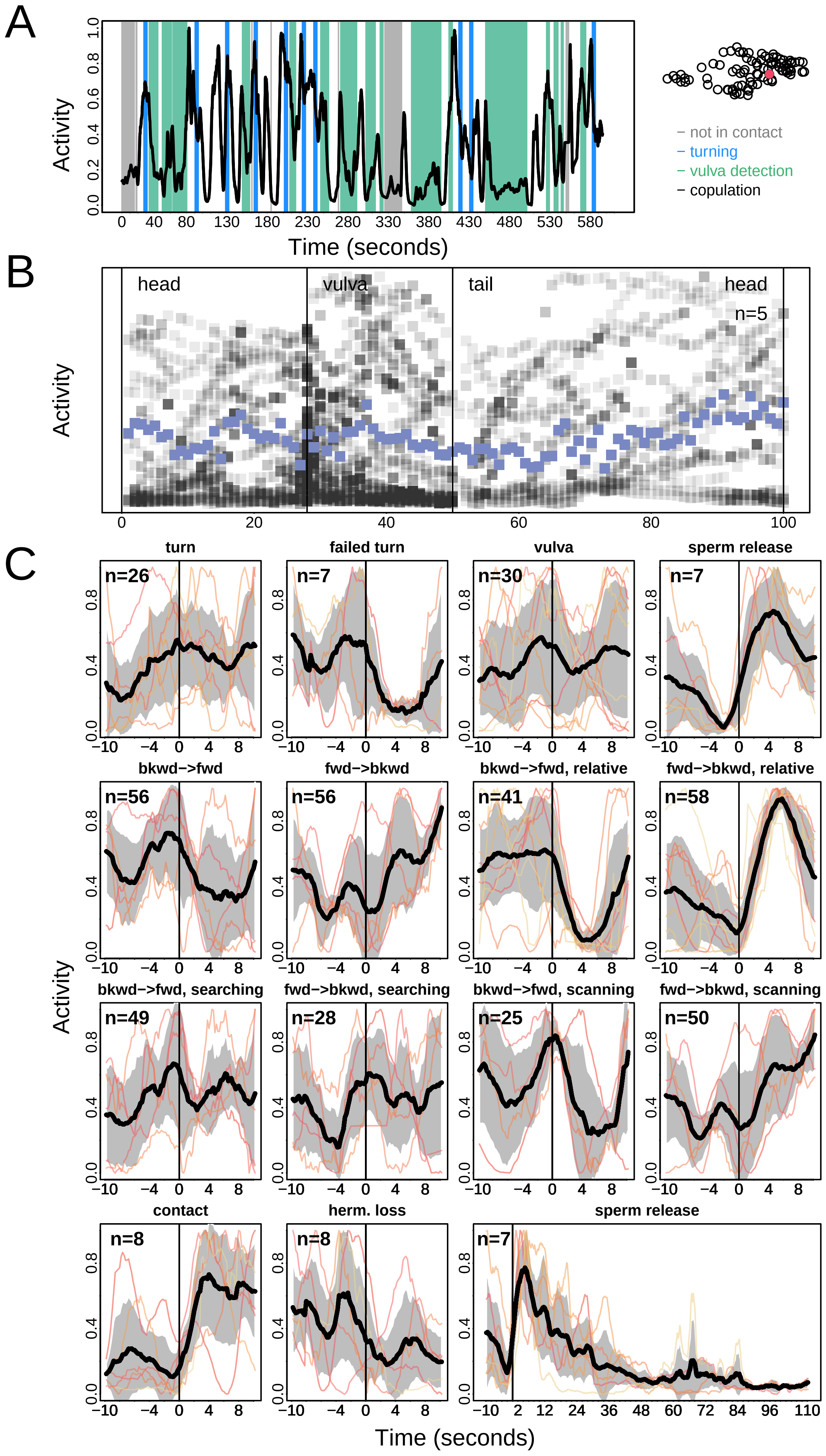
A
Activity
Time (seconds)
– not in contact
– turning
– vulva detection
– copulation
B
head
vulva
tail
head
n=5
Activity
C
turn
failed turn
vulva
sperm release
bkwd−>fwd
fwd−>bkwd
bkwd−>fwd, relative
fwd−>bkwd, relative
bkwd−>fwd, searching
fwd−>bkwd, searching
bkwd−>fwd, scanning
fwd−>bkwd, scanning
contact
herm. loss
sperm release
n=26
n=7
n=30
n=7
n=56
n=56
n=41
n=58
n=49
n=28
n=25
n=50
n=8
n=8
n=7
Activity
Time (seconds)

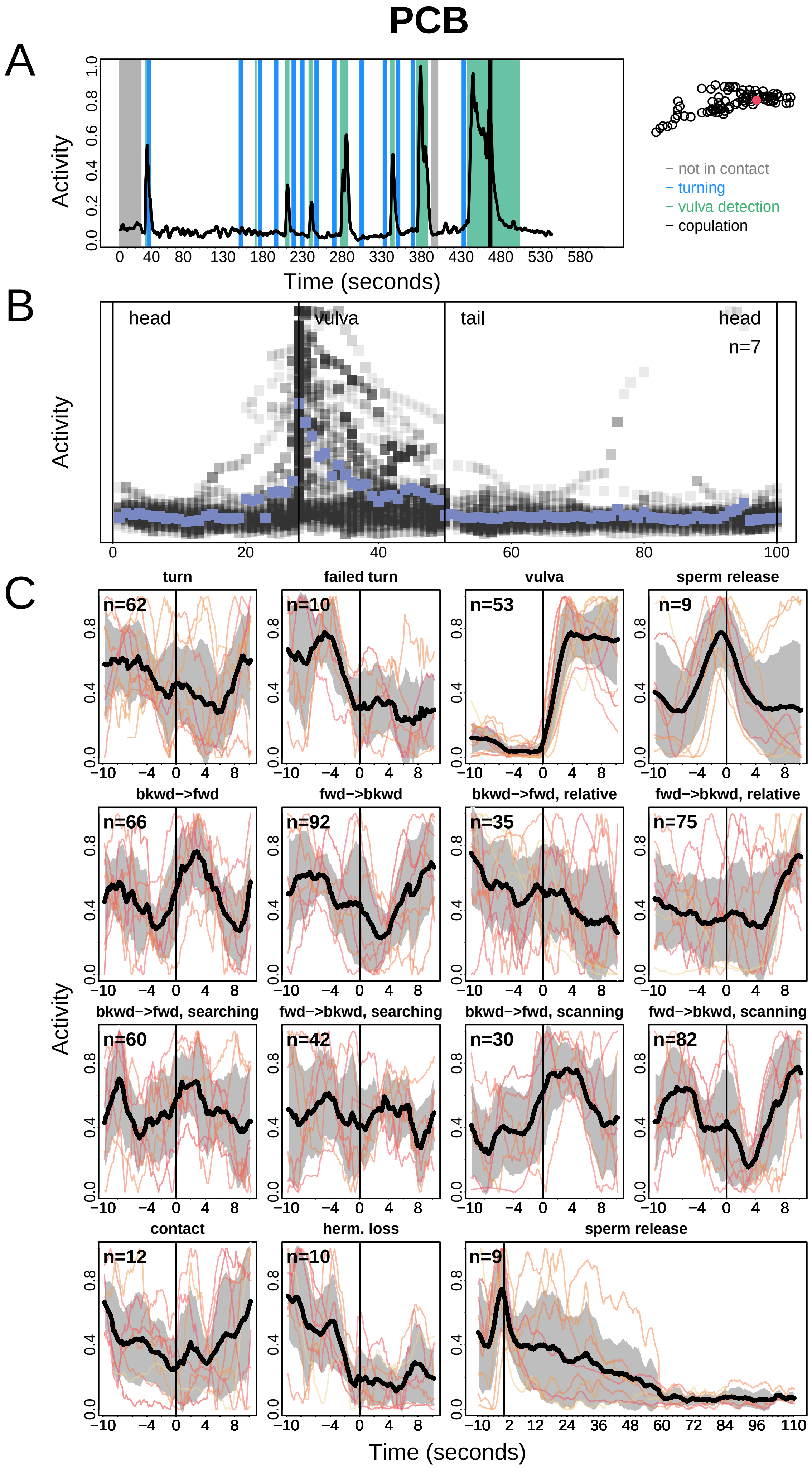

PCB
A
Activity
Time (seconds)
– not in contact
– turning
– vulva detection
– copulation
B
head
vulva
tail
head
n=7
Activity
C
turn
failed turn
vulva
sperm release
bkwd−>fwd
fwd−>bkwd
bkwd−>fwd, relative
fwd−>bkwd, relative
bkwd−>fwd, searching
fwd−>bkwd, searching
bkwd−>fwd, scanning
fwd−>bkwd, scanning
contact
herm. loss
sperm release
n=62
n=10
n=53
n=9
n=66
n=92
n=35
n=75
n=60
n=42
n=30
n=82
n=12
n=10
n=9
Activity
Time (seconds)

### PCC

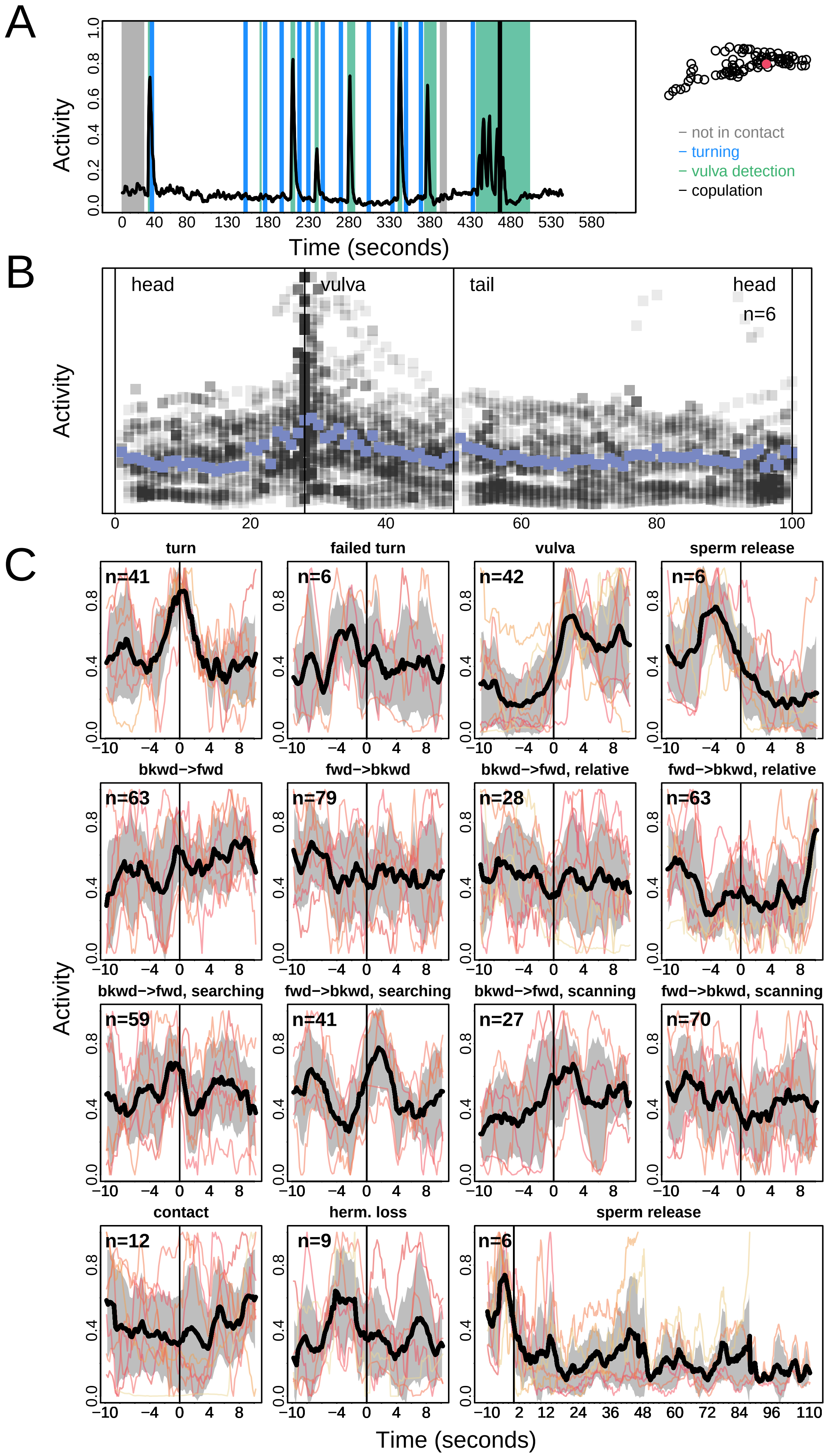

A
Activity
Time (seconds)
– not in contact
– turning
– vulva detection
– copulation
B
head
vulva
tail
head
n=6
Activity
C
turn
failed turn
vulva
sperm release
bkwd->fwd
fwd->bkwd
bkwd->fwd, relative
fwd->bkwd, relative
bkwd->fwd, searching
fwd->bkwd, searching
bkwd->fwd, scanning
fwd->bkwd, scanning
contact
herm. loss
sperm release
n=41
n=6
n=42
n=6
n=63
n=79
n=28
n=63
n=59
n=41
n=27
n=70
n=12
n=9
n=6
Activity
Time (seconds)

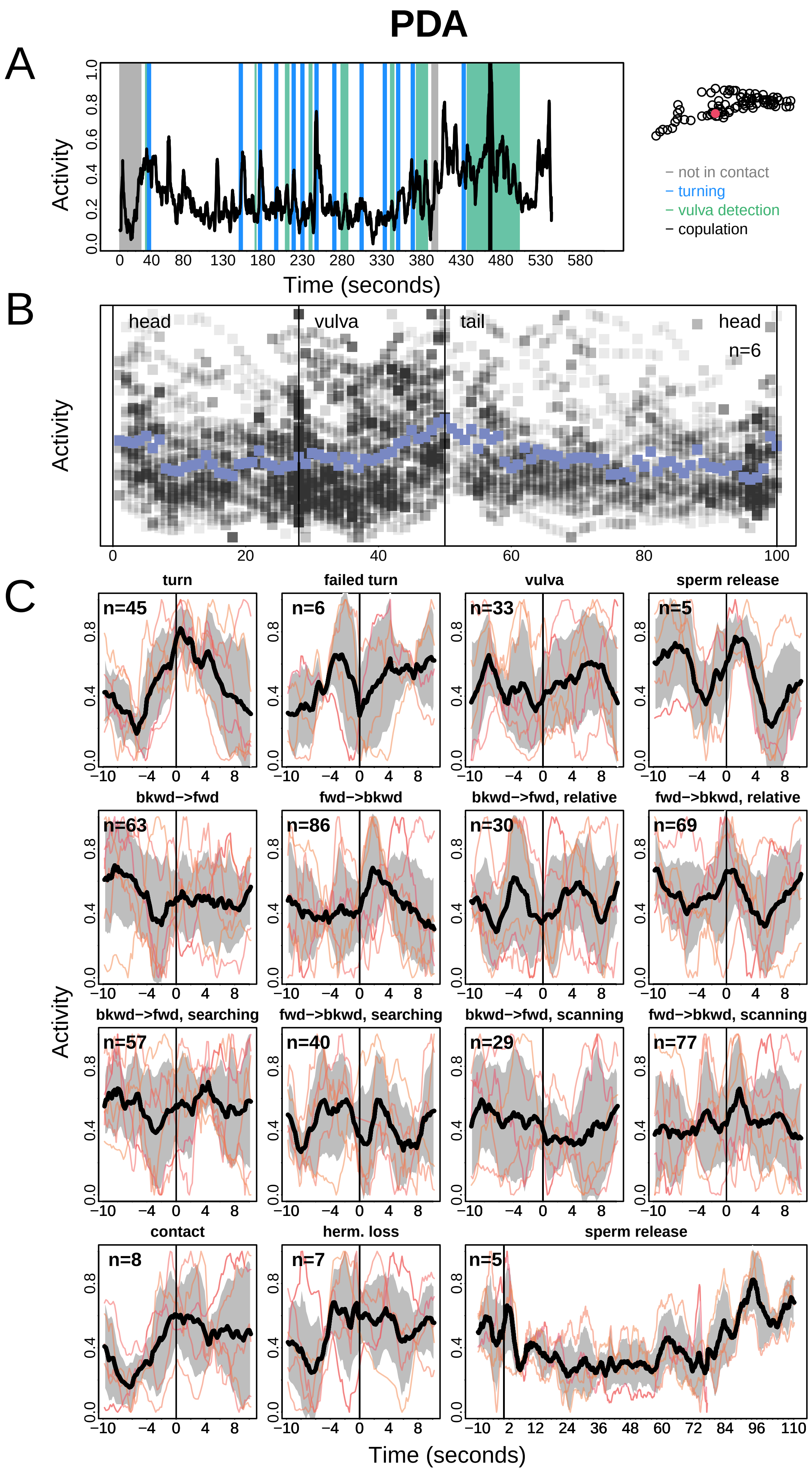

PDA
A
Activity
Time (seconds)
− not in contact
− turning
− vulva detection
− copulation
B
head
vulva
tail
head
n=6
Activity
C
turn
failed turn
vulva
sperm release
n=45
n=6
n=33
n=5
bkwd−>fwd
fwd−>bkwd
bkwd−>fwd, relative
fwd−>bkwd, relative
n=63
n=86
n=30
n=69
bkwd−>fwd, searching
fwd−>bkwd, searching
bkwd−>fwd, scanning
fwd−>bkwd, scanning
n=57
n=40
n=29
n=77
contact
herm. loss
sperm release
n=8
n=7
n=5
Activity
Time (seconds)

### PDB

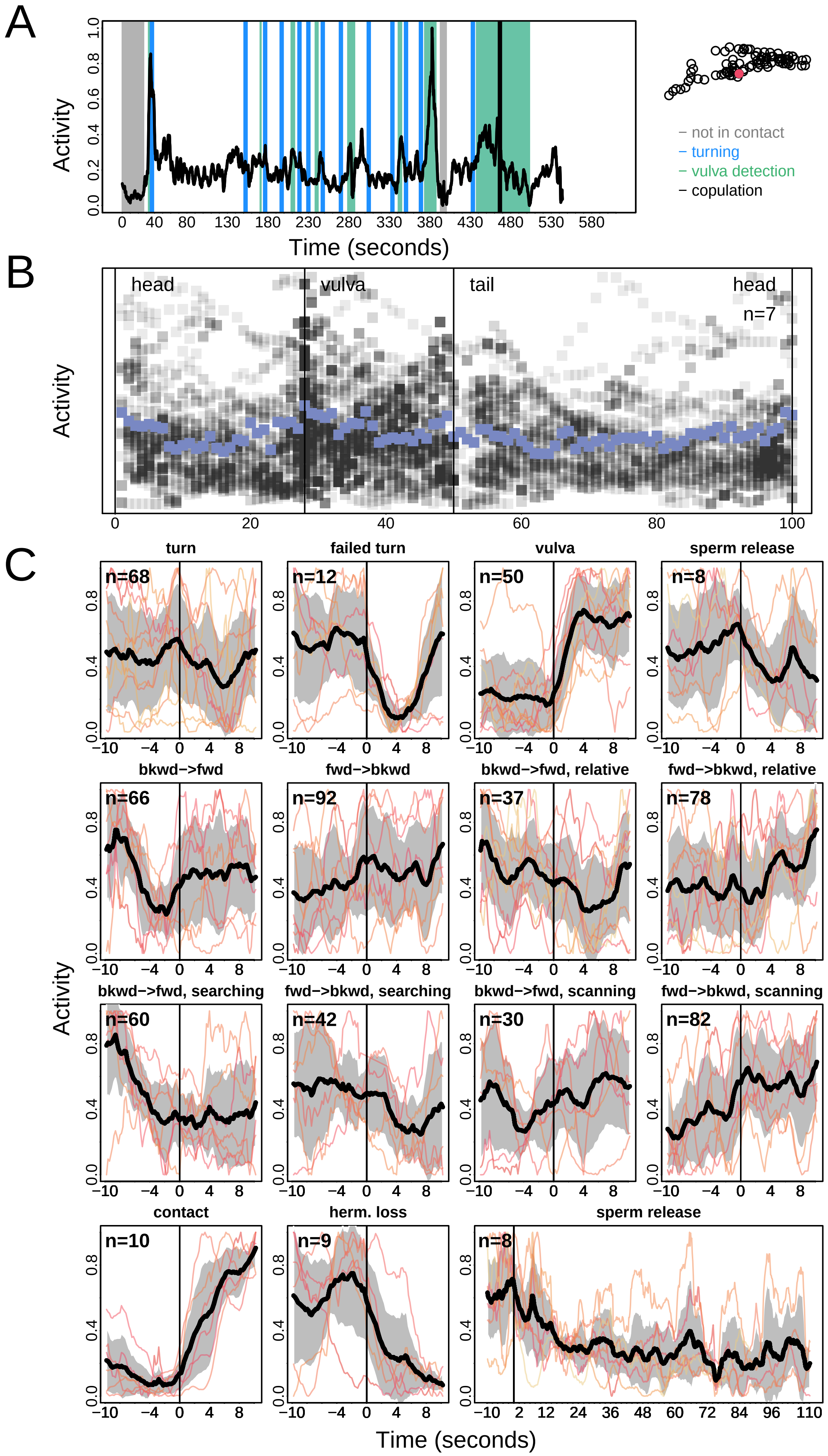

A
B
C
Activity
Time (seconds)
not in contact
turning
vulva detection
copulation
head
vulva
tail
head
n=7
turn
failed turn
vulva
sperm release
bkwd->fwd
fwd->bkwd
bkwd->fwd, relative
fwd->bkwd, relative
bkwd->fwd, searching
fwd->bkwd, searching
bkwd->fwd, scanning
fwd->bkwd, scanning
contact
herm. loss
sperm release
n=68
n=12
n=50
n=8
n=66
n=92
n=37
n=78
n=60
n=42
n=30
n=82
n=10
n=9
n=8

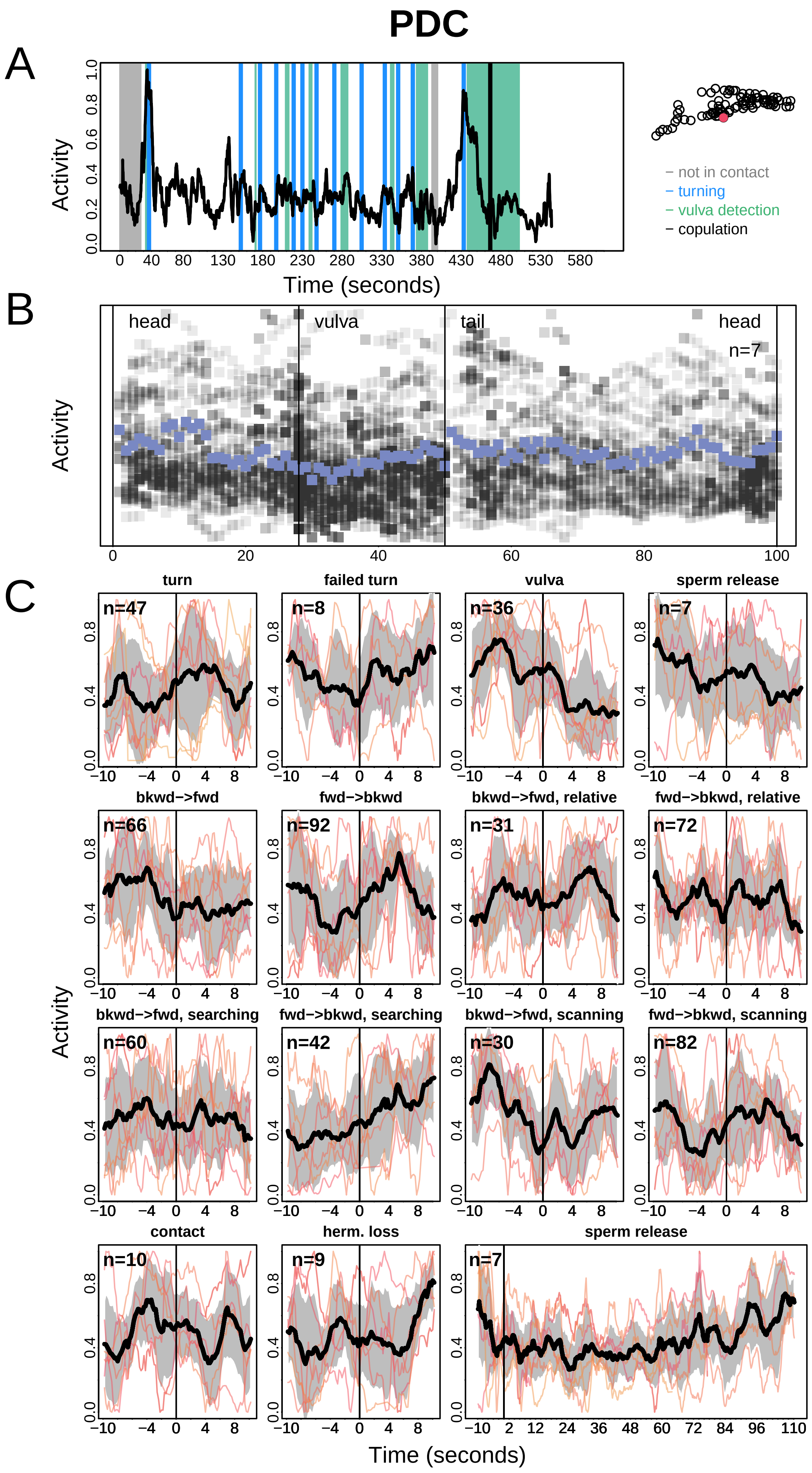
PDC
A
Activity
Time (seconds)
– not in contact
– turning
– vulva detection
– copulation
B
head
vulva
tail
head
n=7
Activity
C
turn
failed turn
vulva
sperm release
bkwd−>fwd
fwd−>bkwd
bkwd−>fwd, relative
fwd−>bkwd, relative
bkwd−>fwd, searching
fwd−>bkwd, searching
bkwd−>fwd, scanning
fwd−>bkwd, scanning
contact
herm. loss
sperm release
n=47
n=8
n=36
n=7
n=66
n=92
n=31
n=72
n=60
n=42
n=30
n=82
n=10
n=9
n=7
Activity
Time (seconds)

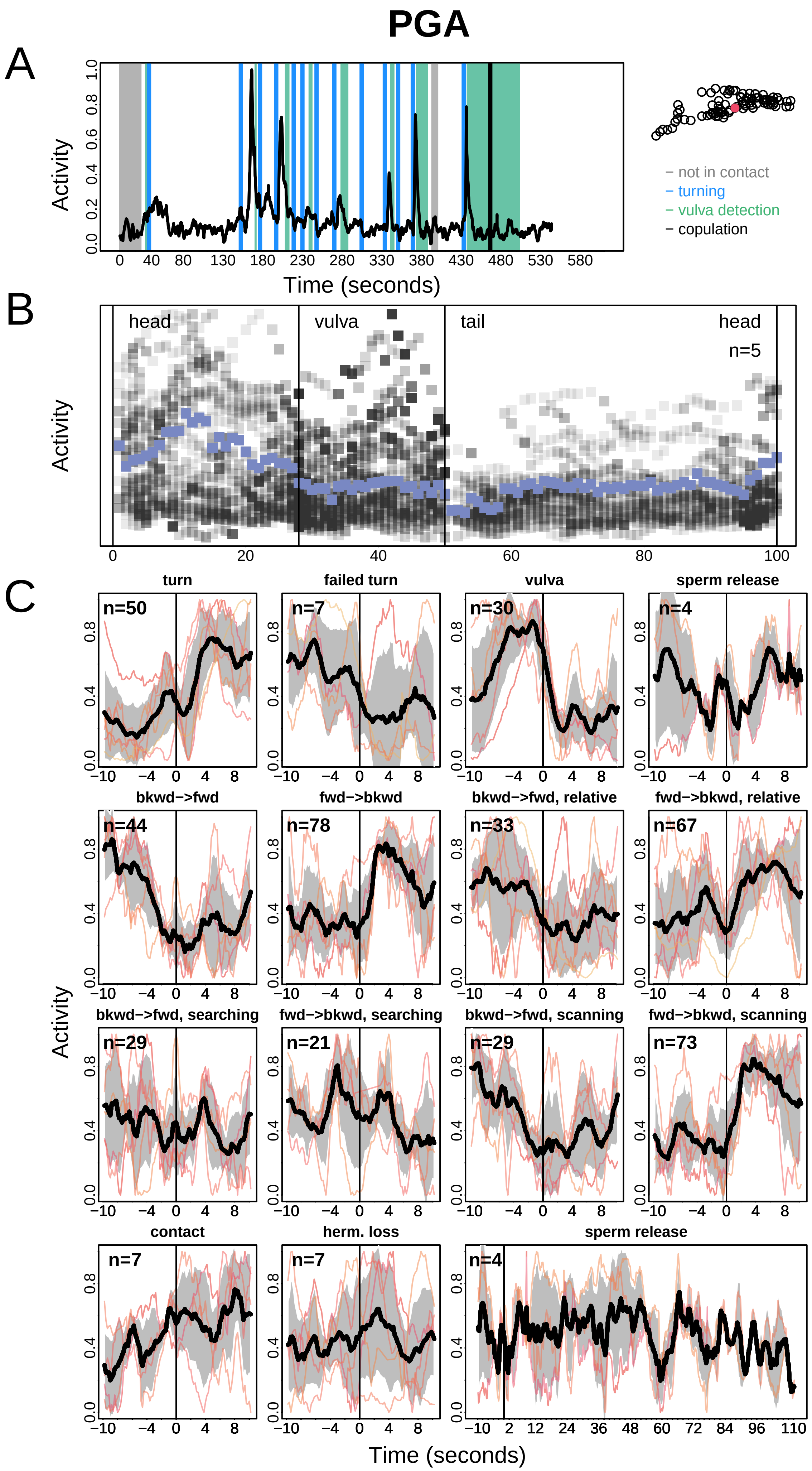
PGA
A
Activity
Time (seconds)
− not in contact
− turning
− vulva detection
− copulation
B
head
vulva
tail
head
n=5
Activity
C
turn
failed turn
vulva
sperm release
bkwd−>fwd
fwd−>bkwd
bkwd−>fwd, relative
fwd−>bkwd, relative
bkwd−>fwd, searching
fwd−>bkwd, searching
bkwd−>fwd, scanning
fwd−>bkwd, scanning
contact
herm. loss
sperm release
n=50
n=7
n=30
n=4
n=44
n=78
n=33
n=67
n=29
n=21
n=29
n=73
n=7
n=7
n=4
Activity
Time (seconds)

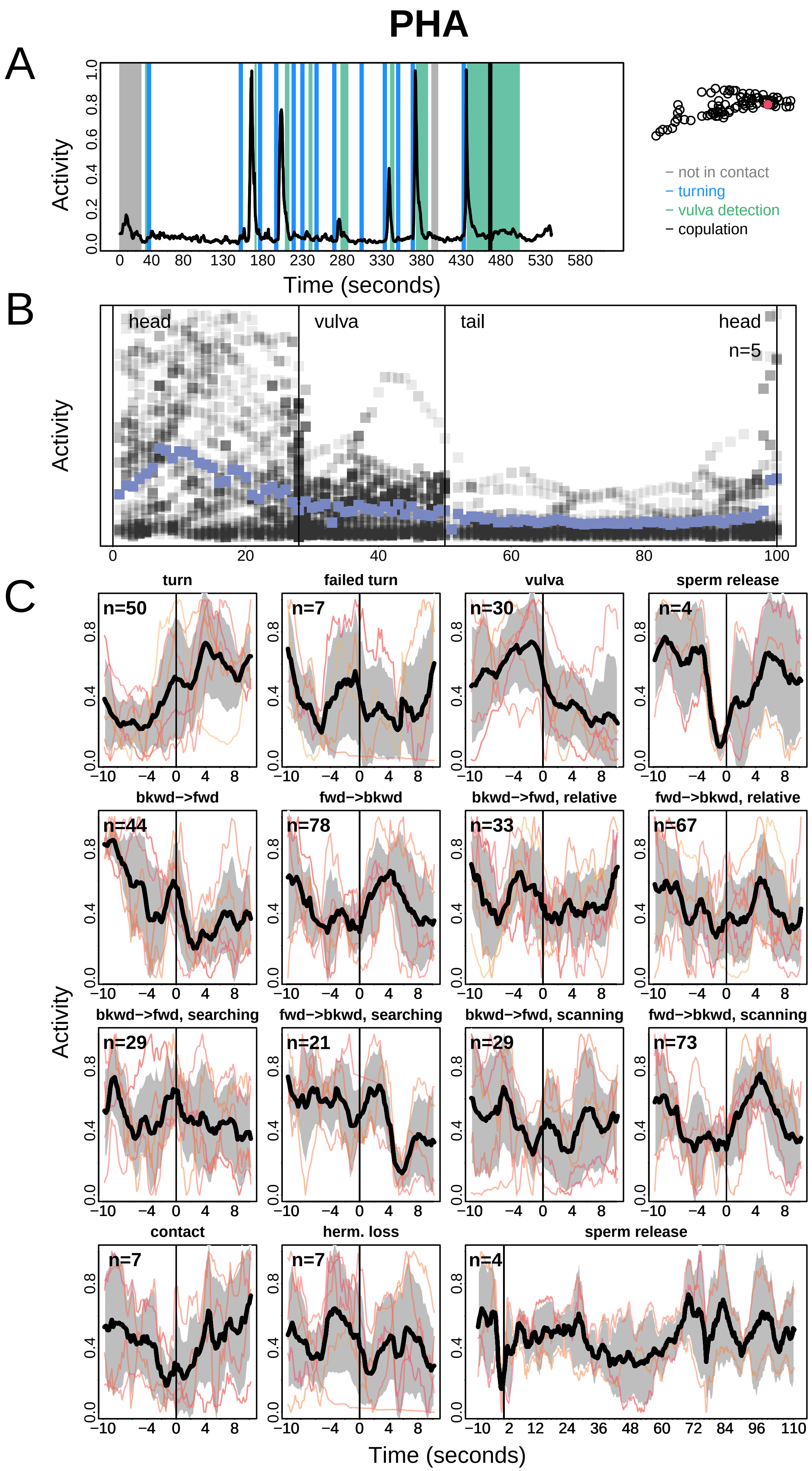

PHA
A
Activity
Time (seconds)
– not in contact
– turning
– vulva detection
– copulation
B
head
vulva
tail
head
n=5
Activity
C
turn
failed turn
vulva
sperm release
bkwd−>fwd
fwd−>bkwd
bkwd−>fwd, relative
fwd−>bkwd, relative
bkwd−>fwd, searching
fwd−>bkwd, searching
bkwd−>fwd, scanning
fwd−>bkwd, scanning
contact
herm. loss
sperm release
Activity
Time (seconds)

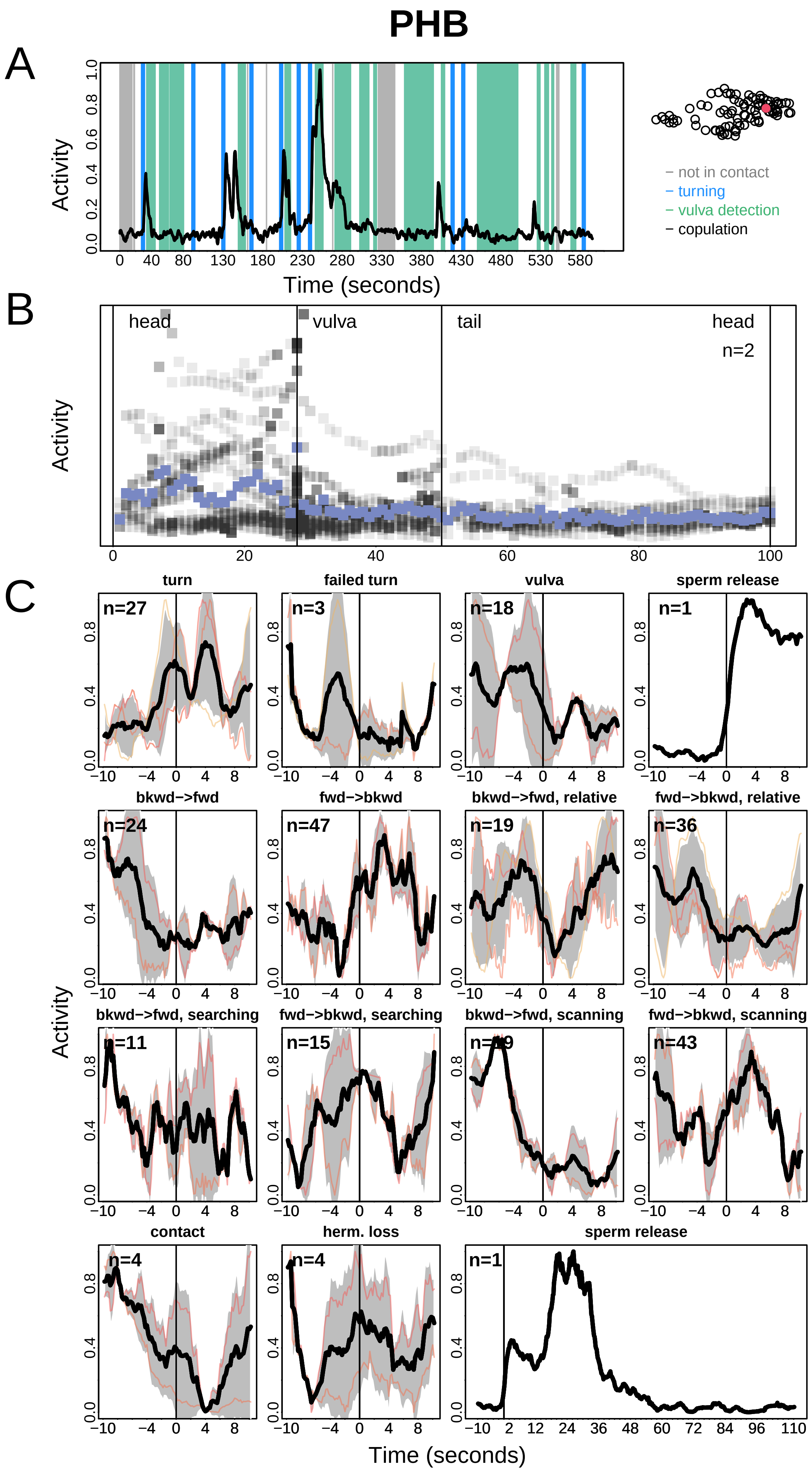
PHB
A
Activity
Time (seconds)
− not in contact
− turning
− vulva detection
− copulation
B
head
vulva
tail
head
n=2
Activity
C
turn
failed turn
vulva
sperm release
bkwd−>fwd
fwd−>bkwd
bkwd−>fwd, relative
fwd−>bkwd, relative
bkwd−>fwd, searching
fwd−>bkwd, searching
bkwd−>fwd, scanning
fwd−>bkwd, scanning
contact
herm. loss
sperm release
Activity
Time (seconds)

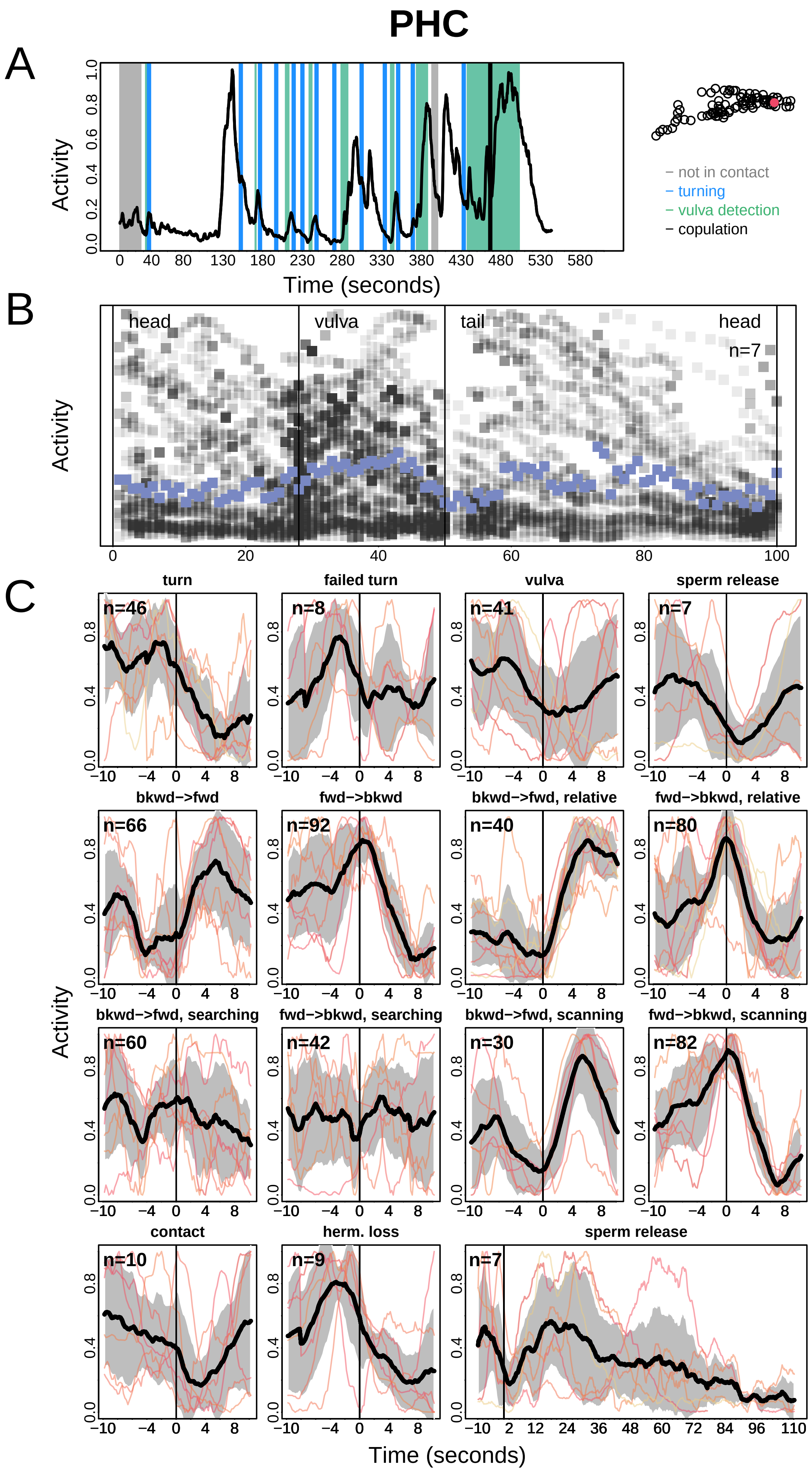

PHC
A
Activity
Time (seconds)
not in contact
turning
vulva detection
copulation
B
head
vulva
tail
head
n=7
Activity
C
turn
failed turn
vulva
sperm release
bkwd−>fwd
fwd−>bkwd
bkwd−>fwd, relative
fwd−>bkwd, relative
bkwd−>fwd, searching
fwd−>bkwd, searching
bkwd−>fwd, scanning
fwd−>bkwd, scanning
contact
herm. loss
sperm release
n=46
n=8
n=41
n=7
n=66
n=92
n=40
n=80
n=60
n=42
n=30
n=82
n=10
n=9
n=7
Activity
Time (seconds)

### PHD

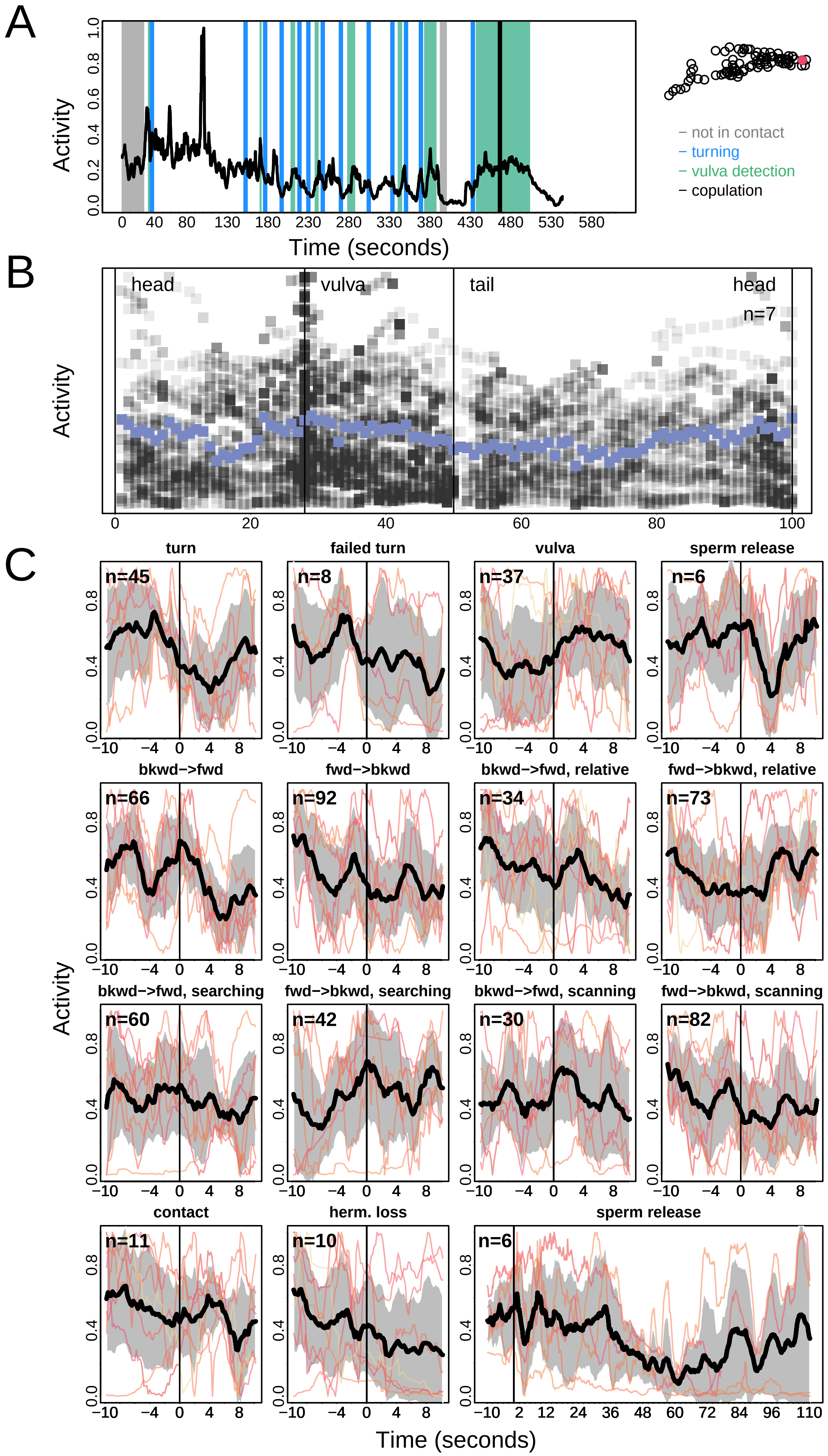
A
Activity
Time (seconds)
not in contact
turning
vulva detection
copulation
B
head
vulva
tail
head
n=7
Activity
C
turn
failed turn
vulva
sperm release
bkwd−>fwd
fwd−>bkwd
bkwd−>fwd, relative
fwd−>bkwd, relative
bkwd−>fwd, searching
fwd−>bkwd, searching
bkwd−>fwd, scanning
fwd−>bkwd, scanning
contact
herm. loss
sperm release
Activity
Time (seconds)

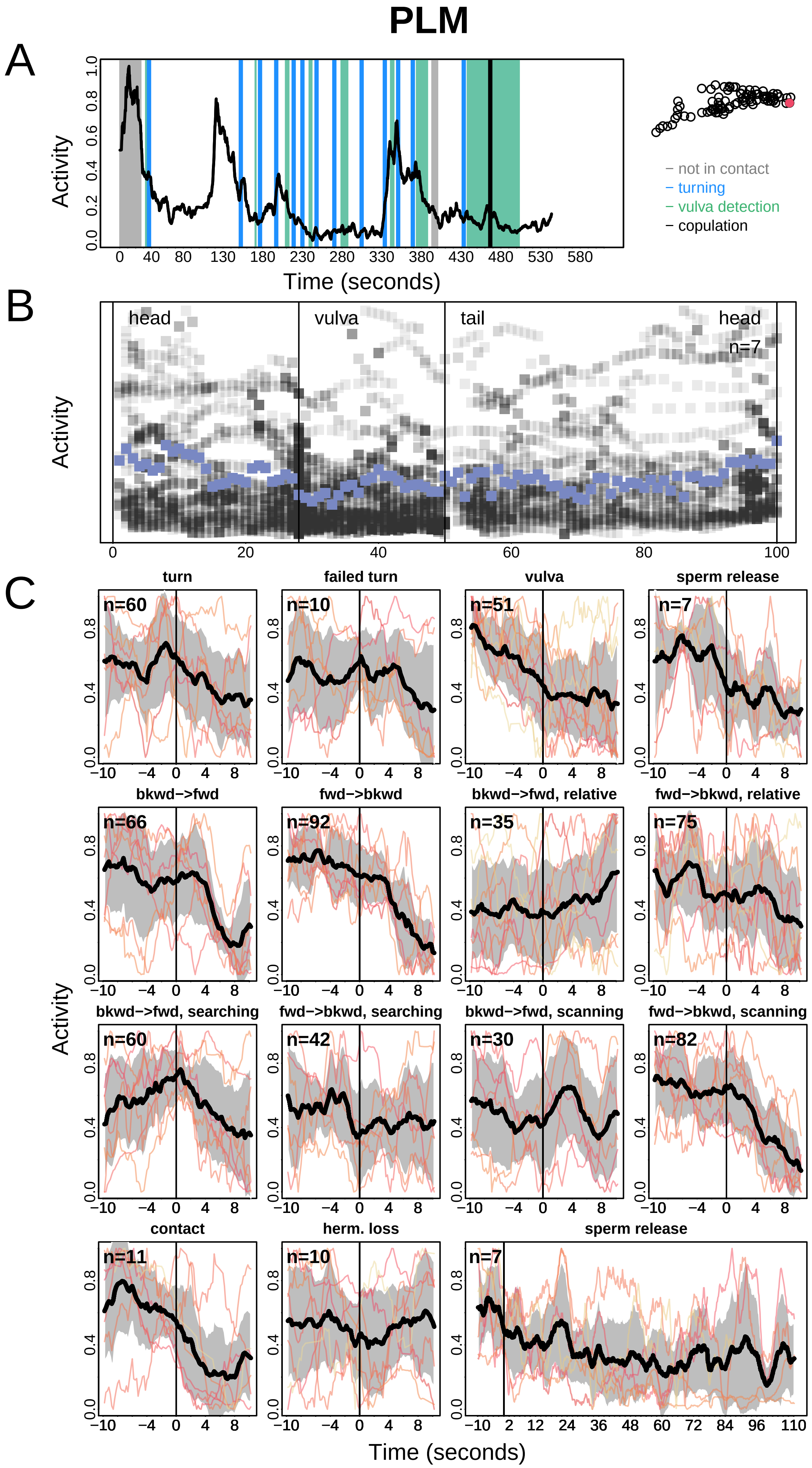
PLM
A
Activity
Time (seconds)
not in contact
turning
vulva detection
copulation
B
head
vulva
tail
head
n=7
Activity
C
turn
failed turn
vulva
sperm release
bkwd−>fwd
fwd−>bkwd
bkwd−>fwd, relative
fwd−>bkwd, relative
bkwd−>fwd, searching
fwd−>bkwd, searching
bkwd−>fwd, scanning
fwd−>bkwd, scanning
contact
herm. loss
sperm release
n=60
n=10
n=51
n=7
n=66
n=92
n=35
n=75
n=60
n=42
n=30
n=82
n=11
n=10
n=7
Activity
Time (seconds)

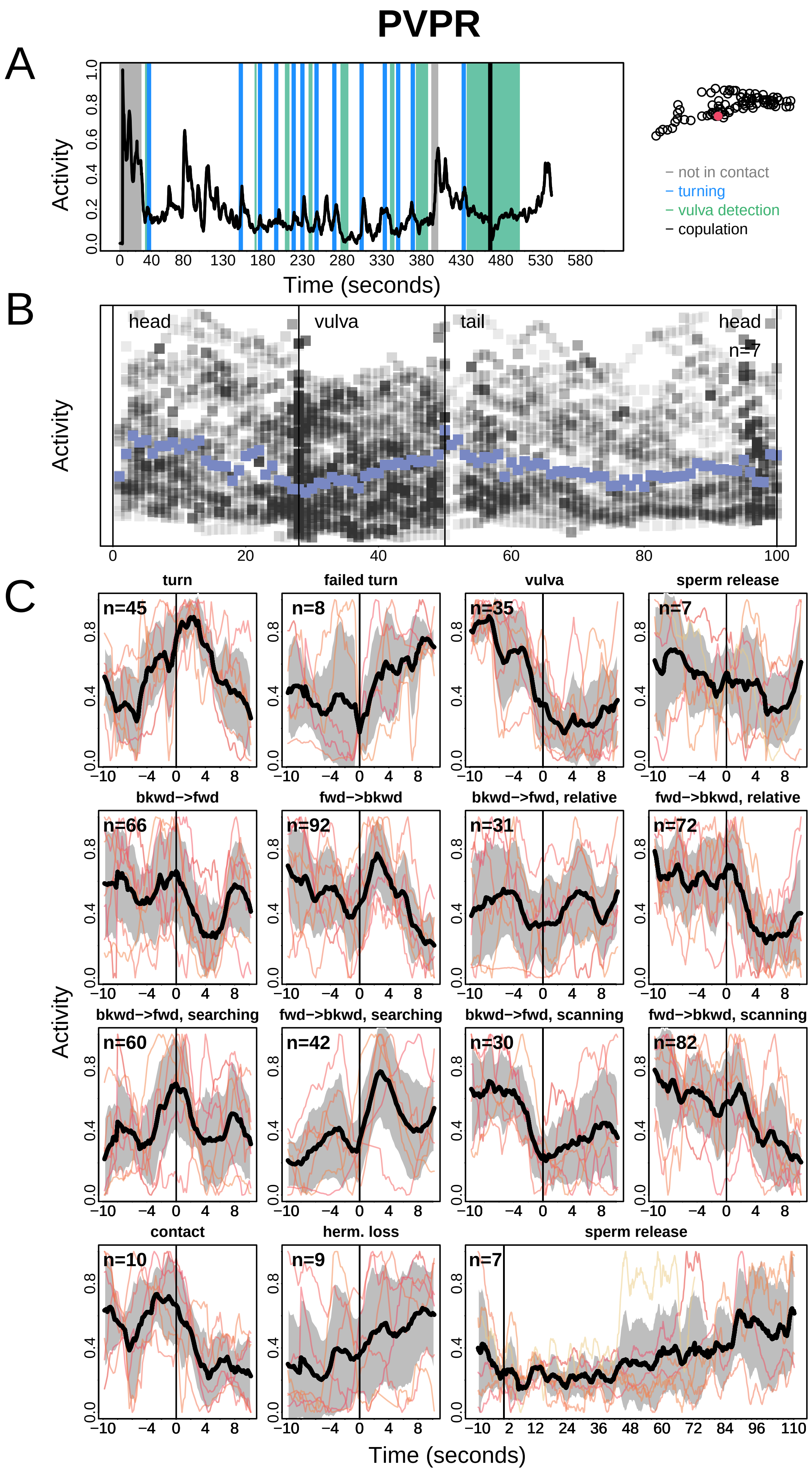
PVPR
A
Activity
Time (seconds)
– not in contact
– turning
– vulva detection
– copulation
B
head
vulva
tail
head
n=7
Activity
C
turn
failed turn
vulva
sperm release
bkwd−>fwd
fwd−>bkwd
bkwd−>fwd, relative
fwd−>bkwd, relative
bkwd−>fwd, searching
fwd−>bkwd, searching
bkwd−>fwd, scanning
fwd−>bkwd, scanning
contact
herm. loss
sperm release
Activity
Time (seconds)

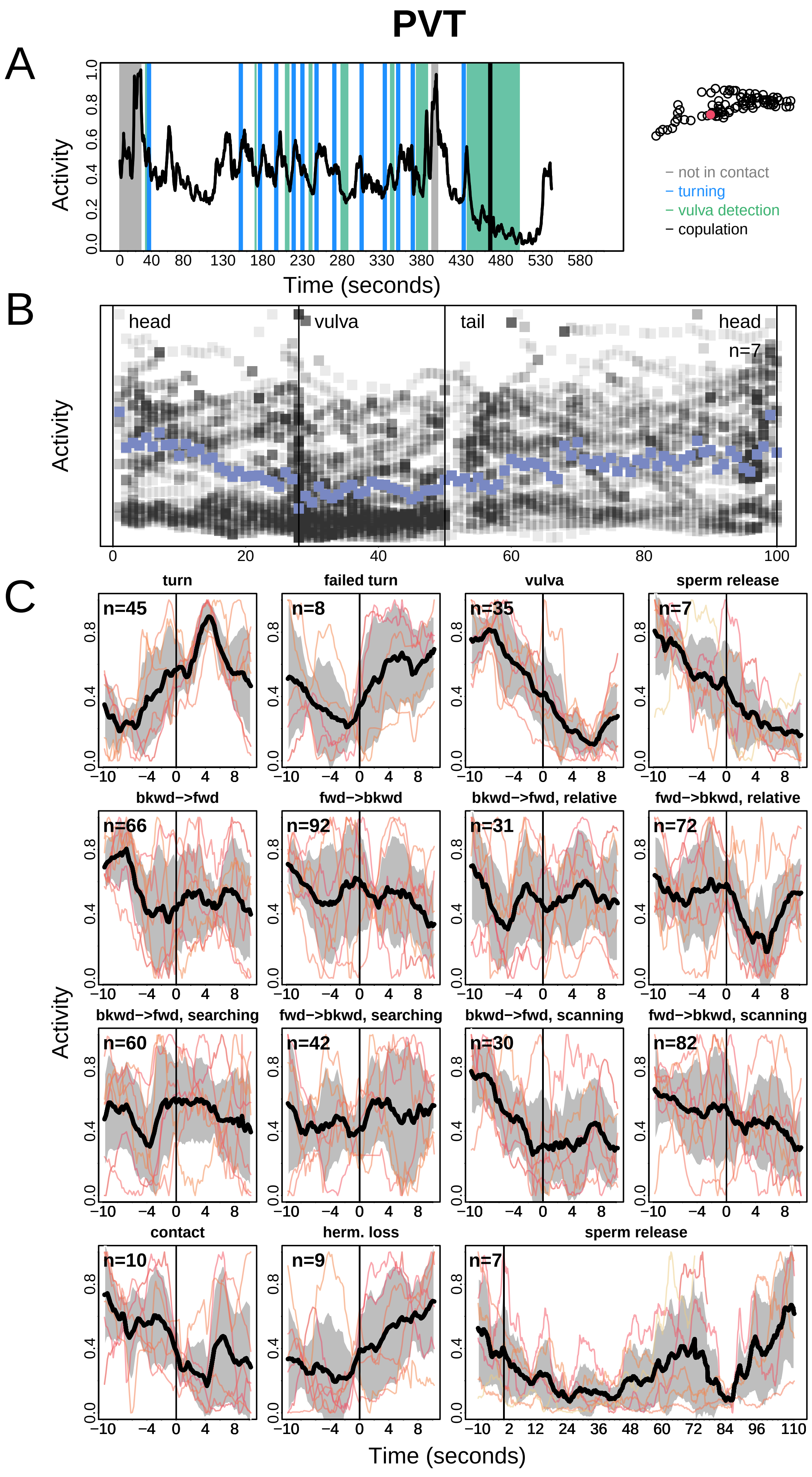
PVT
A
B
C
Activity
Time (seconds)
not in contact
turning
vulva detection
copulation
head
vulva
tail
head
n=7
turn
failed turn
vulva
sperm release
bkwd−>fwd
fwd−>bkwd
bkwd−>fwd, relative
fwd−>bkwd, relative
bkwd−>fwd, searching
fwd−>bkwd, searching
bkwd−>fwd, scanning
fwd−>bkwd, scanning
contact
herm. loss
sperm release
n=45
n=8
n=35
n=7
n=66
n=92
n=31
n=72
n=60
n=42
n=30
n=82
n=10
n=9
n=7

### PVV

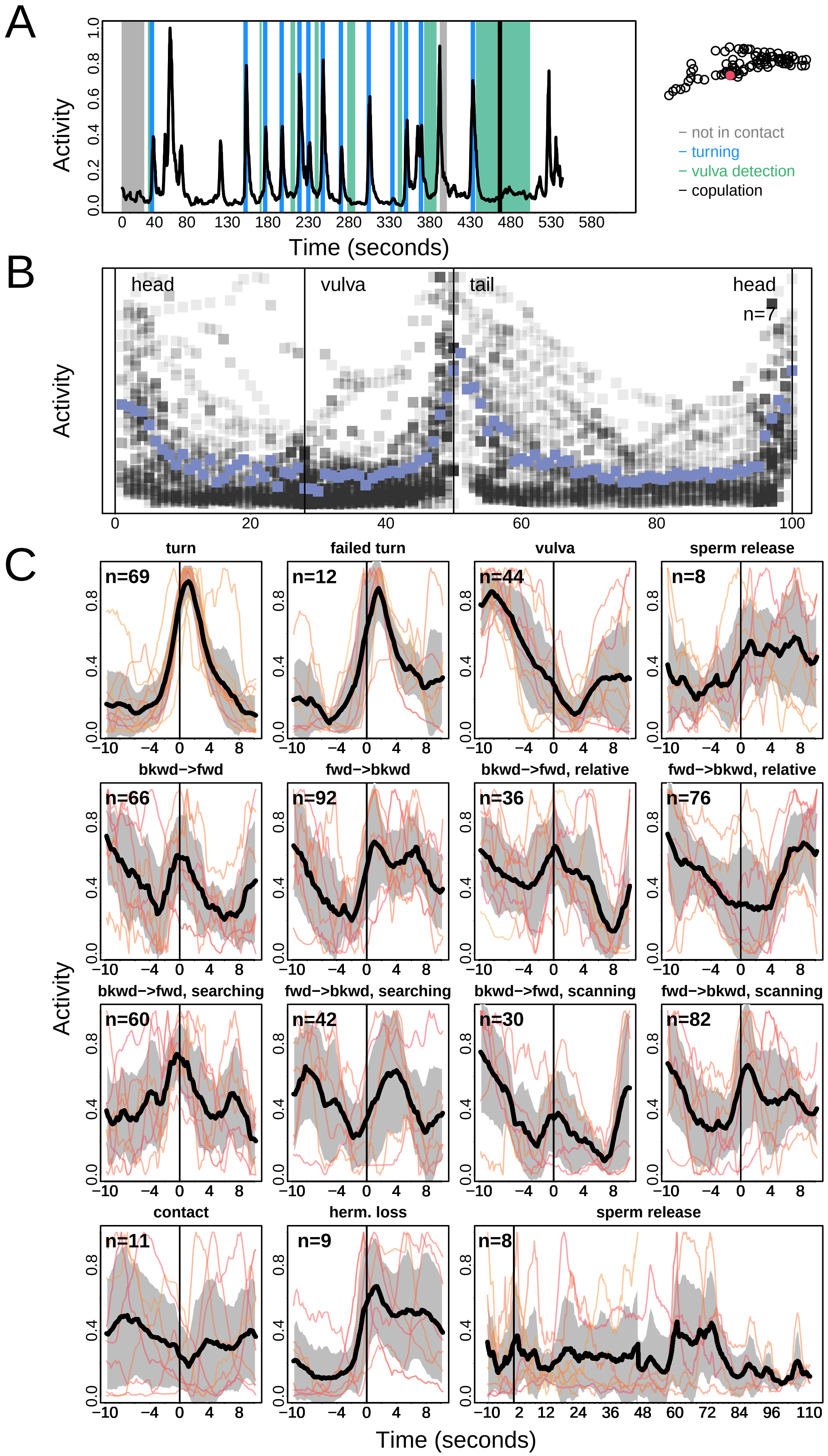
A
Activity
Time (seconds)
not in contact
turning
vulva detection
copulation
B
head
vulva
tail
head
n=7
Activity
C
turn
failed turn
vulva
sperm release
bkwd−>fwd
fwd−>bkwd
bkwd−>fwd, relative
fwd−>bkwd, relative
bkwd−>fwd, searching
fwd−>bkwd, searching
bkwd−>fwd, scanning
fwd−>bkwd, scanning
contact
herm. loss
sperm release
n=69
n=12
n=44
n=8
n=66
n=92
n=36
n=76
n=60
n=42
n=30
n=82
n=11
n=9
n=8
Activity
Time (seconds)

### PVX

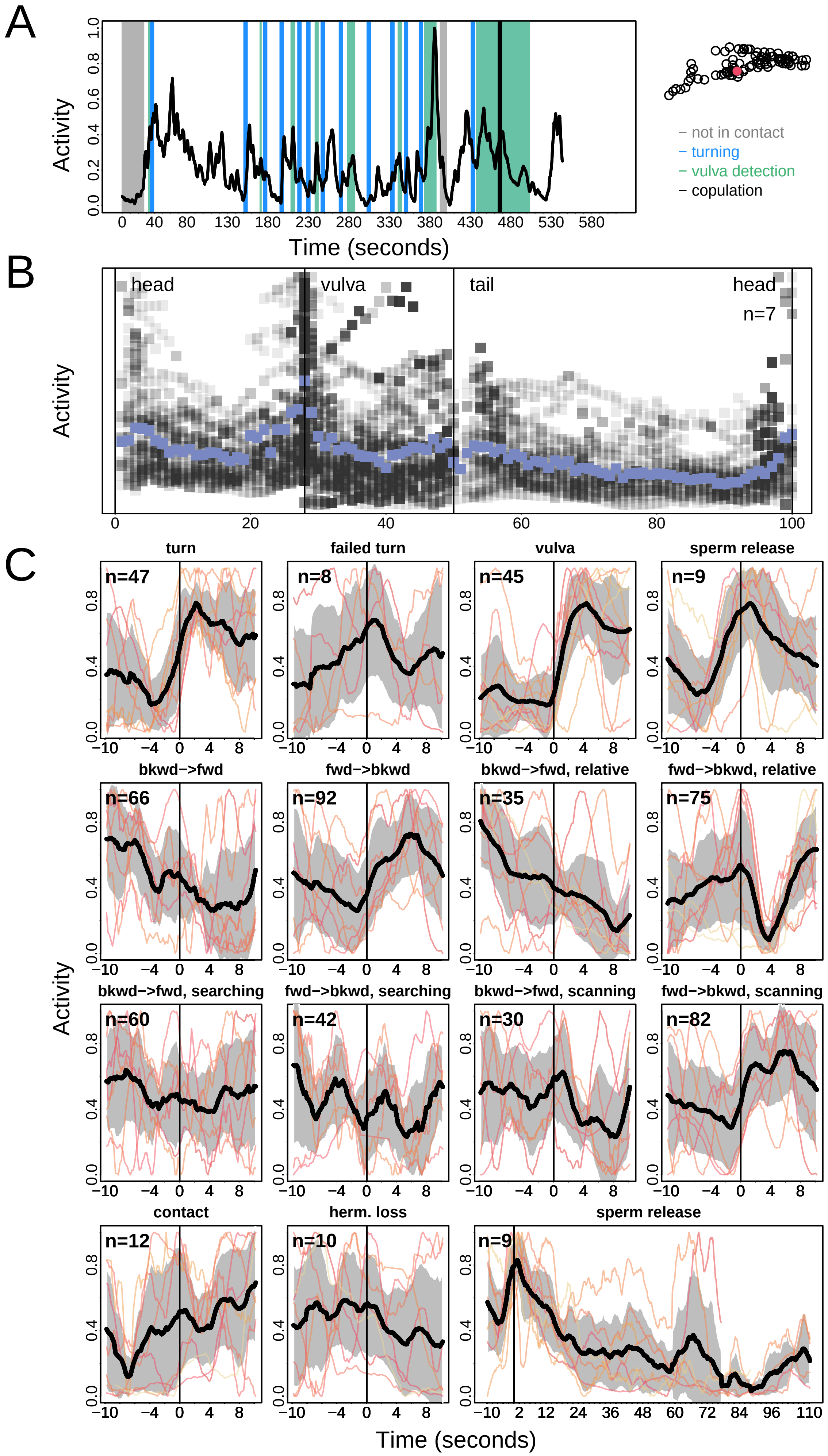

A
Activity
Time (seconds)
not in contact
turning
vulva detection
copulation
B
head
vulva
tail
head
n=7
Activity
C
turn
failed turn
vulva
sperm release
bkwd−>fwd
fwd−>bkwd
bkwd−>fwd, relative
fwd−>bkwd, relative
bkwd−>fwd, searching
fwd−>bkwd, searching
bkwd−>fwd, scanning
fwd−>bkwd, scanning
contact
herm. loss
sperm release
n=47
n=8
n=45
n=9
n=66
n=92
n=35
n=75
n=60
n=42
n=30
n=82
n=12
n=10
n=9
Activity
Time (seconds)

### PVY

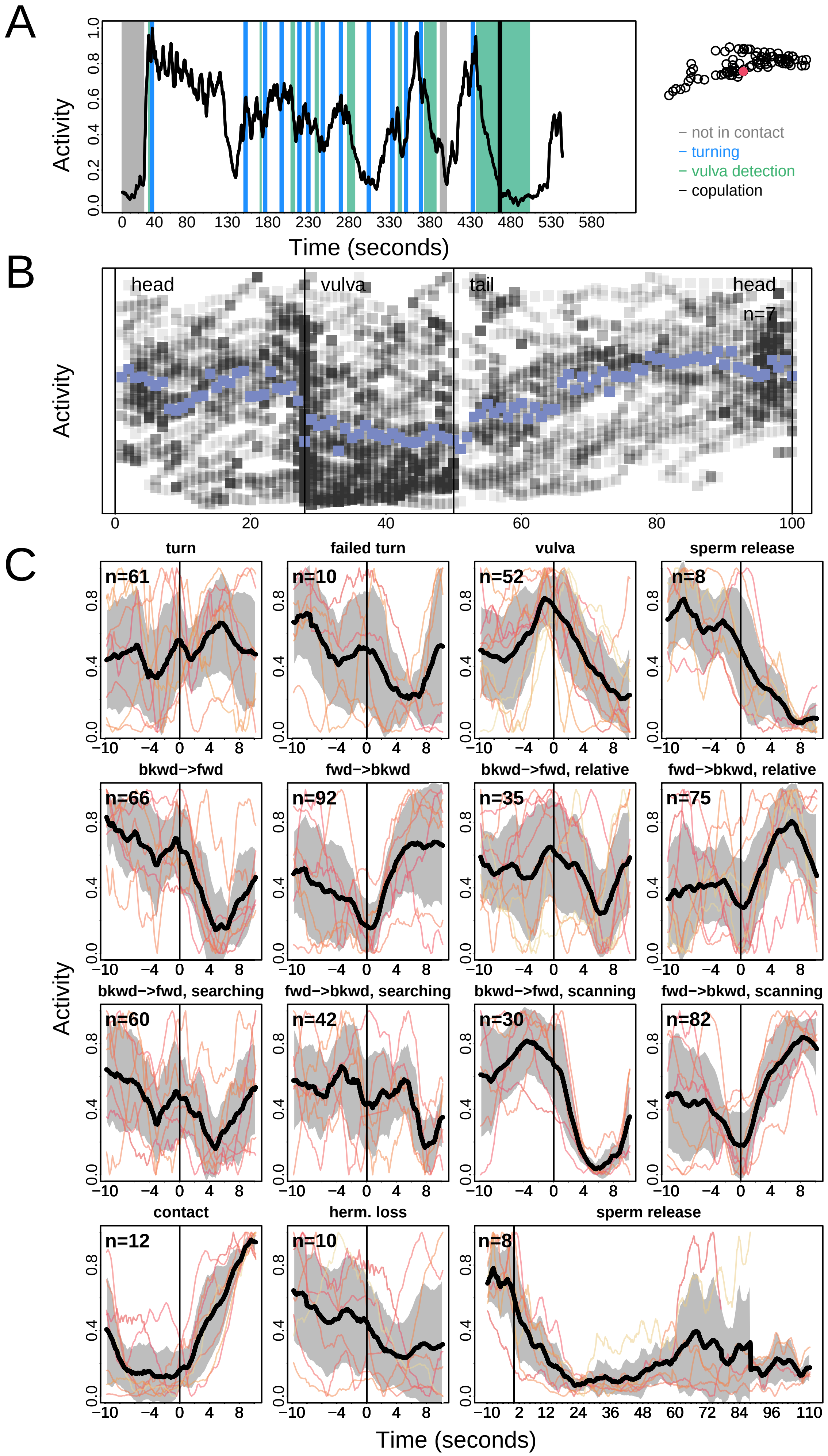
A
B
C
Activity
Time (seconds)
not in contact
turning
vulva detection
copulation
head
vulva
tail
head
n=7
turn
failed turn
vulva
sperm release
bkwd−>fwd
fwd−>bkwd
bkwd−>fwd, relative
fwd−>bkwd, relative
bkwd−>fwd, searching
fwd−>bkwd, searching
bkwd−>fwd, scanning
fwd−>bkwd, scanning
contact
herm. loss
sperm release
n=61
n=10
n=52
n=8
n=66
n=92
n=35
n=75
n=60
n=42
n=30
n=82
n=12
n=10
n=8

### PVZ

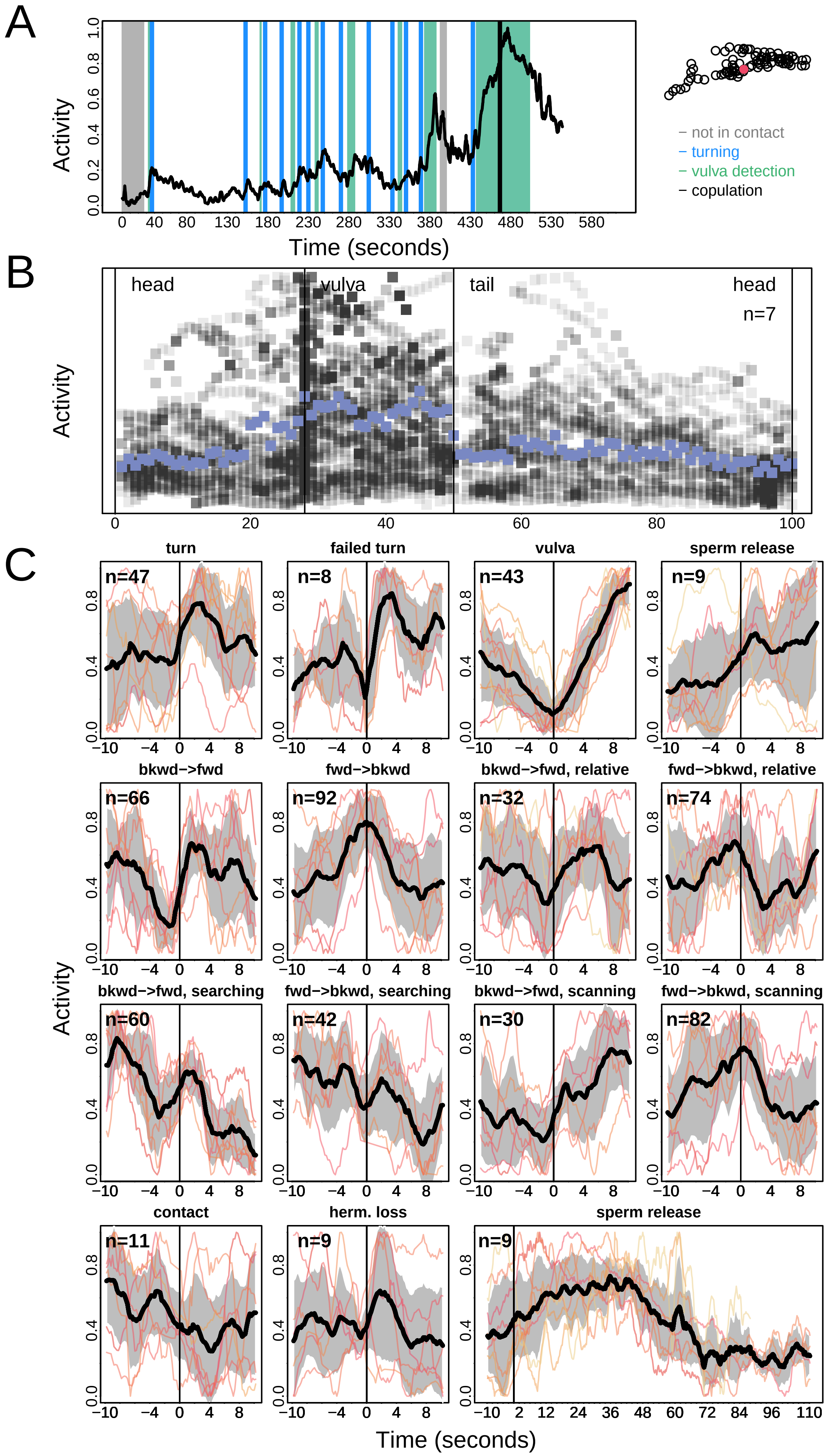
A
Activity
Time (seconds)
not in contact
turning
vulva detection
copulation
B
head
vulva
tail
head
n=7
Activity
C
turn
n=47
failed turn
n=8
vulva
n=43
sperm release
n=9
bkwd−>fwd
n=66
fwd−>bkwd
n=92
bkwd−>fwd, relative
n=32
fwd−>bkwd, relative
n=74
bkwd−>fwd, searching
n=60
fwd−>bkwd, searching
n=42
bkwd−>fwd, scanning
n=30
fwd−>bkwd, scanning
n=82
contact
n=11
herm. loss
n=9
sperm release
n=9
Activity
Time (seconds)

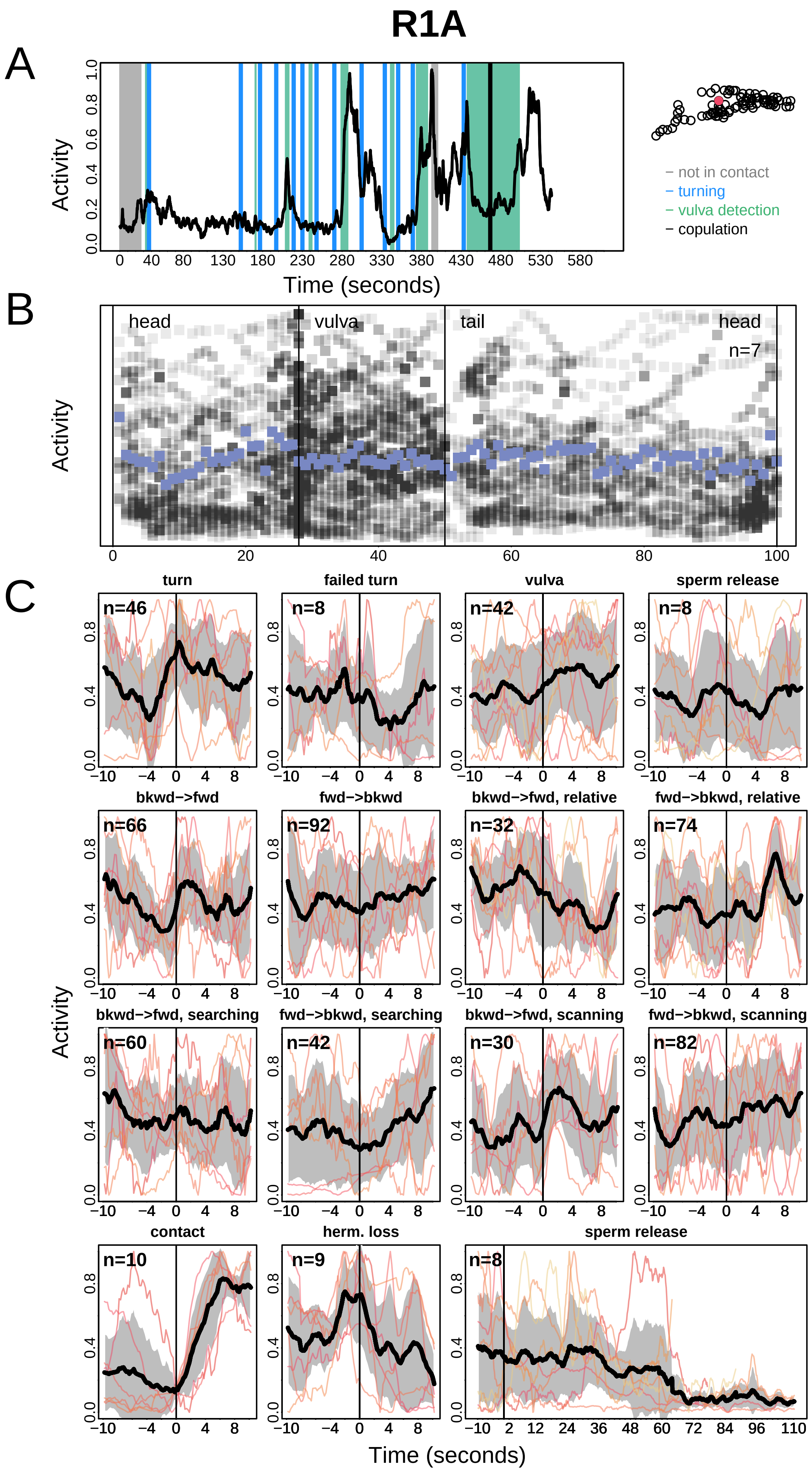
R1A
A
Activity
Time (seconds)
not in contact
turning
vulva detection
copulation
B
head
vulva
tail
head
n=7
Activity
C
turn
failed turn
vulva
sperm release
bkwd−>fwd
fwd−>bkwd
bkwd−>fwd, relative
fwd−>bkwd, relative
bkwd−>fwd, searching
fwd−>bkwd, searching
bkwd−>fwd, scanning
fwd−>bkwd, scanning
contact
herm. loss
sperm release
n=46
n=8
n=42
n=8
n=66
n=92
n=32
n=74
n=60
n=42
n=30
n=82
n=10
n=9
n=8
Activity
Time (seconds)

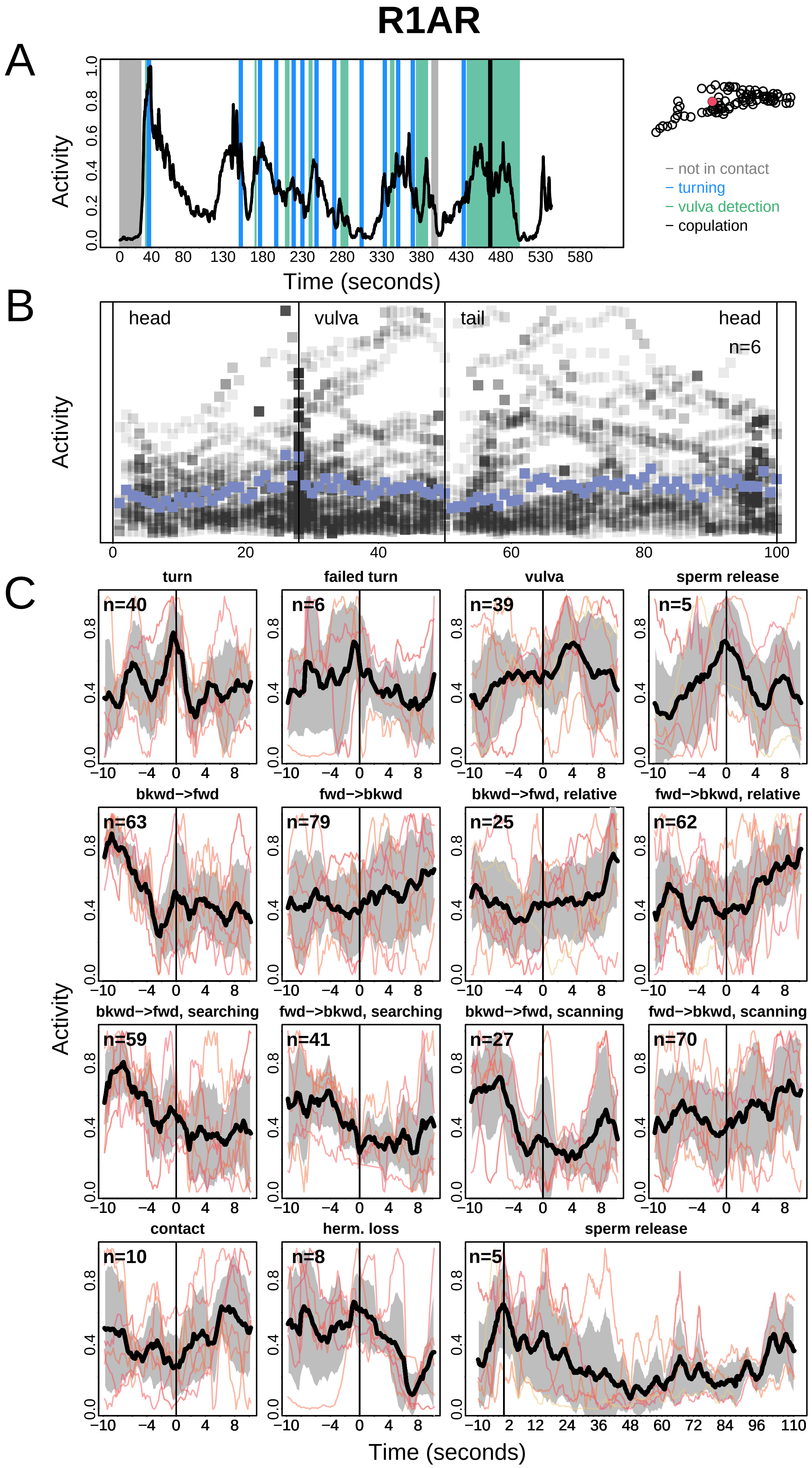
R1AR
A
Activity
Time (seconds)
– not in contact
– turning
– vulva detection
– copulation
B
head
vulva
tail
head
n=6
Activity
C
turn
failed turn
vulva
sperm release
bkwd−>fwd
fwd−>bkwd
bkwd−>fwd, relative
fwd−>bkwd, relative
bkwd−>fwd, searching
fwd−>bkwd, searching
bkwd−>fwd, scanning
fwd−>bkwd, scanning
contact
herm. loss
sperm release
n=40
n=6
n=39
n=5
n=63
n=79
n=25
n=62
n=59
n=41
n=27
n=70
n=10
n=8
n=5
Activity
Time (seconds)

# R1B

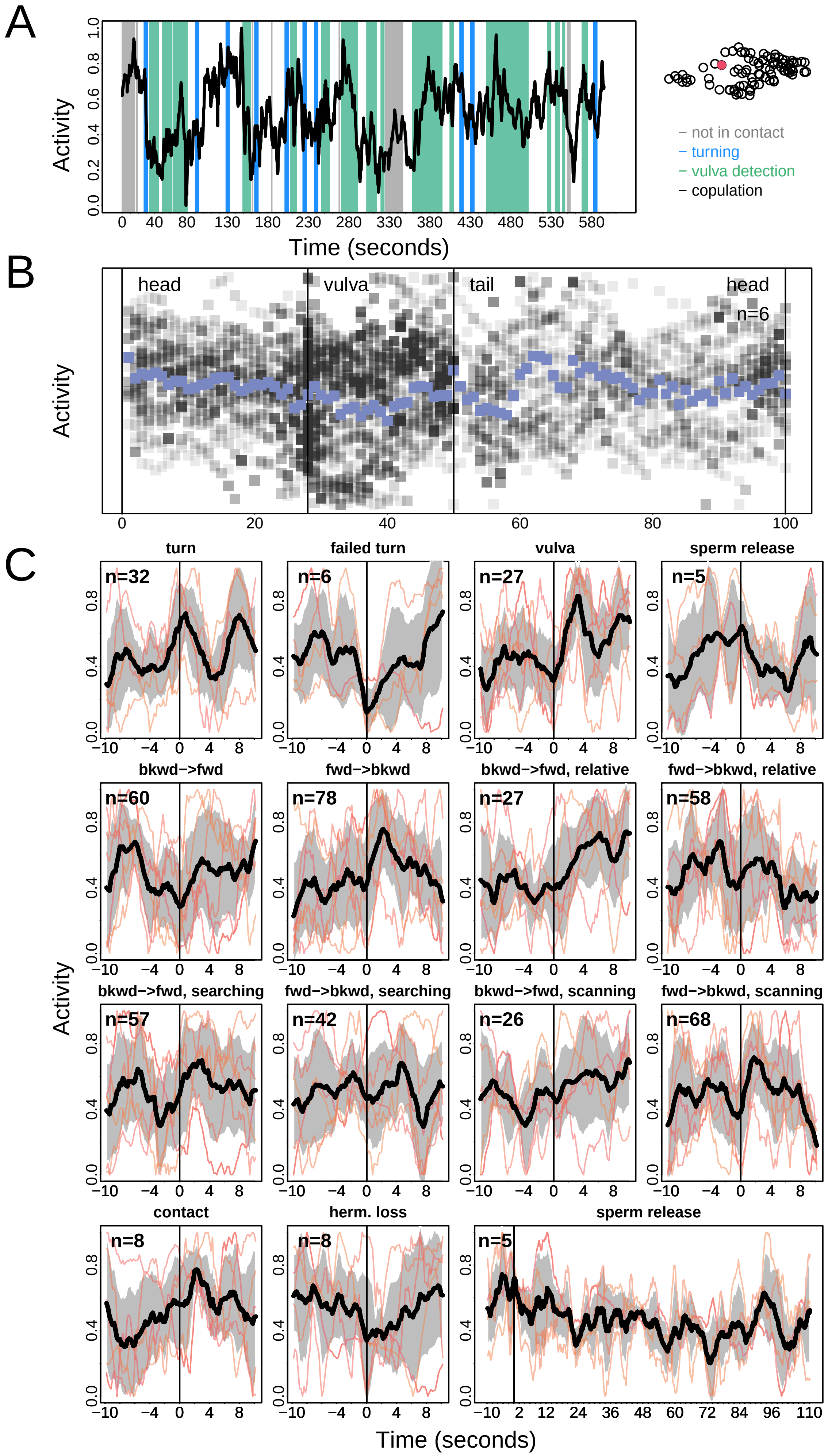

A
Activity
Time (seconds)
– not in contact
– turning
– vulva detection
– copulation
B
head
vulva
tail
head
n=6
Activity
C
turn
n=32
failed turn
n=6
vulva
n=27
sperm release
n=5
bkwd−>fwd
n=60
fwd−>bkwd
n=78
bkwd−>fwd, relative
n=27
fwd−>bkwd, relative
n=58
bkwd−>fwd, searching
n=57
fwd−>bkwd, searching
n=42
bkwd−>fwd, scanning
n=26
fwd−>bkwd, scanning
n=68
contact
n=8
herm. loss
n=8
sperm release
n=5
Activity
Time (seconds)

# R2A

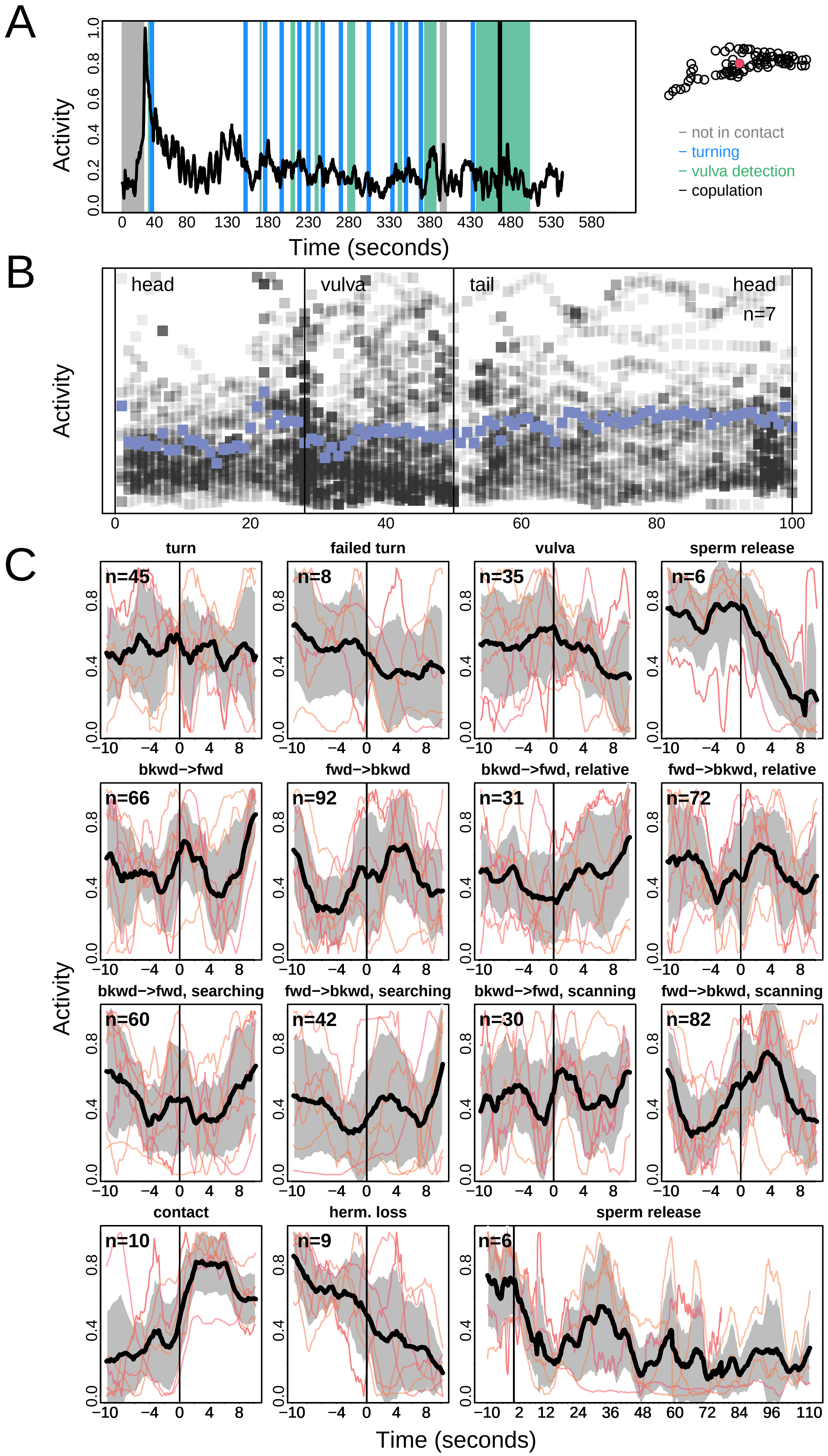
A
Activity
Time (seconds)
not in contact
turning
vulva detection
copulation
B
head
vulva
tail
head
n=7
C
turn
failed turn
vulva
sperm release
bkwd−>fwd
fwd−>bkwd
bkwd−>fwd, relative
fwd−>bkwd, relative
bkwd−>fwd, searching
fwd−>bkwd, searching
bkwd−>fwd, scanning
fwd−>bkwd, scanning
contact
herm. loss
sperm release
n=45
n=8
n=35
n=6
n=66
n=92
n=31
n=72
n=60
n=42
n=30
n=82
n=10
n=9
n=6

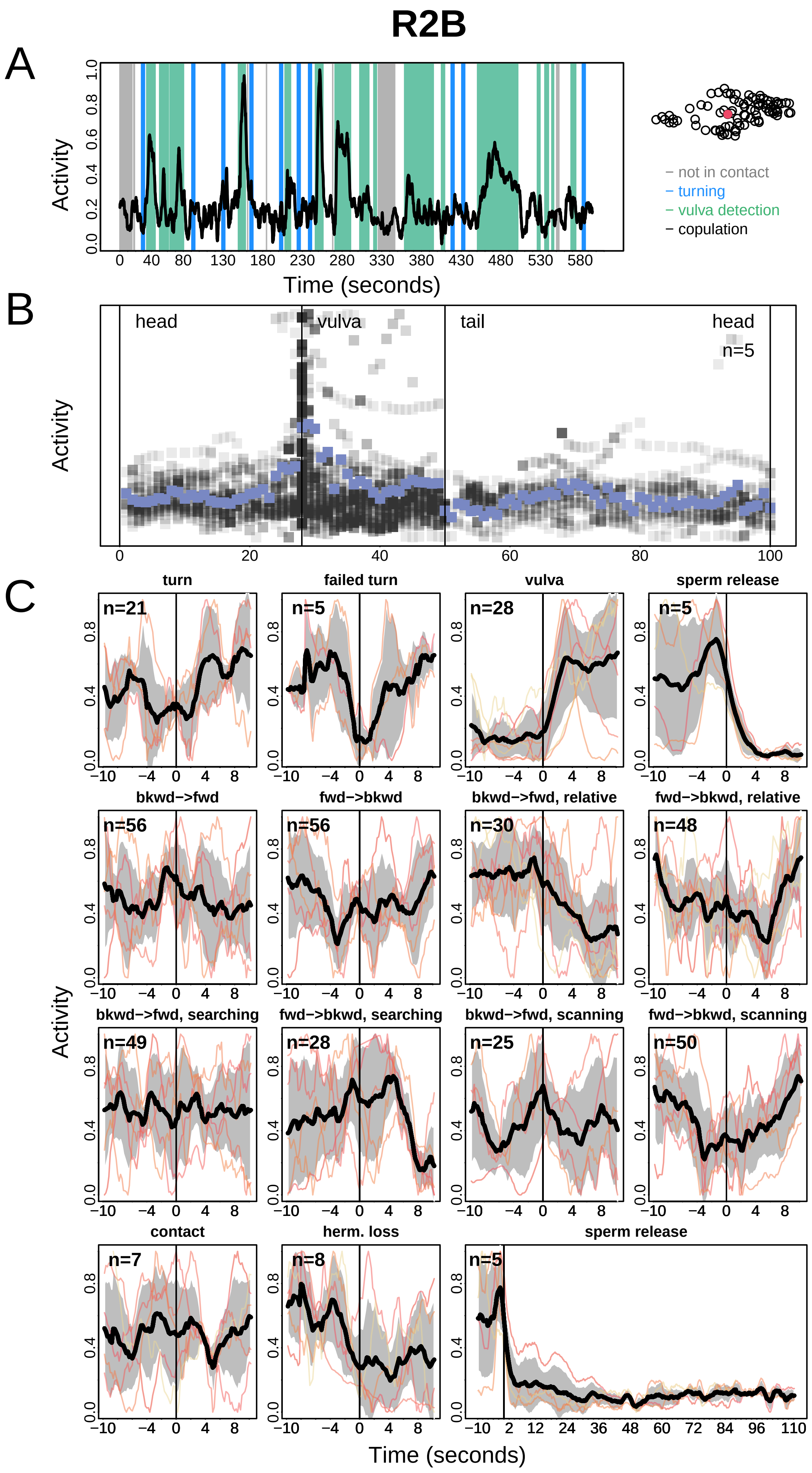
R2B
A
Activity
Time (seconds)
– not in contact
– turning
– vulva detection
– copulation
B
head
vulva
tail
head
n=5
C
turn
failed turn
vulva
sperm release
bkwd−>fwd
fwd−>bkwd
bkwd−>fwd, relative
fwd−>bkwd, relative
bkwd−>fwd, searching
fwd−>bkwd, searching
bkwd−>fwd, scanning
fwd−>bkwd, scanning
contact
herm. loss
sperm release
n=21
n=5
n=28
n=5
n=56
n=56
n=30
n=48
n=49
n=28
n=25
n=50
n=7
n=8
n=5

# R3A

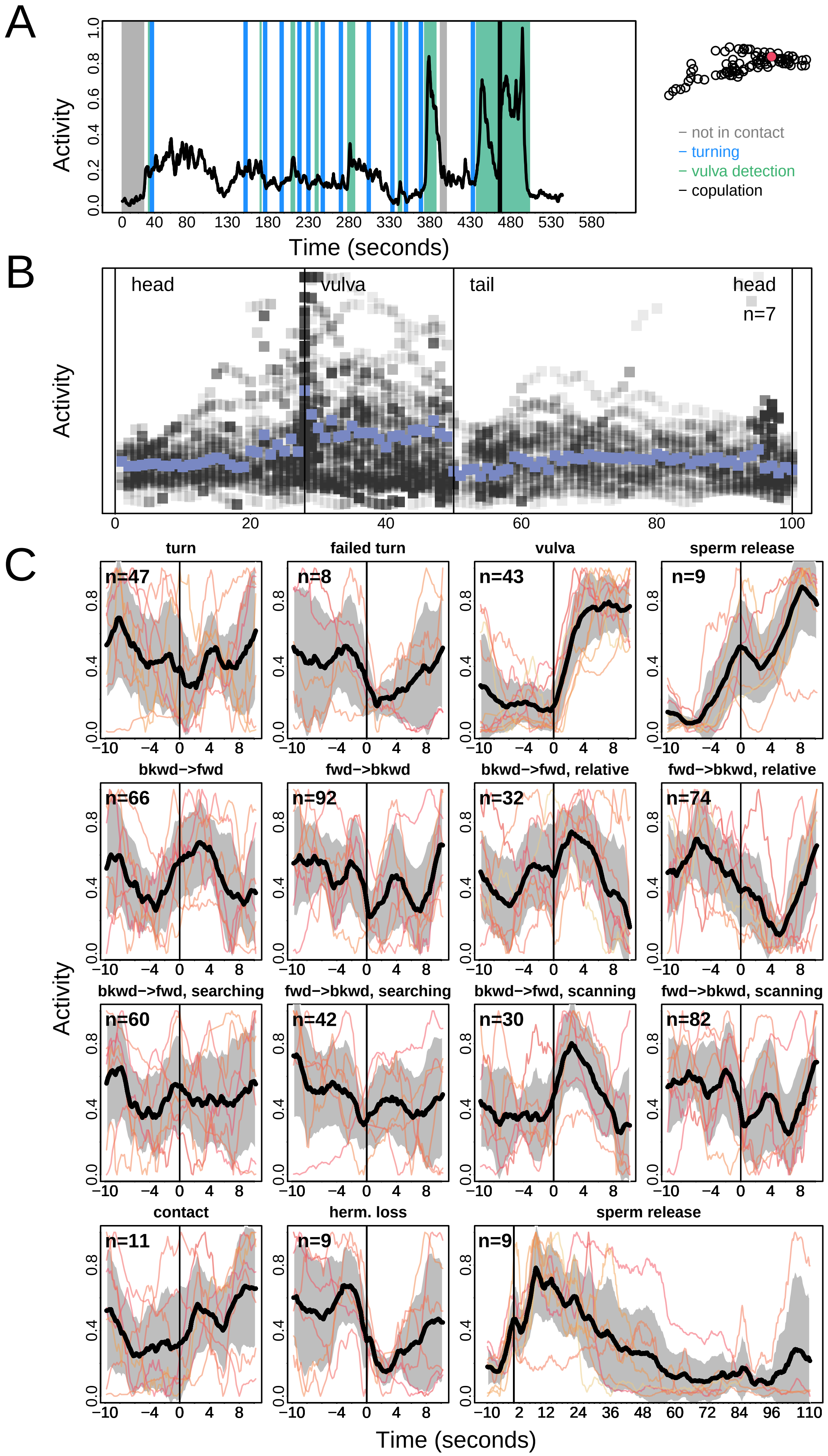
A
Activity
Time (seconds)
0 40 80 130 180 230 280 330 380 430 480 530 580
– not in contact
– turning
– vulva detection
– copulation
B
head
vulva
tail
head
n=7
Activity
0 20 40 60 80 100
C
turn
failed turn
vulva
sperm release
bkwd−>fwd
fwd−>bkwd
bkwd−>fwd, relative
fwd−>bkwd, relative
bkwd−>fwd, searching
fwd−>bkwd, searching
bkwd−>fwd, scanning
fwd−>bkwd, scanning
contact
herm. loss
sperm release
n=47
n=8
n=43
n=9
n=66
n=92
n=32
n=74
n=60
n=42
n=30
n=82
n=11
n=9
n=9
Activity
−10 −4 0 4 8
−10 2 12 24 36 48 60 72 84 96 110
Time (seconds)

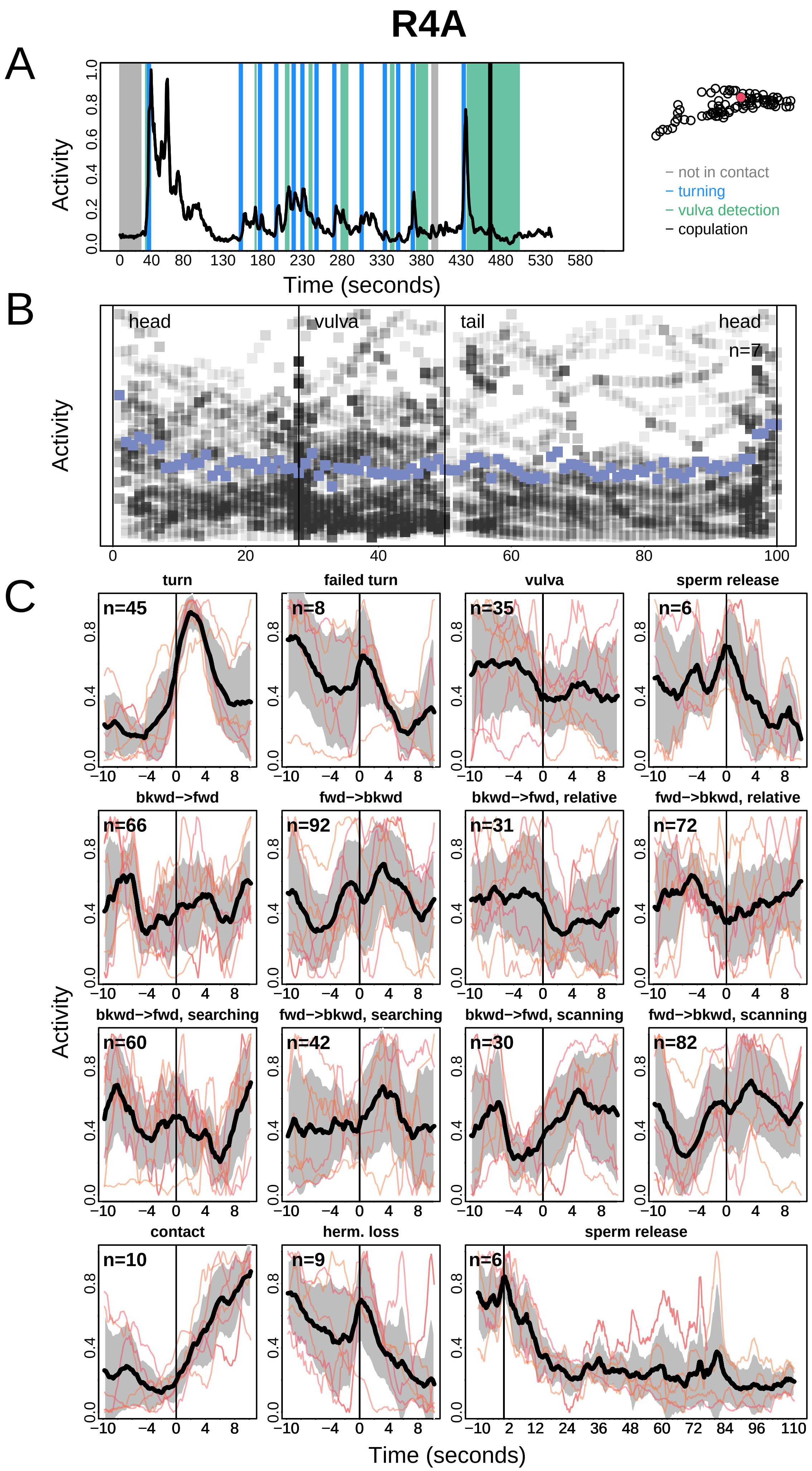
R4A
A
Activity
Time (seconds)
not in contact
turning
vulva detection
copulation
B
head
vulva
tail
head
n=7
Activity
C
turn
failed turn
vulva
sperm release
bkwd−>fwd
fwd−>bkwd
bkwd−>fwd, relative
fwd−>bkwd, relative
bkwd−>fwd, searching
fwd−>bkwd, searching
bkwd−>fwd, scanning
fwd−>bkwd, scanning
contact
herm. loss
sperm release
n=45
n=8
n=35
n=6
n=66
n=92
n=31
n=72
n=60
n=42
n=30
n=82
n=10
n=9
n=6
Activity
Time (seconds)

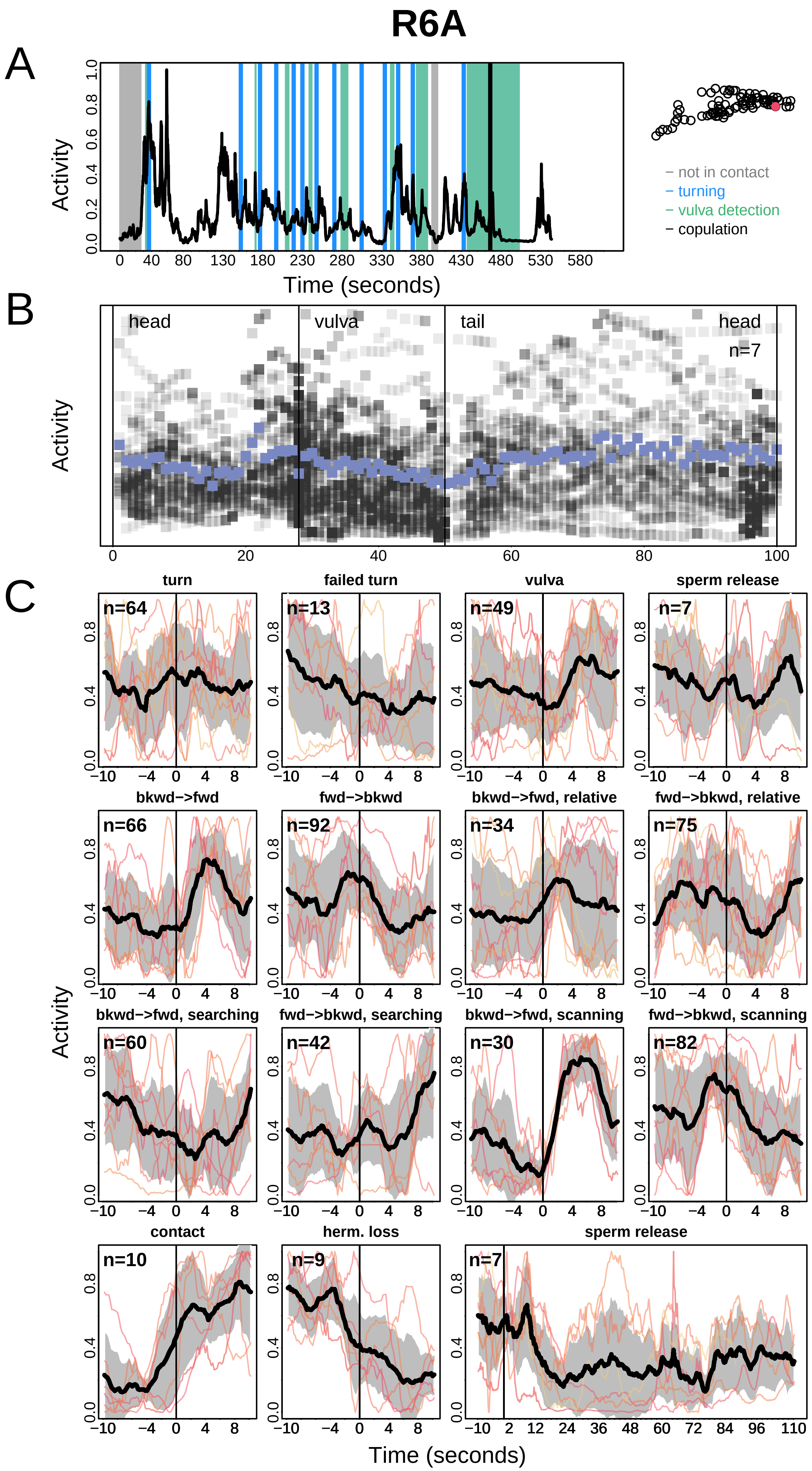

R6A
A
Activity
Time (seconds)
- not in contact
- turning
- vulva detection
- copulation
B
head
vulva
tail
head
n=7
Activity
C
turn
failed turn
vulva
sperm release
bkwd->fwd
fwd->bkwd
bkwd->fwd, relative
fwd->bkwd, relative
bkwd->fwd, searching
fwd->bkwd, searching
bkwd->fwd, scanning
fwd->bkwd, scanning
contact
herm. loss
sperm release
n=64
n=13
n=49
n=7
n=66
n=92
n=34
n=75
n=60
n=42
n=30
n=82
n=10
n=9
n=7
Activity
Time (seconds)

# R8B

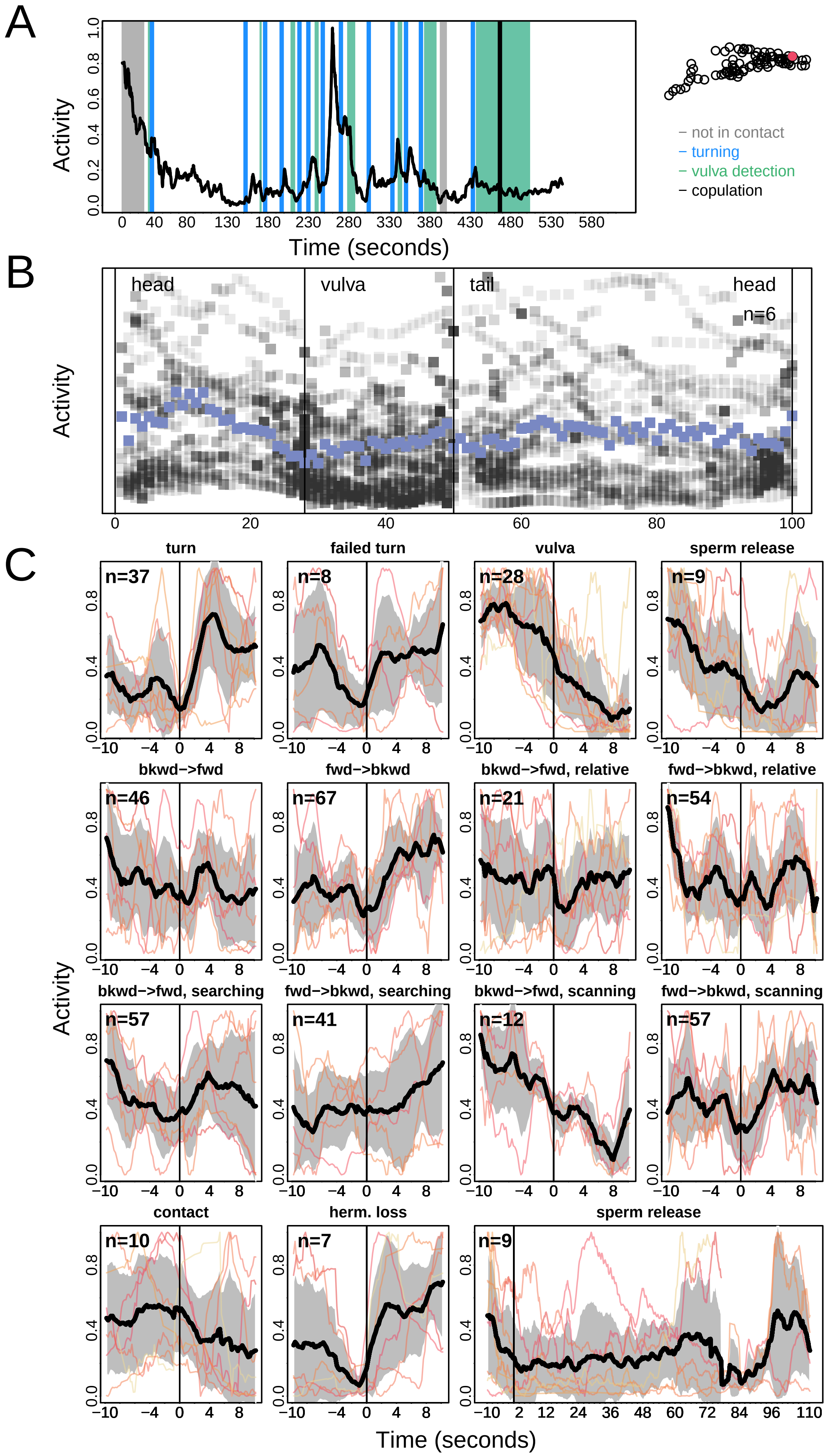
A
Activity
Time (seconds)
– not in contact
– turning
– vulva detection
– copulation
B
head
vulva
tail
head
n=6
Activity
C
turn
failed turn
vulva
sperm release
n=37
n=8
n=28
n=9
bkwd−>fwd
fwd−>bkwd
bkwd−>fwd, relative
fwd−>bkwd, relative
n=46
n=67
n=21
n=54
bkwd−>fwd, searching
fwd−>bkwd, searching
bkwd−>fwd, scanning
fwd−>bkwd, scanning
n=57
n=41
n=12
n=57
contact
herm. loss
sperm release
n=10
n=7
n=9
Activity
Time (seconds)

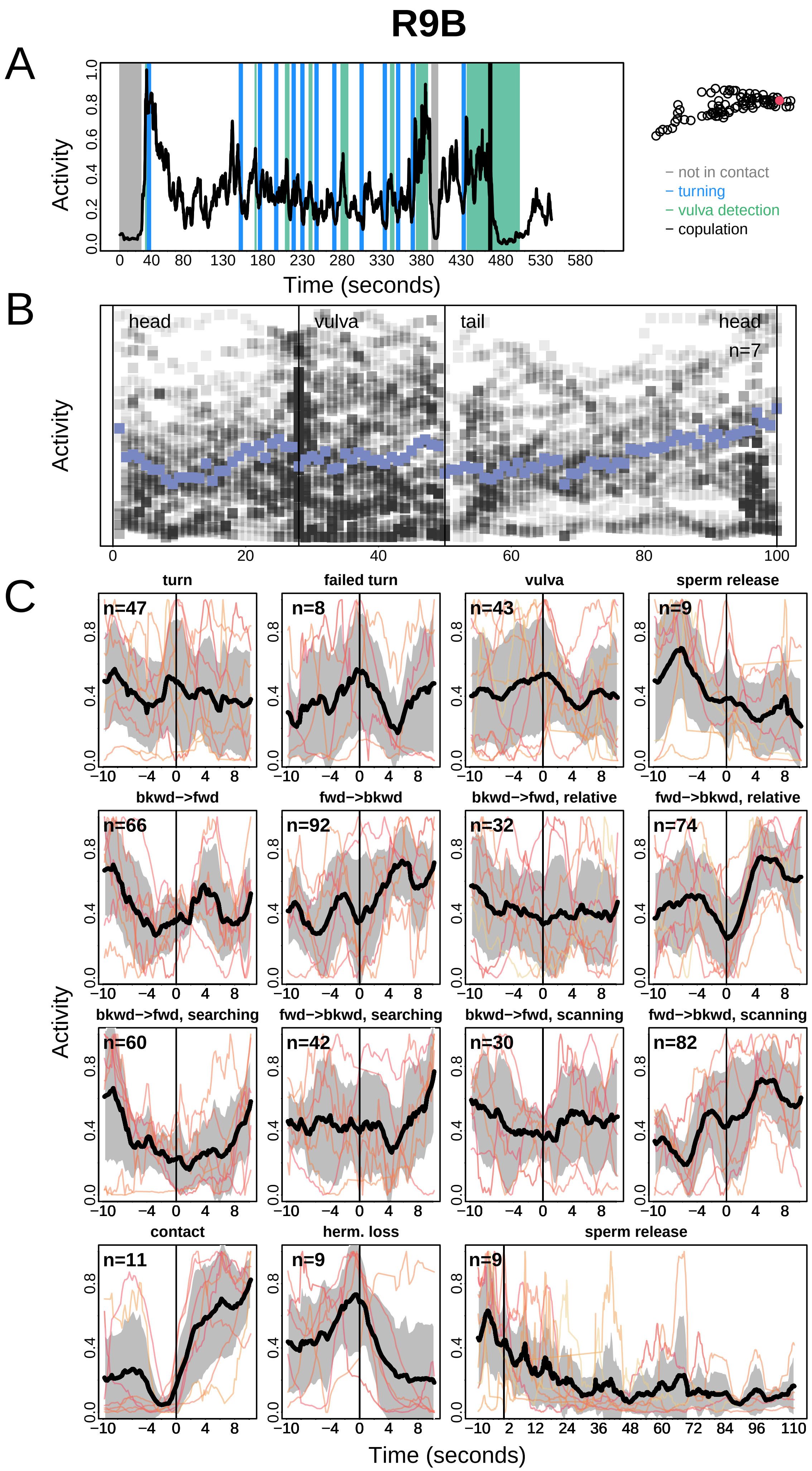
R9B
A
Activity
Time (seconds)
− not in contact
− turning
− vulva detection
− copulation
B
head
vulva
tail
head
n=7
C
turn
failed turn
vulva
sperm release
bkwd−>fwd
fwd−>bkwd
bkwd−>fwd, relative
fwd−>bkwd, relative
bkwd−>fwd, searching
fwd−>bkwd, searching
bkwd−>fwd, scanning
fwd−>bkwd, scanning
contact
herm. loss
sperm release
n=47
n=8
n=43
n=9
n=66
n=92
n=32
n=74
n=60
n=42
n=30
n=82
n=11
n=9
n=9

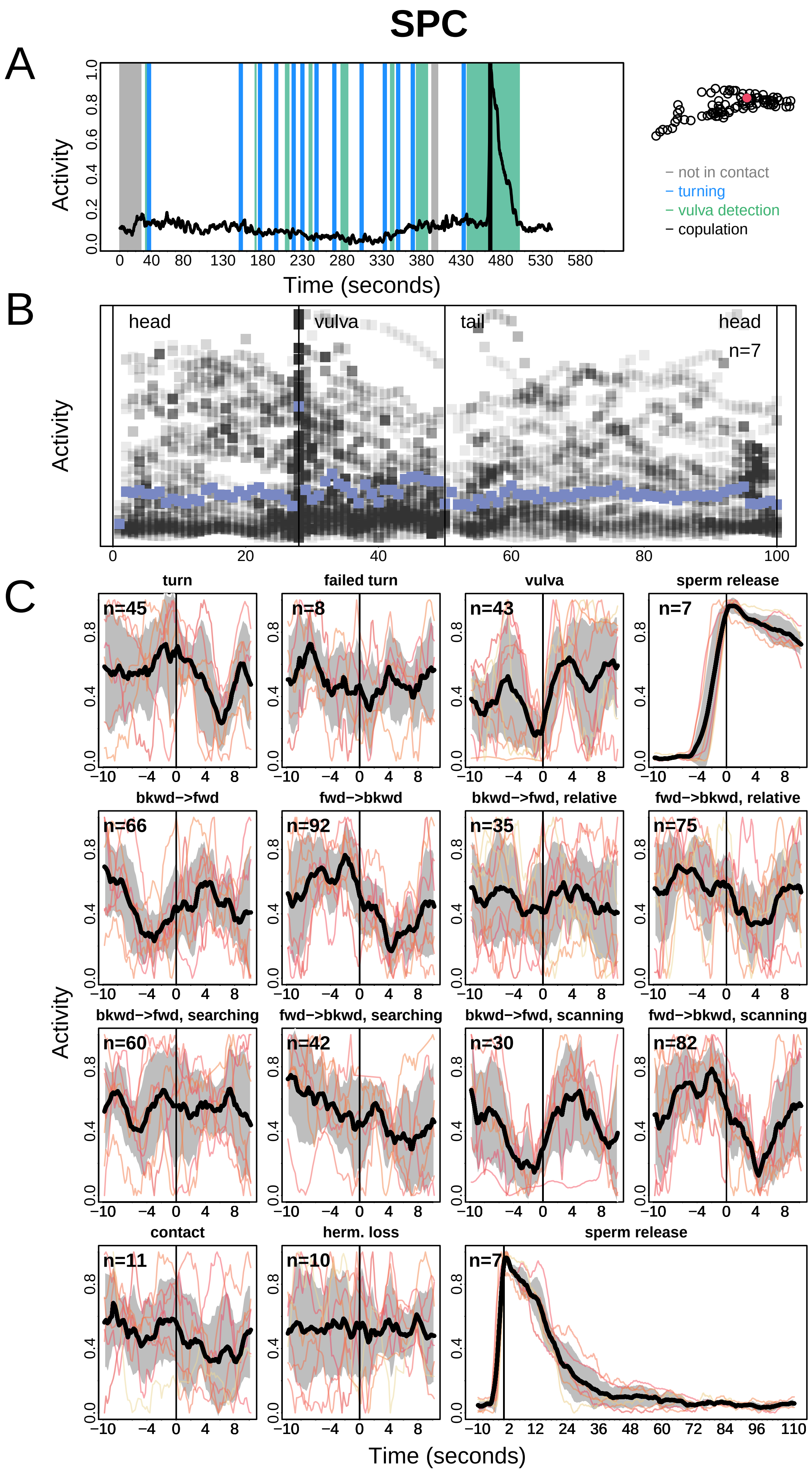
SPC
A
Activity
Time (seconds)
− not in contact
− turning
− vulva detection
− copulation
B
head
vulva
tail
head
n=7
Activity
C
turn
failed turn
vulva
sperm release
n=45
n=8
n=43
n=7
bkwd−>fwd
fwd−>bkwd
bkwd−>fwd, relative
fwd−>bkwd, relative
n=66
n=92
n=35
n=75
bkwd−>fwd, searching
fwd−>bkwd, searching
bkwd−>fwd, scanning
fwd−>bkwd, scanning
n=60
n=42
n=30
n=82
contact
herm. loss
sperm release
n=11
n=10
n=7
Activity
Time (seconds)

### SPD

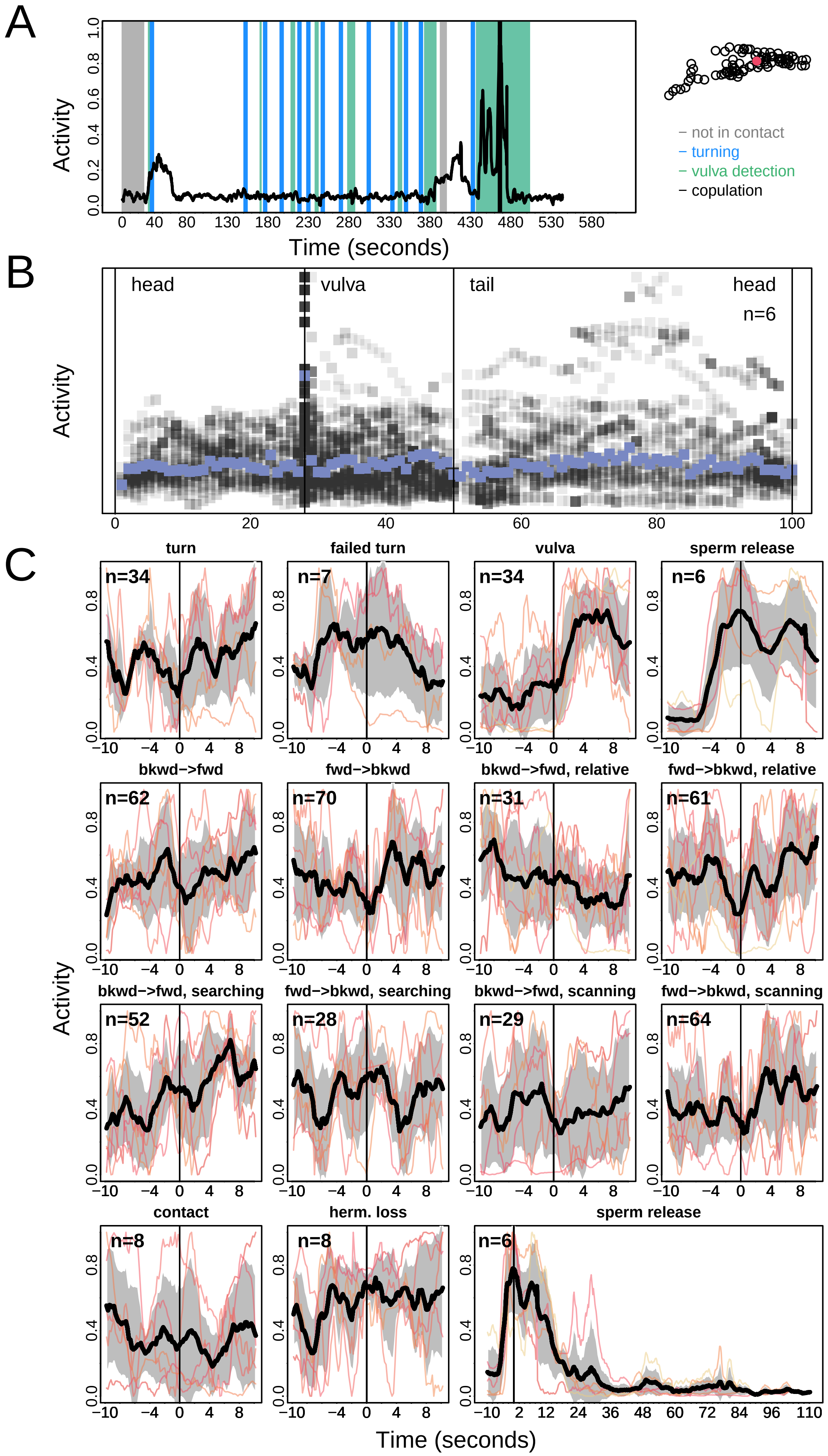
A
Activity
Time (seconds)
not in contact
turning
vulva detection
copulation
B
head
vulva
tail
head
n=6
Activity
C
turn
failed turn
vulva
sperm release
bkwd−>fwd
fwd−>bkwd
bkwd−>fwd, relative
fwd−>bkwd, relative
bkwd−>fwd, searching
fwd−>bkwd, searching
bkwd−>fwd, scanning
fwd−>bkwd, scanning
contact
herm. loss
sperm release
n=34
n=7
n=34
n=6
n=62
n=70
n=31
n=61
n=52
n=28
n=29
n=64
n=8
n=8
n=6
Activity
Time (seconds)

### SPV

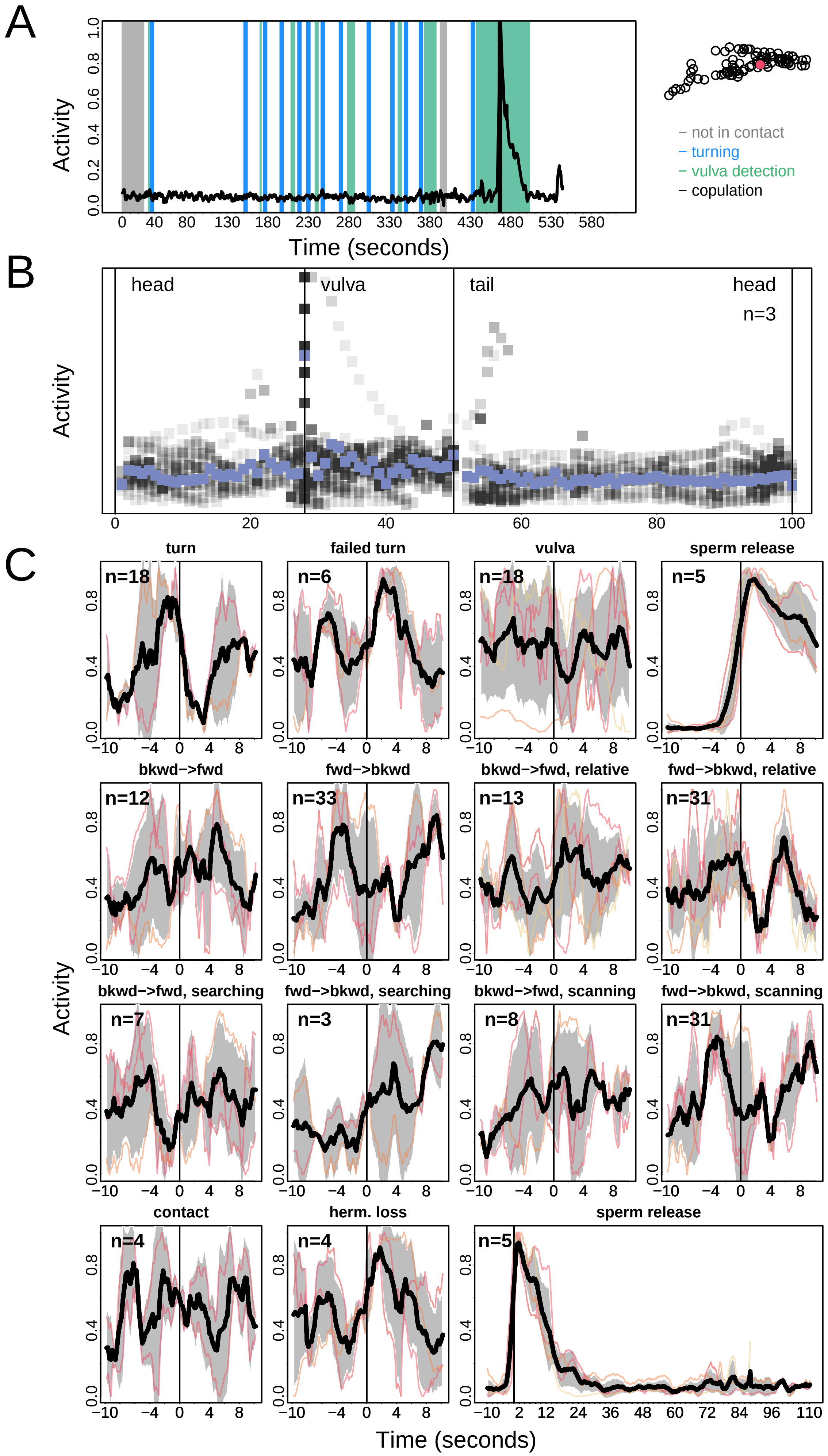
A
Activity
Time (seconds)
– not in contact
– turning
– vulva detection
– copulation
B
head
vulva
tail
head
n=3
Activity
C
turn
failed turn
vulva
sperm release
bkwd−>fwd
fwd−>bkwd
bkwd−>fwd, relative
fwd−>bkwd, relative
bkwd−>fwd, searching
fwd−>bkwd, searching
bkwd−>fwd, scanning
fwd−>bkwd, scanning
contact
herm. loss
sperm release
n=18
n=6
n=18
n=5
n=12
n=33
n=13
n=31
n=7
n=3
n=8
n=31
n=4
n=4
n=5
Activity
Time (seconds)

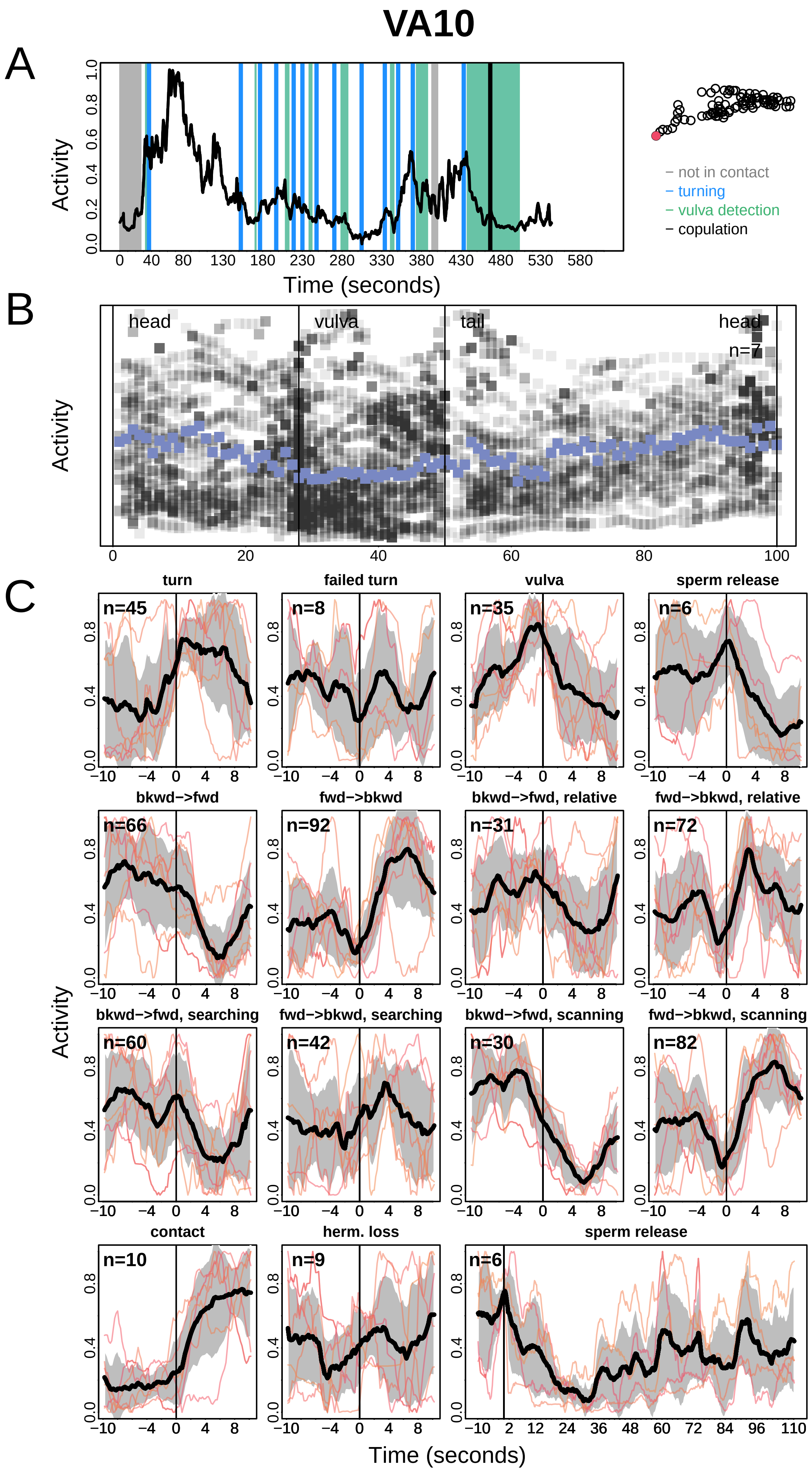

VA10
A
Activity
Time (seconds)
– not in contact
– turning
– vulva detection
– copulation
B
head
vulva
tail
head
n=7
C
turn
failed turn
vulva
sperm release
bkwd−>fwd
fwd−>bkwd
bkwd−>fwd, relative
fwd−>bkwd, relative
bkwd−>fwd, searching
fwd−>bkwd, searching
bkwd−>fwd, scanning
fwd−>bkwd, scanning
contact
herm. loss
sperm release
n=45
n=8
n=35
n=6
n=66
n=92
n=31
n=72
n=60
n=42
n=30
n=82
n=10
n=9
n=6

# VA11

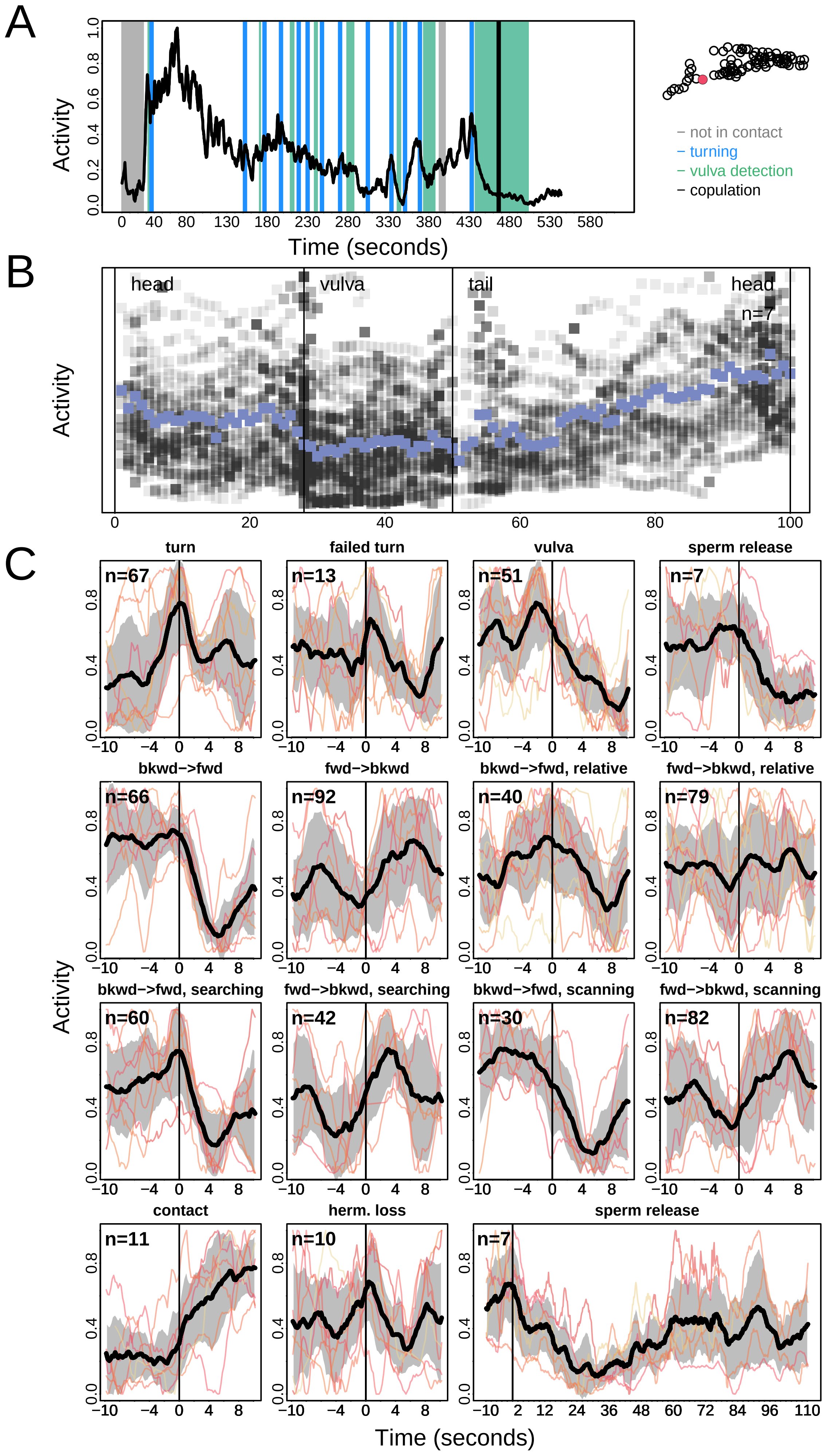
A
Activity
Time (seconds)
0 40 80 130 180 230 280 330 380 430 480 530 580
0.0 0.2 0.4 0.6 0.8 1.0
− not in contact
− turning
− vulva detection
− copulation
B
head
vulva
tail
head
n=7
0 20 40 60 80 100
C
turn
n=67
failed turn
n=13
vulva
n=51
sperm release
n=7
bkwd−>fwd
n=66
fwd−>bkwd
n=92
bkwd−>fwd, relative
n=40
fwd−>bkwd, relative
n=79
bkwd−>fwd, searching
n=60
fwd−>bkwd, searching
n=42
bkwd−>fwd, scanning
n=30
fwd−>bkwd, scanning
n=82
contact
n=11
herm. loss
n=10
sperm release
n=7
−10 −4 0 4 8
−10 2 12 24 36 48 60 72 84 96 110
0.0 0.4 0.8
Time (seconds)

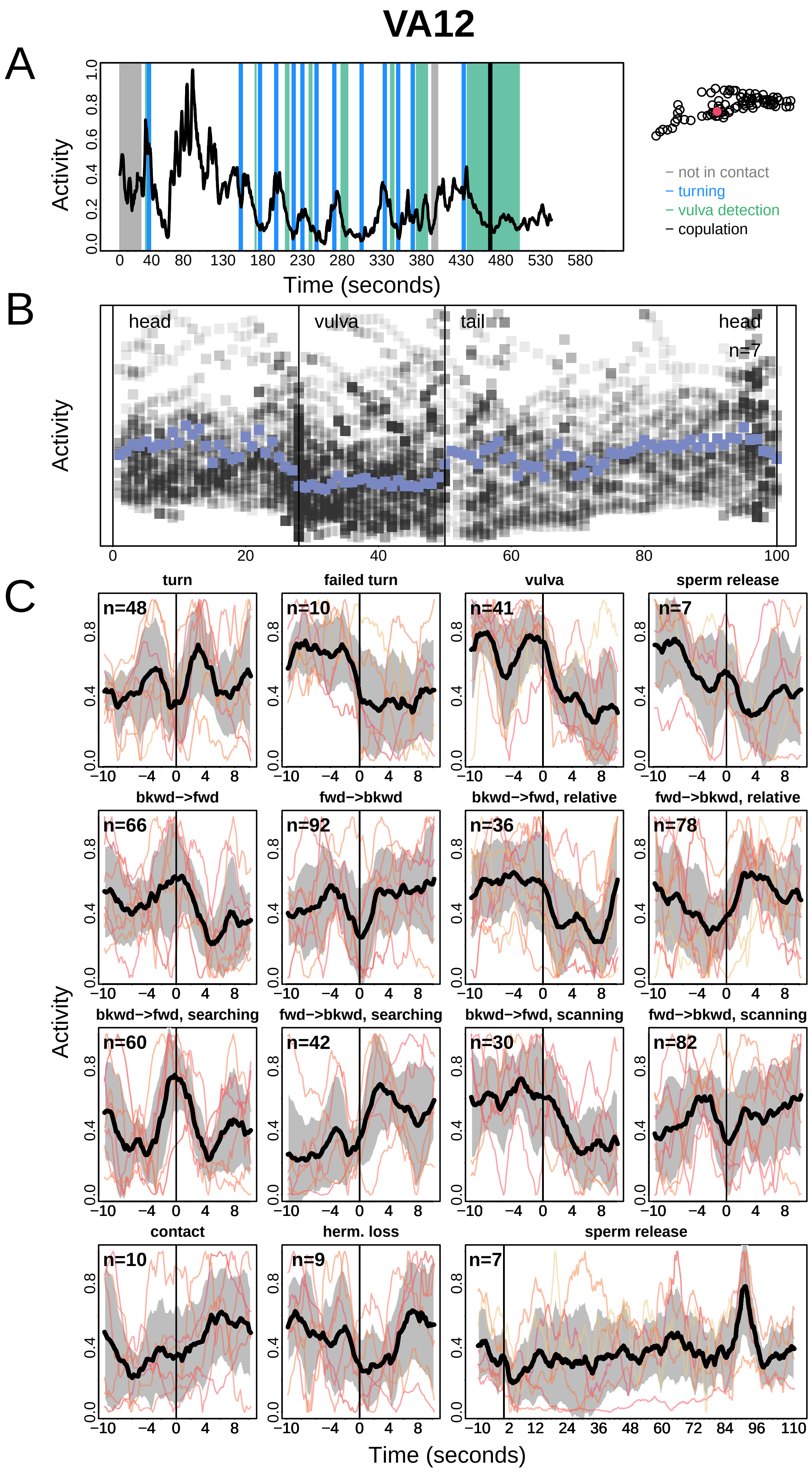

VA12
A
B
C
Activity
Time (seconds)
− not in contact
− turning
− vulva detection
− copulation
head
vulva
tail
head
n=7
turn
failed turn
vulva
sperm release
bkwd−>fwd
fwd−>bkwd
bkwd−>fwd, relative
fwd−>bkwd, relative
bkwd−>fwd, searching
fwd−>bkwd, searching
bkwd−>fwd, scanning
fwd−>bkwd, scanning
contact
herm. loss
sperm release
n=48
n=10
n=41
n=7
n=66
n=92
n=36
n=78
n=60
n=42
n=30
n=82
n=10
n=9
n=7

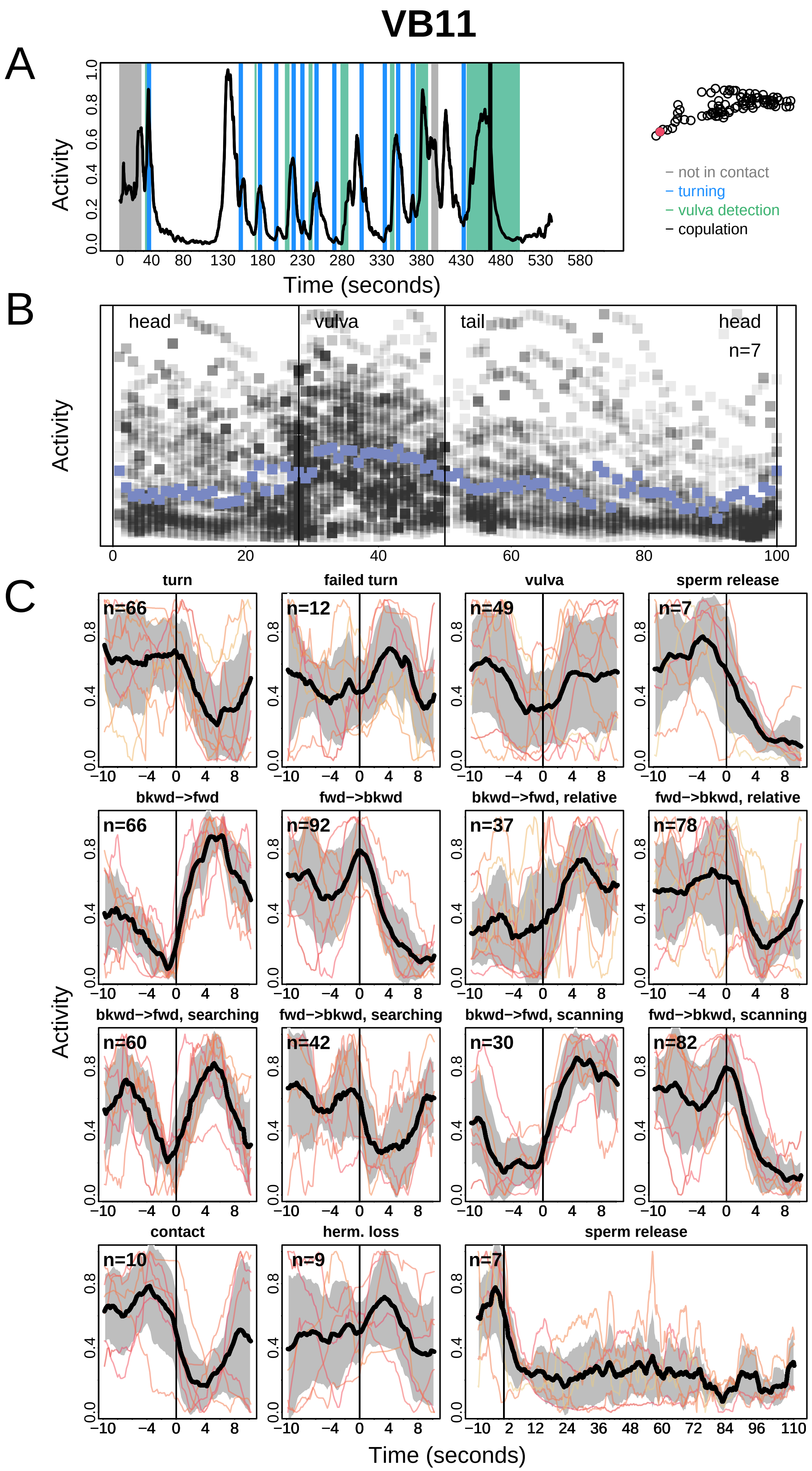
VB11
A
Activity
Time (seconds)
– not in contact
– turning
– vulva detection
– copulation
B
head
vulva
tail
head
n=7
Activity
C
turn
failed turn
vulva
sperm release
bkwd–>fwd
fwd–>bkwd
bkwd–>fwd, relative
fwd–>bkwd, relative
bkwd–>fwd, searching
fwd–>bkwd, searching
bkwd–>fwd, scanning
fwd–>bkwd, scanning
contact
herm. loss
sperm release
n=66
n=12
n=49
n=7
n=66
n=92
n=37
n=78
n=60
n=42
n=30
n=82
n=10
n=9
n=7
Activity
Time (seconds)

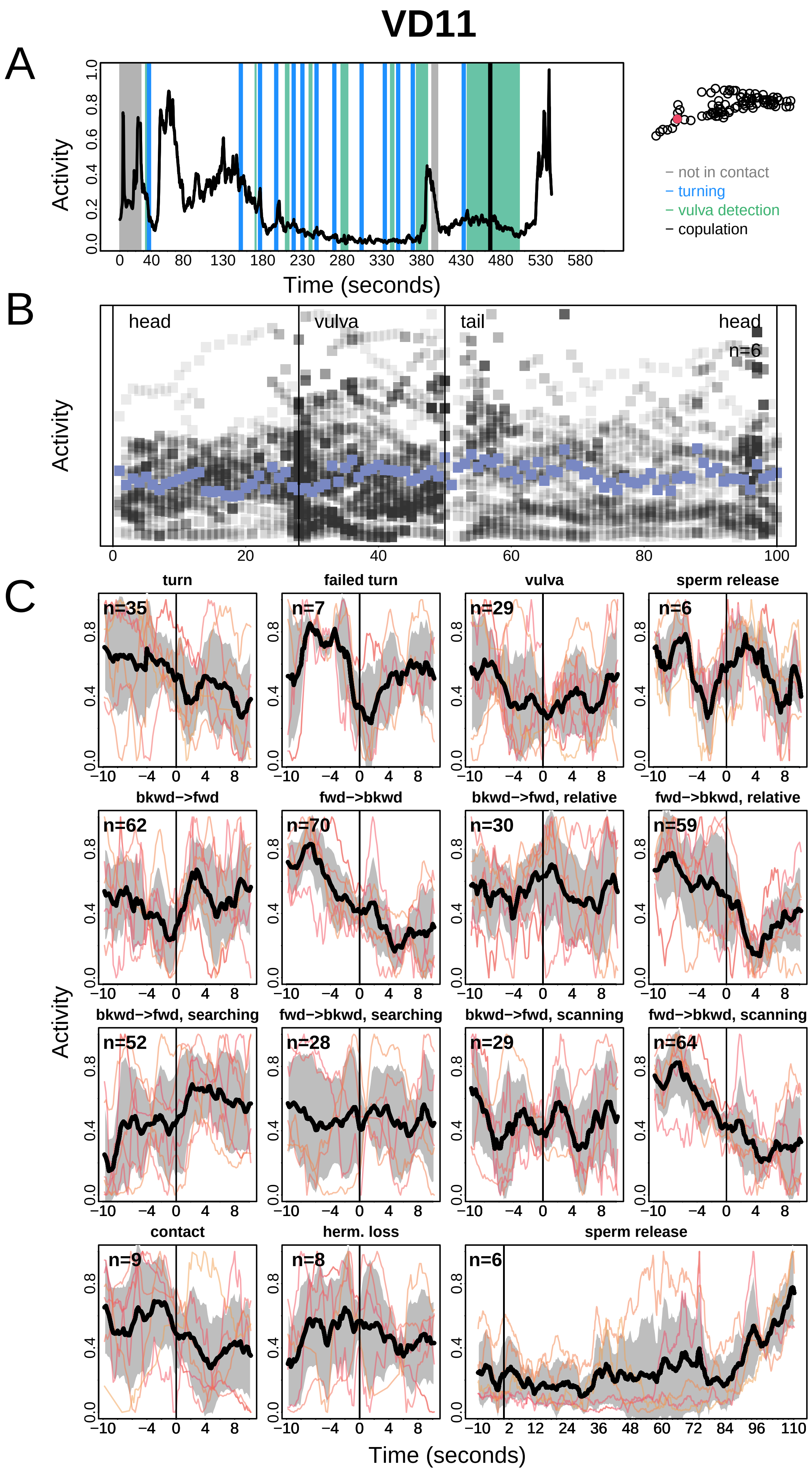
VD11
A
Activity
Time (seconds)
0 40 80 130 180 230 280 330 380 430 480 530 580
0.0 0.2 0.4 0.6 0.8 1.0
− not in contact
− turning
− vulva detection
− copulation
B
Activity
head
vulva
tail
head
n=6
0 20 40 60 80 100
C
turn
n=35
failed turn
n=7
vulva
n=29
sperm release
n=6
bkwd−>fwd
n=62
fwd−>bkwd
n=70
bkwd−>fwd, relative
n=30
fwd−>bkwd, relative
n=59
bkwd−>fwd, searching
n=52
fwd−>bkwd, searching
n=28
bkwd−>fwd, scanning
n=29
fwd−>bkwd, scanning
n=64
contact
n=9
herm. loss
n=8
sperm release
n=6
−10 −4 0 4 8
−10 2 12 24 36 48 60 72 84 96 110
0.0 0.4 0.8
Activity
Time (seconds)

# VD12

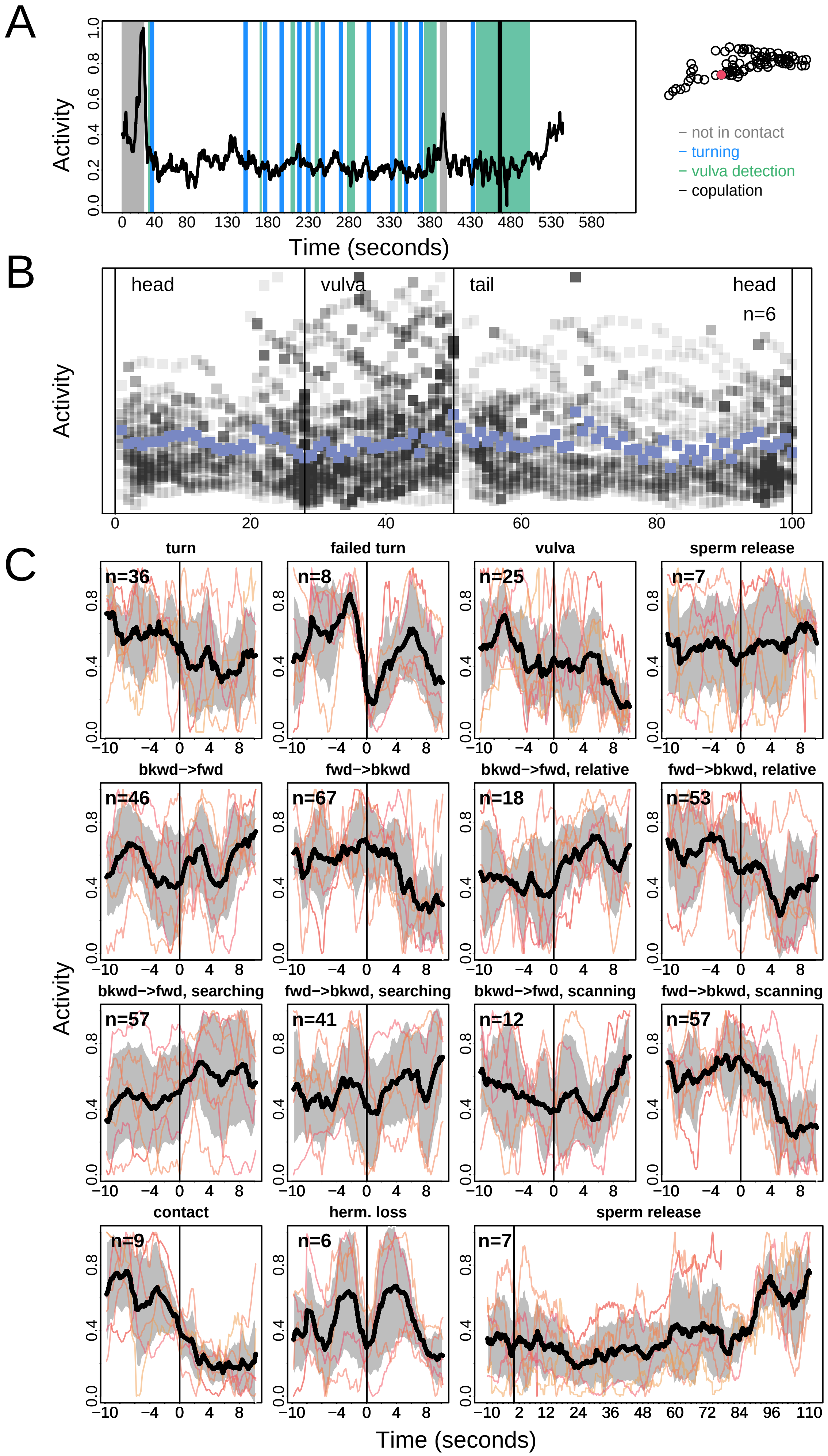
A
B
C
Activity
Time (seconds)
not in contact
turning
vulva detection
copulation
head
vulva
tail
head
n=6
turn
failed turn
vulva
sperm release
bkwd−>fwd
fwd−>bkwd
bkwd−>fwd, relative
fwd−>bkwd, relative
bkwd−>fwd, searching
fwd−>bkwd, searching
bkwd−>fwd, scanning
fwd−>bkwd, scanning
contact
herm. loss
sperm release
n=36
n=8
n=25
n=7
n=46
n=67
n=18
n=53
n=57
n=41
n=12
n=57
n=9
n=6
n=7

# VD13

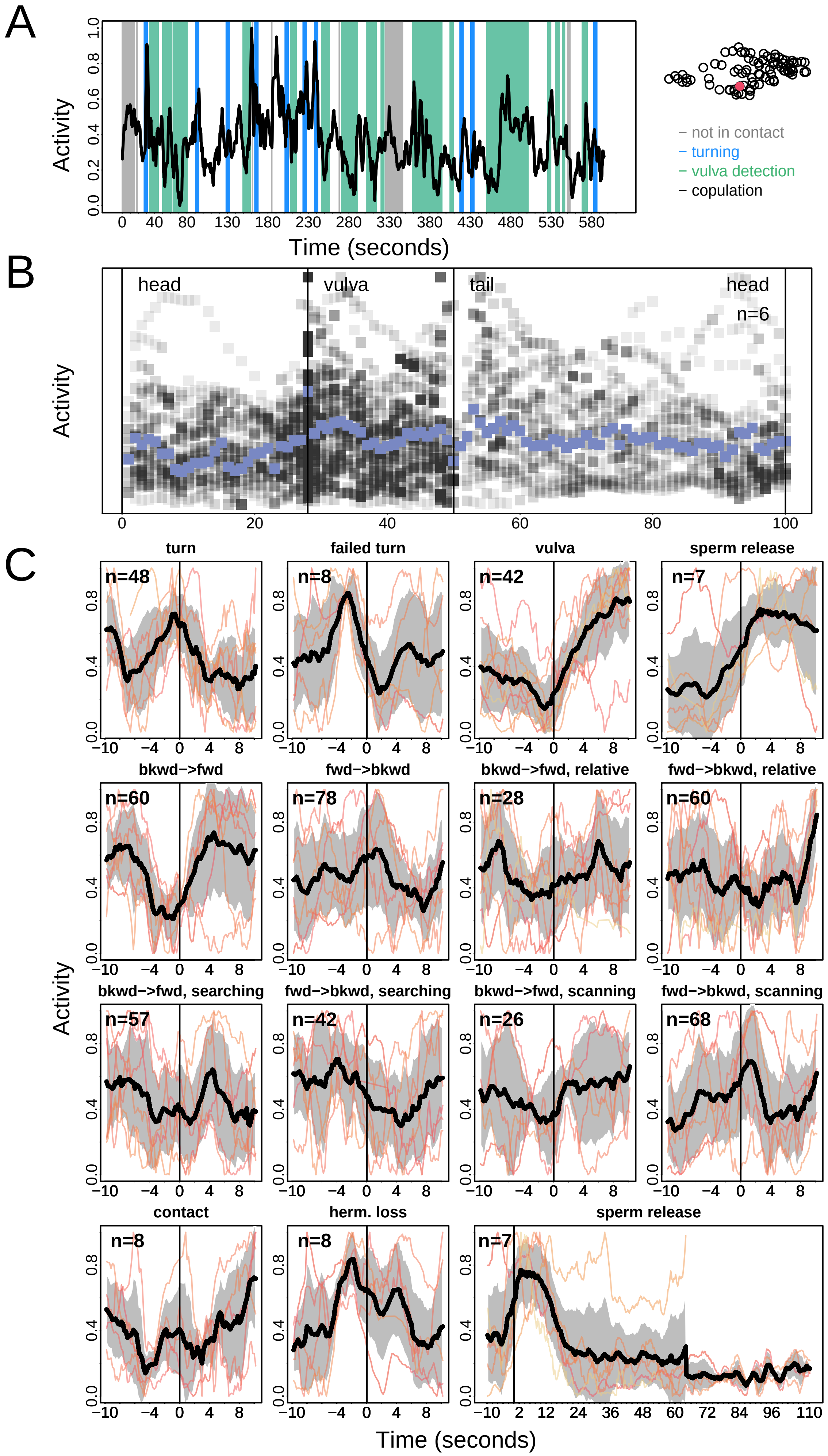

A
B
C
Activity
Time (seconds)
not in contact
turning
vulva detection
copulation
head
vulva
tail
head
n=6
turn
failed turn
vulva
sperm release
bkwd−>fwd
fwd−>bkwd
bkwd−>fwd, relative
fwd−>bkwd, relative
bkwd−>fwd, searching
fwd−>bkwd, searching
bkwd−>fwd, scanning
fwd−>bkwd, scanning
contact
herm. loss
sperm release
n=48
n=8
n=42
n=7
n=60
n=78
n=28
n=60
n=57
n=42
n=26
n=68
n=8
n=8
n=7
